# A multi-tissue study of immune gene expression profiling highlights the key role of the nasal epithelium in COVID-19 severity

**DOI:** 10.1101/2021.10.27.466206

**Authors:** Alberto Gómez-Carballa, Irene Rivero-Calle, Jacobo Pardo-Seco, José Gómez-Rial, Carmen Rivero-Velasco, Nuria Rodríguez-Núñez, Gema Barbeito-Castiñeiras, Hugo Pérez-Freixo, Miriam Cebey-López, Ruth Barral-Arca, Carmen Rodriguez-Tenreiro, Ana Dacosta-Urbieta, Xabier Bello, Sara Pischedda, María José Currás-Tuala, Sandra Viz-Lasheras, Federico Martinón-Torres, Antonio Salas, GEN-COVID (www.gencovid.eu) study group

## Abstract

**Background:** COVID-19 symptoms range from mild to severe illness; the cause for this differential response to infection remains unknown. Unravelling the immune mechanisms acting at different levels of the colonization process might be key to understand these differences.

**Methods and findings:** We carried out a multi-tissue (nasal, buccal and blood; *n* = 156) gene expression analysis of immune-related genes from patients affected by different COVID-19 severities, and healthy controls through the nCounter technology. We then used a differential expression approach and pathways analysis to detect tissue specific immune severity signals in COVID-19 patients.

Mild and asymptomatic cases showed a powerful innate antiviral response in nasal epithelium, characterized by activation of interferon (IFN) pathway and downstream cascades, successfully controlling the infection at local level. In contrast, weak macrophage/monocyte driven innate antiviral response and lack of IFN signalling activity were shown in severe cases. Consequently, oral mucosa from severe patients showed signals of viral activity, cell arresting and viral dissemination to the lower respiratory tract, which ultimately could explain the exacerbated innate immune response and impaired adaptative immune responses observed at systemic level. Results from saliva transcriptome suggest that the buccal cavity might play a key role in SARS-CoV-2 infection and dissemination in patients with worse prognosis.

**Conclusions:** We found severity-related signatures in patient tissues mainly represented by genes involved in the innate immune system and cytokine/chemokine signalling. Local immune response could be key to determine the course of the systemic response and thus COVID-19 severity. Our findings provide a framework to investigate severity host gene biomarkers and pathways that might be relevant to diagnosis, prognosis, and therapy.

## Introduction

The severe acute respiratory syndrome coronavirus 2 (SARS-CoV-2), which causes the coronavirus disease 19 (COVID-19), was first identified in patients suffering from a respiratory disease in Wuhan (Hubei province, China) in December 2019 (1, 2). Soon after, SARS-CoV-2 quickly spread across multiple countries, reaching high frequencies in some geographic regions favored by two main catalyzers, namely variants showing higher transmissibility (3) and superspreading events (4–6). To date, the SARS-CoV-2 pandemic has caused >3.8M deaths and >175M confirmed cases worldwide (www.who.int; 15-06-2021). SARS-CoV-2 is primarily transmitted by airway droplets; the nasal epithelium has been postulated as the main portal of entrance and transmission of the virus, and the location where higher viral loads and replication rates were observed (7–9).

COVID-19 is mainly characterized by producing influenza-like symptoms, and other (e.g., gastro-intestinal) non-related complications (10, 11). A wide range of symptoms have been described, from absolutely no symptoms to severe pneumonia, lung damage, and progressive respiratory failure (12).Investigating the immunological processes underpinning SARS-CoV-2 pathogenesis is key for the development of efficient therapeutic strategies and vaccines against COVID-19. The immune system dysregulation described in severe patients is characterized by a reduce number of lymphocyte T subpopulations CD4+ and CD8+, pointing to an absence of T-cell specific response (13–15). Impairment of the adaptative immune system, together with a non-effective host immune response by innate cells, could be the cause of a higher viral replication and the activation of inflammatory processes, resulting in tissue damage and a more severe disease course (13, 16). The hyper-inflammatory response occurring during the acute phase is mainly driven by the monocyte-macrophage activation (17), the release of pro-inflammatory cytokines (13, 18, 19) and increase of other mediators e.g. ferritin and C-reactive protein (CRP). The infiltration of monocytes-derived macrophages from blood points to these cells as the main cause of the lung tissue damage in severe cases (16).

Analysis of blood gene expression of immune-related genes has allowed to shed light on the immune response to pathogen infection. In COVID-19, results from transcriptomic studies conducted on peripheral blood mononuclear cells (PBMCs) indicate that genes associated with inflammatory and hypercoagulability pathways (20), and the imbalance between innate and adaptive immune responses, are the main factors responsible for severe disease course (21). Immune gene expression analysis carried out in serial whole blood from three patients with mild, moderate, and severe disease (22) showed a reduce adaptative response in severe cases, and an attenuated T-cell response in mild disease probably related to a prolonged infection. Thair et al. (23) found a differential transcriptomic pattern in whole blood transcriptomes that differentiated COVID-19 from other viral infections, suggesting that their top differentially expressed genes (DEGs) may be linked to new mechanisms of pathogen evasion from host immune response. Most recently, Ashenbrenner et al. (24) revealed neutrophil activation signatures in severe cases, coupled with an elevated expression of genes related to coagulation and platelet function, coupled with an absence of T-cell activation. From a non-systemic perspective, most of the studies were carried out *in vitro* or nasal-derived epithelial cells (7, 25–27). Only a few studies were performed on naso-pharyngeal (NP) swabs of infected patients; the results point to different expression patterns depending on the viral load, age, and sex (28). Comparison of expression profiles of infected adults and children (29) indicates an early and more efficient innate immune response in nasal mucosa from children, thus conditioning the final outcome of the disease. So far, only one study undertaken in three severe patients has proposed candidate transcripts for severity and disease outcome in NP samples (30). Saliva has also been proposed as a good alternative to NP swabs for SARS-CoV-2 detection through a non-invasive approach (31–33), since it is easier to collect and yield comparable results (34). The study of the specific host gene expression profile produced by the SARS-CoV-2 infection in oral mucosa has not raised comparable interest; there are only a few *in-silico* predictive studies using retrospective available data from non-infected subjects (35, 36), and a recent study (37) highlighting the importance of the oral cavity in the viral transmission and dissemination.

We carried out a multi-source gene expression study at three different transcriptomic layers, using blood, nasal and saliva samples, collected on asymptomatic, mild, and severe patients. Our study aims at understanding the differential immune response that leads to different disease severities, and finding specific biomarkers of severity to improve patient management and prognostic prediction.

## Methods

### Patient recruitment

We prospectively collected blood, saliva, and nasal epithelium samples from 52 COVID-19 patients with confirmed PCR-positive diagnosis of SARS-CoV-2 infection in the Hospital Clínico Universitario of Santiago de Compostela (Galicia; Spain) from March to June 2020. Patients and healthy controls were self-reported as of South-European ancestry; and cases were classified at the time of sample collection by severity illness: severe (ICU admission), moderate (non-ICU but admitted to hospital) and mild (domiciliary lockdown patients with mild symptoms or asymptomatic; herein mild patients). Comparisons were carried out as follows: *(i)* whole blood from 41 patients and 13 controls, (*ii*) nasal epithelium from 38 patients and 11 controls, and (*iii*) saliva from 41 patients and 12 controls. The complete tissue sample set could be collected for 27 out of the 52 patients. Demographic and clinical data from the patients are summarized in Table 1 (see **S1 Table** for more details). Healthy controls are all from the same geographic region as the cases.

**Table 1.**
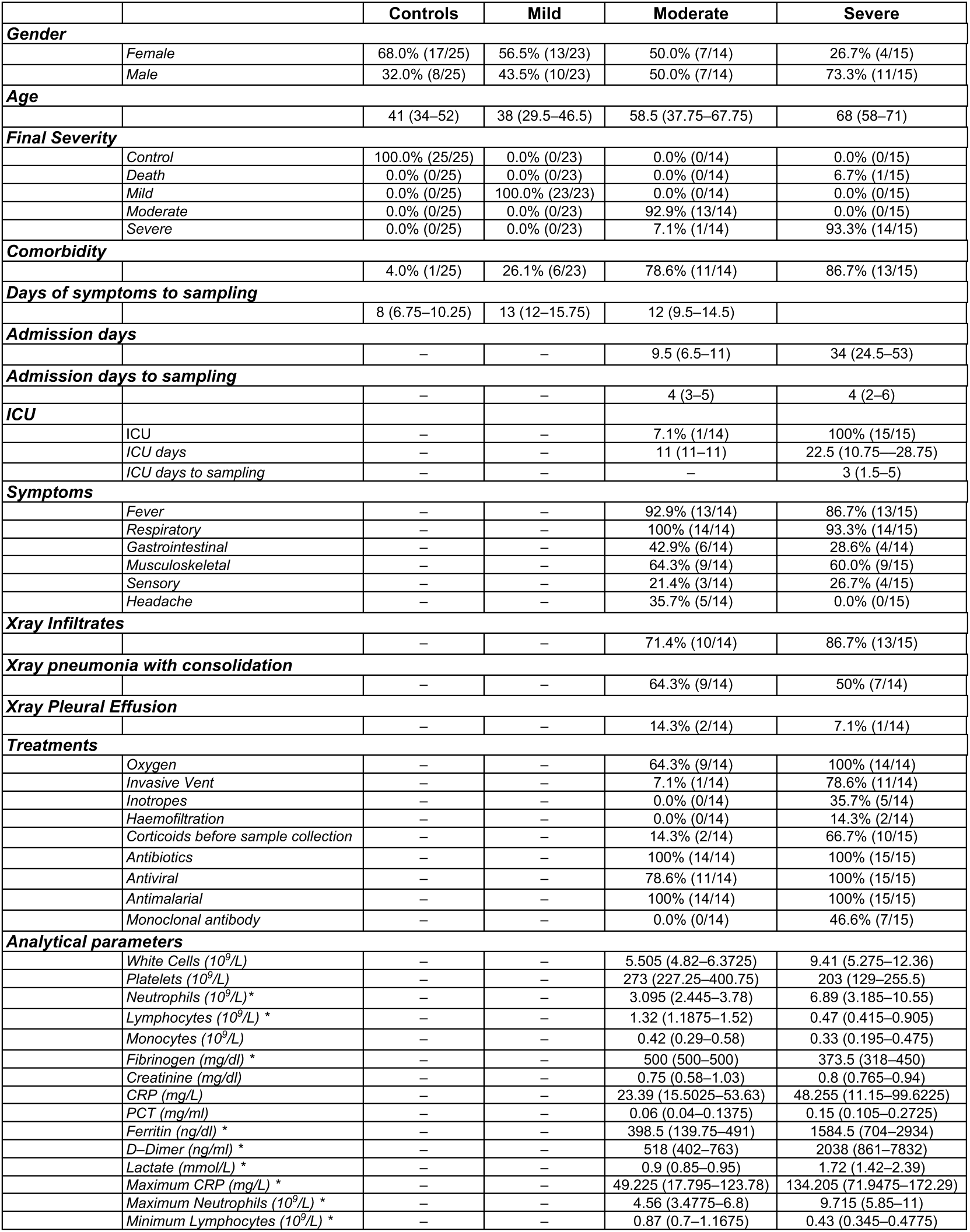
Demographic and clinical characteristics of the COVID-19 cohort and controls. Phenotype stratification of the patients refers to the severity at the time of sample collection. Some of the clinical parameters are only available for hospitalized cases (moderate and severe). Quantitative variables are represented by the median (IQR) and qualitative variables as percentages with absolute numbers in brackets. Asterisks indicate statistical significant differences between groups.

|  |  | Controls | Mild | Moderate | Severe |
| --- | --- | --- | --- | --- | --- |
| **Gender** |  |  |  |  |  |
|  | Female | 68.0% (17/25) | 56.5% (13/23) | 50.0% (7/14) | 26.7% (4/15) |
|  | Male | 32.0% (8/25) | 43.5% (10/23) | 50.0% (7/14) | 73.3% (11/15) |
| **Age** |  |  |  |  |  |
|  |  | 41 (34–52) | 38 (29.5–46.5) | 58.5 (37.75–67.75) | 68 (58–71) |
| **Final Severity** |  |  |  |  |  |
|  | Control | 100.0% (25/25) | 0.0% (0/23) | 0.0% (0/14) | 0.0% (0/15) |
|  | Death | 0.0% (0/25) | 0.0% (0/23) | 0.0% (0/14) | 6.7% (1/15) |
|  | Mild | 0.0% (0/25) | 100.0% (23/23) | 0.0% (0/14) | 0.0% (0/15) |
|  | Moderate | 0.0% (0/25) | 0.0% (0/23) | 92.9% (13/14) | 0.0% (0/15) |
|  | Severe | 0.0% (0/25) | 0.0% (0/23) | 7.1% (1/14) | 93.3% (14/15) |
| **Comorbidity** |  |  |  |  |  |
|  |  | 4.0% (1/25) | 26.1% (6/23) | 78.6% (11/14) | 86.7% (13/15) |
| **Days of symptoms to sampling** |  |  |  |  |  |
|  |  | 8 (6.75–10.25) | 13 (12–15.75) | 12 (9.5–14.5) |  |
| **Admission days** |  |  |  |  |  |
|  |  | – | – | 9.5 (6.5–11) | 34 (24.5–53) |
| **Admission days to sampling** |  |  |  |  |  |
|  |  | – | – | 4 (3–5) | 4 (2–6) |
| **ICU** |  |  |  |  |  |
|  | ICU | – | – | 7.1% (1/14) | 100% (15/15) |
|  | ICU days | – | – | 11 (11–11) | 22.5 (10.75–28.75) |
|  | ICU days to sampling | – | – | – | 3 (1.5–5) |
| **Symptoms** |  |  |  |  |  |
|  | Fever | – | – | 92.9% (13/14) | 86.7% (13/15) |
|  | Respiratory | – | – | 100% (14/14) | 93.3% (14/15) |
|  | Gastrointestinal | – | – | 42.9% (6/14) | 28.6% (4/14) |
|  | Musculoskeletal | – | – | 64.3% (9/14) | 60.0% (9/15) |
|  | Sensory | – | – | 21.4% (3/14) | 26.7% (4/15) |
|  | Headache | – | – | 35.7% (5/14) | 0.0% (0/15) |
| **Xray Infiltrates** |  |  |  |  |  |
|  |  | – | – | 71.4% (10/14) | 86.7% (13/15) |
| **Xray pneumonia with consolidation** |  |  |  |  |  |
|  |  | – | – | 64.3% (9/14) | 50% (7/14) |
| **Xray Pleural Effusion** |  |  |  |  |  |
|  |  | – | – | 14.3% (2/14) | 7.1% (1/14) |
| **Treatments** |  |  |  |  |  |
|  | Oxygen | – | – | 64.3% (9/14) | 100% (14/14) |
|  | Invasive Vent | – | – | 7.1% (1/14) | 78.6% (11/14) |
|  | Inotropes | – | – | 0.0% (0/14) | 35.7% (5/14) |
|  | Haemofiltration | – | – | 0.0% (0/14) | 14.3% (2/14) |
|  | Corticoids before sample collection | – | – | 14.3% (2/14) | 66.7% (10/15) |
|  | Antibiotics | – | – | 100% (14/14) | 100% (15/15) |
|  | Antiviral | – | – | 78.6% (11/14) | 100% (15/15) |
|  | Antimalarial | – | – | 100% (14/14) | 100% (15/15) |
|  | Monoclonal antibody | – | – | 0.0% (0/14) | 46.6% (7/15) |
| **Analytical parameters** |  |  |  |  |  |
| | White Cells ($10^9/L$) | – | – | 5.505 (4.82–6.3725) | 9.41 (5.275–12.36) |
| | Platelets ($10^9/L$) | – | – | 273 (227.25–400.75) | 203 (129–255.5) |
| | Neutrophils ($10^9/L$) * | – | – | 3.095 (2.445–3.78) | 6.89 (3.185–10.55) |
| | Lymphocytes ($10^9/L$) * | – | – | 1.32 (1.1875–1.52) | 0.47 (0.415–0.905) |
| | Monocytes ($10^9/L$) | – | – | 0.42 (0.29–0.58) | 0.33 (0.195–0.475) |
|  | Fibrinogen (mg/dl) * | – | – | 500 (500–500) | 373.5 (318–450) |
|  | Creatinine (mg/dl) | – | – | 0.75 (0.58–1.03) | 0.8 (0.765–0.94) |
|  | CRP (mg/L) | – | – | 23.39 (15.5025–53.63) | 48.255 (11.15–99.6225) |
|  | PCT (mg/ml) | – | – | 0.06 (0.04–0.1375) | 0.15 (0.105–0.2725) |
|  | Ferritin (ng/dl) * | – | – | 398.5 (139.75–491) | 1584.5 (704–2934) |
|  | D-Dimer (ng/ml) * | – | – | 518 (402–763) | 2038 (861–7832) |
|  | Lactate (mmol/L) * | – | – | 0.9 (0.85–0.95) | 1.72 (1.42–2.39) |
|  | Maximum CRP (mg/L) * | – | – | 49.225 (17.795–123.78) | 134.205 (71.9475–172.29) |
| | Maximum Neutrophils ($10^9/L$) * | – | – | 4.56 (3.4775–6.8) | 9.715 (5.85–11) |
| | Minimum Lymphocytes ($10^9/L$) * | – | – | 0.87 (0.7–1.1675) | 0.43 (0.345–0.4775) |

The study was approved by the Ethical Committee of Clinical Investigation of Galicia (CEIC ref. 2020/178) and was conducted in accordance with the principles of the Declaration of Helsinki and all applicable Spanish legislation, namely Biomedical Research Act (Law 14/2007-3 of July), Act 41/2002 on the Autonomy of the Patient, Decree SAS/3470/2009 for Observational Studies, and 15/1999 Data Protection Act. Written informed consent was obtained for subjects before study inclusion.

### Sample collection and processing

Blood, nasal epithelium, and saliva samples were sampled from patients and controls at the same timepoints. Blood samples (2.5ml) were collected in PAXgene tubes (BD). RNA was isolated using PAXgene blood miRNA extraction kit (Qiagen) following manufacturer recommendations and carrying out the on-column DNase I treatment during the extraction process. Saliva and nasal epithelium samples were collected in Oragene CP-190 kit (DNA Genotek). We used sponges (SC-2 kit from DNA Genotek) to assist saliva collection in ICU patients unable to spit. RNA from saliva and nasal epithelium specimens was isolated using 500μl of sample and the RNeasy microkit (Qiagen). We used Oragene RNA neutralizer solution (DNA Genotek) to precipitate impurities and inhibitors and slightly modified the protocol provided by the extraction kit as recommended by the Oragene tubes supplier. After isolation, RNA amount and integrity was checked using TapeStation 4200 (Agilent), and DV200 values were calculated to both ensure that >50% of the of the RNA fragments were above 200nt and estimate the optimal sample input for the nCounter analysis. Universal Transport Medium tubes (UTM) nasopharyngeal samples were collected to check for the presence as well as the viral load of SARS-CoV-2 at the same time as collection of blood, saliva, and nasal epithelium samples. Molecular detection of viral particles was performed through a multiplex real-time PCR using the Allplex™SARS-CoV-2 Assay (Seegene).

#### Nanostring nCounter expression assay

Immunological gene expression patterns of different tissues from COVID-19 patients and controls were evaluated through the *SPRINT* nCounter system (NanoString Technologies) and the Immunology Panel (594 genes involved in several immunological processes). This panel covers the main immunological routes, including major classes of cytokines and their receptors, chemokine ligands and receptors, IFNs and their receptors, the TNF-receptor superfamily, and the KIR family genes. We used the standard gene expression protocol for 12x RNA hybridization with 5μl of RNA as input, a hybridization time of 18h for all samples and mixing control and COVID-19 samples to avoid technical sample bias. We first carried out a quality control (QC) of the raw data checking some technical parameters following manufacturer recommendations to verify the absence of technical problems. Samples that did not pass technical QC, or with low number of genes detected, were excluded for downstream analysis.

#### Sample normalization and differential expression analysis

For the longitudinal multi-tissue experiment, normalization of the gene expression data was carried out for each tissue separately by selecting the optimal number and best reference candidate from the set of housekeeping (HK) genes included in the panel, as well as other endogenous genes that showed high levels of stability between samples. The *GeNorm* algorithm (38) implemented in the R package *CrtlGene* (39) was used to test for the most stable and optimal number of genes for normalization. We excluded HK candidate genes with less than 50 raw counts in any of the samples compared from the reference selection analysis. Data normalization was performed through a combined approach using both *DESeq2* (40) and *RUVSeq* (41) as described in (42). Genes with counts below the background (mean + 2 standard deviations (SD) of the negative control spikes in the code set) were not included in the differential expression (DE) analysis. After data normalization and background thresholding, we eliminated lower expressed genes from the list of DEG, retaining a total of 495 genes in blood samples, 544 genes in nasal epithelium samples and 545 genes in saliva samples for downstream analysis. DEG between cohorts were obtained using *DESeq2* (40). We used age and sex as covariables to correct the model. Volcano plots and heatmaps were built with *EnhancedVolcano* (43) and *ComplexHeatmap* (44) R packages, respectively. All graphics were created using R (www.r-project.org) software and the *ggplot2* package (45). Wilcoxon test was used to assess statistical significance between patient groups, and Spearman test for the correlation indices. We focused on highly statistically significant expression changes by considering as thresholds an adjusted *P*-value < 0.05, and log_2_FC ≥ |1.5| for downstream analysis. For the transversal multi-tissue experiment we sub-selected 21 patients for which samples from the three tissues were available (mild: *n* = 11; severe: *n* = 10). We employed a complex model design including the tissue/severity interaction to find tissue-specific severity effects. As reference for the analyses, we used blood and the mild category. We followed the same procedure as in the longitudinal analysis but data from the three tissues were normalized together after selecting a set of adequate reference genes.

### Pathway analysis

We followed both over-representation (ORA) and gene-set enrichment (GSEA) (46) approaches to identify biological functions, cellular components and pathways related to the differences in gene expression. For ORA we used the DEG as input (threshold: log_2_FC ≥ |1.5|), and the genes included in the expression panel as pool for statistical calculations. For GSEA we used all available molecular measurements (log_2_FC) from the genes included in the DE to detect coordinated changes in the expression of genes from same pathway. Analyses were carried out using the *Clusterprofiler* (47) R package. We applied the Benjamini-Hochberg (H-B) procedure for multiple test correction and thresholds were set to 0.05. We tested both the GO (Gene Ontology) and Reactome databases. Graphics were obtained with the R package *enrichplot* (48).

### Validation analysis

We used three additional datasets to validate and check the specificity of the transcriptomic findings using independent data: a) Thair et al. (23) included RNA-seq transcriptomes for 62 SARS-CoV-2 positive and 24 healthy controls; and b) Aschenbrenner et al. (24) included RNA-seq transcriptomes for 39 COVID-19 patients and 10 healthy controls. We additionally checked the specificity of the candidate genes obtained to differentiate COVID-19 from other viral infections using log_2_FC values reported in (23), through the comparison of other mixed viral infections (*n* = 652) and healthy control samples (*n* = 672). Independently, to further test the status of our best candidate genes in bacterial and other viral infections, we carried out an independent validation study by meta-analyzing samples from bacterial (*n* = 382) and viral (*n* = 513) infections, and healthy controls (*n* = 215). Microarray and RNA-seq data were downloaded from the public gene expression microarray repository Gene Expression Omnibus (GEO) under accession numbers: GSE64456 (49), GSE40012 (50), GSE42026 (51), GSE60244 (52), those also included in Thair et al. (23) as viral cohorts; and GSE72829 (53), GSE69529 (54). To merge and integrate independently viral *vs*. healthy controls and bacterial *vs*. healthy controls, we first normalized and pre-processed each dataset separately using *Lumi* (55) for Illumina® microarrays data and *Oligo* (56) for Affymetrix® datasets, while RNA-seq data were pre-processed as described previously (57). Then, we merged these databases retaining only the common genes. Subsequently, we used the R package *COCONUT* to combine all datasets into one and reduce batch effects in the meta-analysis (58). Next, we used *Limma* package (59) to detect DEG and calculate log_2_FC values between groups. Scatterplots of log_2_FC values and Pearson correlation indexes were obtained for the pseudo-validation analysis.

### Leukocyte proportion estimation from gene expression data

We imputed immune cell subset abundances from nCounter expression data in different patient categories, including mild and healthy controls, from which real immune cell counts from blood tests were not available. We used the CIBERSORTX (60) tool and a leukocyte gene signature matrix (61) obtained from microarray data as reference. We applied a B-mode batch correction to account for cross-platform variation and 500 permutations for the analysis. We graphically represented immune cell proportions using R software and carried out a Wilcoxon test to assess statistical significance between categories.

### Data availability

Expression data that support the findings of the present study have been deposited in Gene Expression Omnibus (GEO) database under the accession code GSE183071.

## Results

### Cohort description

Most of the clinical features were only available for hospitalized patients (**Table 1**). Similar median symptoms days to sampling were obtained for mild, moderate, and severe cases, as well as admission days to sampling in the case of moderate and severe patients (*P*-value = 0.236 and 0.704, respectively). During admission, oxygen support was required for 64.3% of the moderate cases and 100% of the severe cases (78.6% required invasive ventilation). All moderate cases were treated with a double therapy including antibiotic and antimalarial drugs (in 78.6% of the moderate cases, antiviral therapy was also administrated). Severe cases received a combined triple therapy of antibiotic, antiviral (lopinavir and ritonavir) and antimalarial (hydroxychloroquine) treatment, and 46.6% of them received tocilizumab.

No statistical differences were observed in viral loads at the time of sample collection depending on COVID-19 severity, nor statistical correlation between days of symptoms to sample collection and Ct values from the SARS-CoV-2 qPCR assay (**S1A Fig in S1 Appendix**) on nasopharyngeal swabs obtained at the point of sample collection. However, we obtained statistically significant differences in some analytical parameters between severe and moderate cases at the sampling timepoint: higher neutrophil counts (*P*-value = 0.012), ferritin (*P*-value = 0.002), D-Dimer (*P*-value = 0.006) and lactate (*P*-value = 0.019) values, but lower lymphocyte counts (*P*-value = 0.002) and fibrinogen (*P*-value = 0.004) values in severe cases. Maximum values for C-reactive protein (CRP) and neutrophils (*P*-values = 0.02 and 0.006, respectively) and minimum values for lymphocytes (*P*-value < 0.001) also showed statistically significant differences between these groups.

Immune cell fractions inferred from gene expression data also pointed to a significantly lower proportion of T lymphocyte subpopulations CD4 and CD8 in severe cases than in other severity categories and healthy controls (**S1B Fig in S1 Appendix**). Likewise, the proportion of B cells was significantly higher in mild when compared to moderate and severe cases. Cell numbers of CD4 and NK subpopulation also showed a significant decrease with disease severity. Finally, a higher proportion of monocytes was also observed in moderate cases, correlating well with real monocyte counts detected in the blood tests from moderate and severe cases.

### Transcriptomic immune response in blood

Principal component analysis (PCA) of blood gene expression revealed a moderate differentiation by severity in the first principal component (PC1; Fig 1A). When we compared the whole COVID-19 cohort against controls we obtained nine DEG (**S2 Table**, Fig 1A**; S2 Fig in S1 Appendix**), all of them over-expressed in patients except for *FCER1A* gene that is under-expressed (log_2_FC = -1.7; adjusted *P*-value = 2.3E-5). The most statistically significant gene was *TNFRSF17* gene (log_2_FC = 1.9; adjusted *P*-value = 2.4E-10), followed by *FCER1A* and *SERPING1* genes. Three out of the nine DEG were found to be still significant when using a model that accounted for differences in treatment with immunomodulators (steroids), administered only in some severe and moderate patients: *TNFRSF17* (log_2_FC = 1.72; adjusted *P*-value = 4.1E-6), *SERPING1* (log_2_FC = 2.2; adjusted *P*-value = 4.3E-4) and *MX1* (log_2_FC = 1.97; adjusted *P*-value = 3.0E-3) genes. To test the robustness of this COVID-19-related immune signature we carried out a validation in two recently published COVID-19 RNA-seq datasets (23, 24) (**S3A Fig in S1 Appendix**). The 9-transcript immune signature correlated very well with the data from the two validation cohorts (R = 0.96, *P*-value = 5.3E-5; R = 0.92, *P*-value = 4.0E-4; respectively). As in our study, all genes were DE in both datasets when comparing patients *vs*. healthy controls.

**Fig 1.**
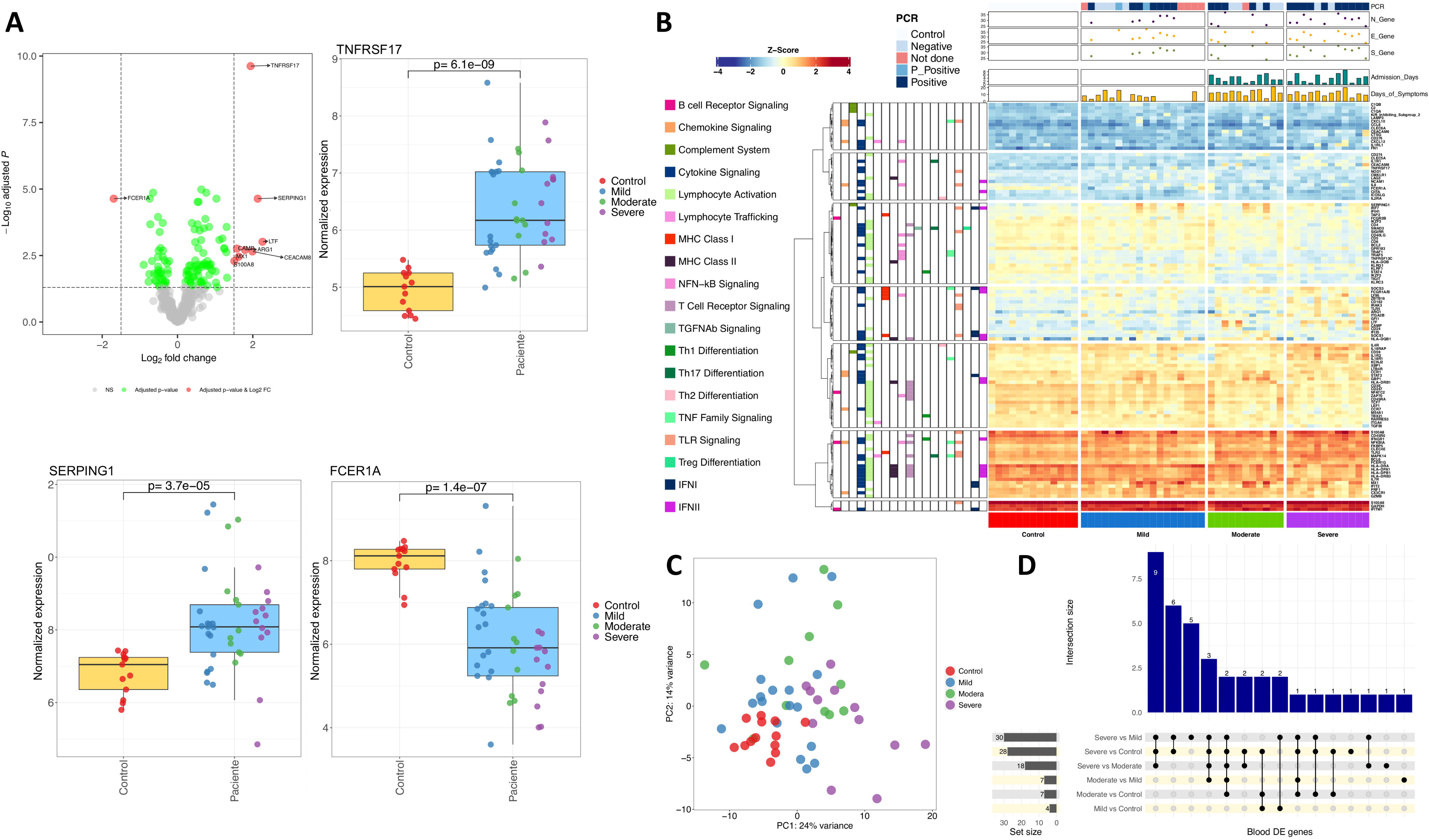
A) Volcano plot showing the DEGs between cases and healthy controls in blood samples. Boxplots of the most relevant genes and the *P*-value of the statistical test are also shown. B) Heatmap and cluster analysis of the DEGs between all categories in blood samples. Only genes with a log_2_FC |>1| were represented. Gene clusters were generated using k-means partitioning. C) PCA of immune transcriptomic data from blood samples of the different COVID-19 categories analyzed using all genes from the panel detected above the background. D) Upsetplot of the common DEGs between categories compared in blood samples.

Next, we studied the specificity of this 9-transcript signature by examining other viral and bacterial infections. Firstly, transcriptomes of different viral infections (*n* = 655) and healthy controls (*n* = 672) from Thair et al. (23) were contrasted with the results from our study (**S3 Fig in S1 Appendix**). We found a moderate correlation (R = 0.85; *P*-value = 0.0037) between log_2_FC values from our study (COVID-19 *vs*. control) and their study (viral infections *vs*. healthy controls). Notably, *TNFRSF17* gene (log_2_FC = 0.16; adjusted *P*-value = 0.50) did not show appreciable differences between non-COVID-19 viral infections and controls. To further test the differential pattern of this gene in COVID-19 patients, we used a combined dataset from different microarray and RNA-seq studies that includes transcriptomes from viral and bacterial infected patients, and healthy controls. We consistently observed again a moderate correlation between our results and those from the viral infections *vs*. healthy control dataset (R = 0.81, *P*-value = 0.0078), and a weaker correlation with those from bacterial infections *vs*. healthy control dataset (R = 0.61, *P*-value = 0.081). Again, *TNFRSF17* gene showed a log_2_FC value very close to 0, and non-significant results in the viral combined dataset (log_2_FC = -0.30; adjusted *P*-value = 0.28). Furthermore, in the bacterial combined dataset, the *TNFRSF17* gene (and *MX1*) was under-expressed in bacterial infections *vs*. controls, even though this contrast was also statistically significant (log_2_FC = -0.81, adjusted *P*-value = 1.9E-4) as in COVID-19 datasets.

We performed a more detailed analysis of the immunological processes underlying COVID-19 by comparing different severities (Fig 1B; Fig 1C; **S4 Fig in S1 Appendix**). We found a similar number of DEG when comparing severe cases *vs*. healthy controls and severe *vs*. mild patients (28 and 30 respectively; **S2 Table**; Fig 1D), pointing to a limited systemic immune response (Fig 1D**; S4 Fig in S1 Appendix**). We found 14 common DEG in all comparisons involving severe cases (Fig 1D); *ARG1* gene was the most remarkable one over-expressed in severe cases (Log_2_FC = 3.98 – 4.71; adjusted *P*-value = 1.53E-18 – 1.78E-22) (**S2 Table**). Other significatively over-expressed genes in severe cases were those related to pro-inflammatory pathways *IL-1* (*IL1R1* and *IL1R2*) and *IL-18* (*IL18R1*) and immunoregulatory functions (*FKBP5* and *S100A8*); **S5 Fig in S1 Appendix**. To further test this 14-transcript COVID-19 signature for severe patients, we compared our results to an independent cohort (24); after excluding two genes (*ZBTB16* and *IL1RL1*) that were not found to be DE between their severe cases *vs*. healthy controls (24), we found a notable correlation between both datasets (R = 0.87; *P*-value = 0.00023); **S3 Fig in S1 Appendix**.

Analysis of moderate *vs*. healthy controls and mild cases *vs*. healthy controls returned a few DEG coinciding with the previous 14-transcript signature, e.g. *TNFRSF17* and *SERPING1*. *TNFRSF17* represents the strongest signal of COVID-19 disease of all the genes analyzed, with an increased gene expression associated with severity (log_2_FC values compared to healthy controls: 1.66 in mild, 1.95 in moderate, and 2.71 in severe cases); **S2 Table**. Conversely*, FCER1A* gene was under-expressed in patients and shows a gradual decreased gene expression with severity (log_2_FC values compared to healthy controls: not DE in mild, -1.83 in moderate, and -3.06 in severe cases). Genes *MX1* and *C1QB* showed an increased expression in mild (log_2_FC = 2.10; adjusted *P*-value= 1.32E-5) and moderate cases (log_2_FC = 1.78; adjusted *P*-value = 5.59E-5) respectively, with respect to controls (**S2 Table**). When comparing DEGs between hospitalized and non-hospitalized patients, the most relevant genes were *CAMP*, *LTF*, *CEACAM8* and *CEACAM6*, all involved in neutrophil degranulation processes triggered by the innate immune response (**S6 Fig in S1 Appendix**).

Pathway analysis by way of a GSEA of severe cases *vs*. healthy controls and using Reactome revealed: (*i*) an up-regulation of innate immune processes (mainly driven by neutrophil activation and degranulation, the *IL-1* pathways, and the TLR cascade (**S3 Table**, **S7 Fig in S1 Appendix**); and (*ii*) down-regulation of the adaptative immune response and related pathways such as Translocation of ZAP−70 to Immunological synapse, T cell antigen receptor (TCR) signalling, co-stimulation of CD28 family, and PD−1 signalling. Meanwhile, GSEA analysis using GO showed a comparable pattern: (i) up-regulation of the neutrophil mediated immunity and downstream processes; (*ii*) under-regulation of B cell proliferation/activation; and (*iii*) antigen receptor-mediated signalling (**S3 Table**, **S7 Fig in S1 Appendix**). Similar findings were observed in comparisons of severe *vs*. moderate (**S3 Table**, **S8 Fig in S1 Appendix)** and severe *vs*. mild cases (**S3 Table**, **S9 Fig in S1 Appendix**) from Reactome, except for the IFN pathway, which is under-regulated in severe cases in comparison to moderate and mild patients. GO pathway analysis found under-expression of genes involved in antigen processing and presentation *via* MHC-II complex and viral defense. The only process detected by GSEA in mild cases were an up-regulation of the IFN signalling pathway with the Reactome database; whereas routes related to host response to viral infection were the most over-regulated in GO (**S3 Table**, **S10 Fig in S1 Appendix)**. Overall, there exists a significant up-regulation of the neutrophil activation and degranulation pathways, which increase with severity, and a lack of IFN response in severe cases. The differential neutrophil-driven immune response observed in moderate and severe patients is also detected when comparing hospitalized against non-hospitalized cases and healthy controls (**S11 Fig in S1 Appendix; S3 Table)**.

### Transcriptomic immune response in nasal epithelium

PCA indicates a differentiation of severe cases against controls that is only notable in the PC2 dimension (12% of the variation); Fig 2A (see also heat maps in Fig 2B). Conversely, mild and moderate cases show more remarkable differences with severe and controls in the PC1 dimension (23% of the variation). Of the seven DEGs observed in the comparison patients *vs*. healthy controls, one is under-expressed and six over-expressed (**S4 Table;** Fig 2C). Thus, *FCER1A* is under-expressed in patients when compared against controls (log_2_FC = -1.63; adjusted *P*-value = 0.004; severe *vs*. controls: log_2_FC = -3.28; adjusted *P*-value = 2.75E-9; severe *vs*. moderate: log_2_FC = -2.01; adjusted *P*-value = 0.0004; severe *vs*. mild: log_2_FC = -1.96; adjusted *P*-value = 0.00097), even though it was not detected as DE when comparing moderate and mild cases with healthy controls (**S4 Table**; however the absolute expression values showed only weak statistical significance in mild and moderate patients *vs*. healthy controls; Fig 2D). Of the over-expressed genes, *SERPING1* gene leads the DEG list; the main signal comes from the high expression levels observed in moderate and mild patients against controls; this gene was however not DE in severe cases when compared with controls. Thus, *SERPING1* over-expression decreased gradually from mild to severe disease (**S4 Table;** Fig 2D). A similar pattern was observed for other genes (*CXCL9*, *LAG3* and *GZMB*), all showing a gradual over-expression in moderate and mild cases against controls (**S4 Table**; **S12 Fig in S1 Appendix**) but not DE in severe patients. Overall, gene expression of immune genes in nasal epithelium samples showed a very similar response to infection in mild and moderate patients, with seven overlapping DEGs (*LAG3*, *SERPING1*, *CXCL9*, *CXCL10*, *CXCL11, IFITM1* and *GZMB*; all over-expressed) between both categories *vs*. controls, but no DEG when comparing mild *vs*. moderate cases (Fig 2C**; S13 Fig in S1 Appendix; S14 Fig in S1 Appendix**). Despite the similarities in immune response of both phenotypic categories, these genes showed a gradual increase in expression from severe to mild disease; the differences were found to be more notable when comparing mild *vs*. controls than in moderate *vs*. controls (**S4 Table**).

**Fig 2.**
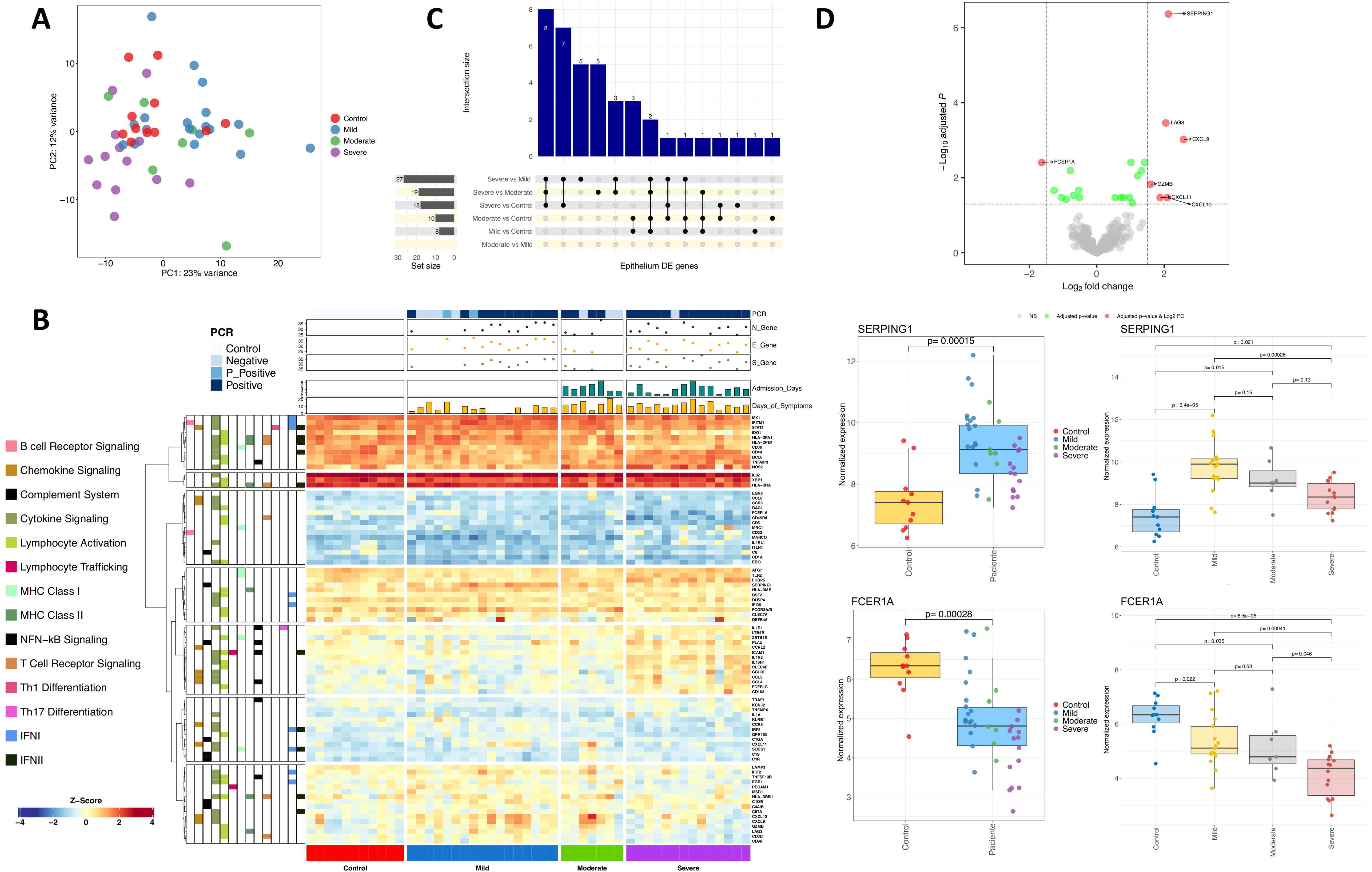
Fig 1. A) PCA of immune transcriptomic data from nasal epithelium samples of the different COVID-19 categories analyzed using all genes from the panel detected above the background. B) Heatmap and cluster analysis of the DEGs between all categories in nasal epithelium samples. Only genes with a log_2_FC |>1| were represented. Gene clusters were generated using k-means partitioning. C) Upsetplot of the common DEGs between categories compared in nasal epithelium samples. D) Volcano plot showing the DEGs between COVID-19 samples and healthy controls in nasal epithelium samples. Boxplots of the most relevant genes and the *P*-value of the statistical test are also shown.

The comparison of severe *vs*. controls/moderate/mild yielded 18, 19 and 27 DEGs genes, respectively (Fig 2C; **S4 Table**); eight genes are shared between these three comparisons (*FCER1A*, *IL18R1*, *IL1R2*, *CCL3 CCL4*, *CLEC4E*, *CD163* and *MARCO*), all over-expressed in severe cases (except *FCER1A*) and all mainly involved in the innate immune and inflammatory response (**S15 Fig in S1 Appendix**). *IL18R1* and *IL1R2* appeared in the top three most DEGs in severe cases compared to all categories, with a log_2_FC ∼2 in severe *vs*. moderate cases, and even higher in severe *vs*. controls/mild cohorts. Other under-expressed genes in severe compared to mild/moderate cases included *IFIT2* (Interferon Induced Protein with Tetratricopeptide Repeats 2) and *IDO1*, whose encoded protein is produced in response to inflammation and has autoimmune protective features (**S4 Table**; **S13 Fig in S1 Appendix**).

We did not find statistically significant pathways in the nasal samples from severe cases *vs*. controls using both GSEA and ORA. Moderate *vs*. controls comparisons highlighted some up-regulated pathways in GSEA mainly connected to interaction and response to other organism processes (**S5 Table**), and inhibition of endopeptidase activity (**S16 Fig in S1 Appendix)**. Comparison of mild cases *vs*. controls using ORA suggested two main processes: (*i*) antimicrobial immune humoral response (an adaptative immune response mainly driven by B lymphocytes), and (*ii*) chemokine related pathways (**S5 Table**; **S17 Fig in S1 Appendix**). The comparison of severe *vs*. mild patient revealed up-regulated coordinated responses in various nested pathways involving NAD metabolism (**S5 Table**), a regulator of infection and inflammation. Molecular functions showing down-regulation in severe *vs*. mild cases were MHC protein binding and hydrolase activity (**S18 Fig in S1 Appendix)**. ORA analysis of DEGs revealed alterations in chemokine dynamics-related pathways, with over-expression of CC-type chemokines and *MARCO*, and under-expression of CXC-type chemokines and *DEFB4A* (**S18 Fig in S1 Appendix)**. The comparison of severe *vs*. moderate patients using GSEA analysis showed some similarities with that of severe *vs*. mild cases (**S19 Fig in S1 Appendix**). Reactome and GO showed some common biological processes and/or cellular components, such as under-regulation of type I and II IFN pathway and MHC class II antigen presentation, and an over-regulation of neutrophil activation and degranulation processes. When exploring pathways associated with the comparison of moderate *vs*. mild cases, GSEA analysis found an upper-regulation of serine−type endopeptidase activity and a down-regulation of molecular transducer receptor activities (**S20 Fig in S1 Appendix; S5 Table**).

### Transcriptome immune response in saliva

Among the genes detected in saliva samples, there was no DEG when comparing patients and healthy controls. This fact is reflected in a PCA (and heatmap analysis), which shows a large cluster with overlapping controls, mild and moderate, and only slightly differentiation of the severe cases in the PC1 (35% of the variation) and the PC2 (20% of the variation); Fig 3A,D. Therefore, the data seem to indicate very limited immune response to infection in the oral cavity of mild and moderate patients. A few DEGs were detected when comparing hospitalized with non-hospitalized patients (Fig 3B**; S21 Fig in S1 Appendix**) and against controls (**S6 Table**). As expected, most DEGs are the same in both comparisons, with 11 out of the 12 DEGs detected in hospitalized *vs*. non-hospitalized cases also appearing in hospitalized *vs*. controls, all of them over-expressed (**S21 Fig in S1 Appendix**; **S6 Table)**. The strongest differences in hospitalized *vs*. non-hospitalized patients were represented by (*i*) *PLAU*, a gene encoding a secreted serine endopeptidase that transforms plasminogen in plasmin and is implicated in the rearrangement of extracellular matrix and cell migration, and (*ii*) *LGALS3,* which plays a role in apoptosis, innate immunity (attracting macrophages and activating neutrophils), cell adhesion and T-cell regulation processes as well as in acute inflammatory responses (Fig 3C). These two genes were the only ones detected as DE when comparing moderate *vs*. mild cases; they probably constitute the strongest and most specific immune signature in oral mucosa from hospitalized individuals (**S6 Table; S22 Fig in S1 Appendix**). The other DEGs of the signature from hospitalized samples also exhibited expression levels that increased with severity (**S22 Fig in S1 Appendix)**.

**Fig 3.**
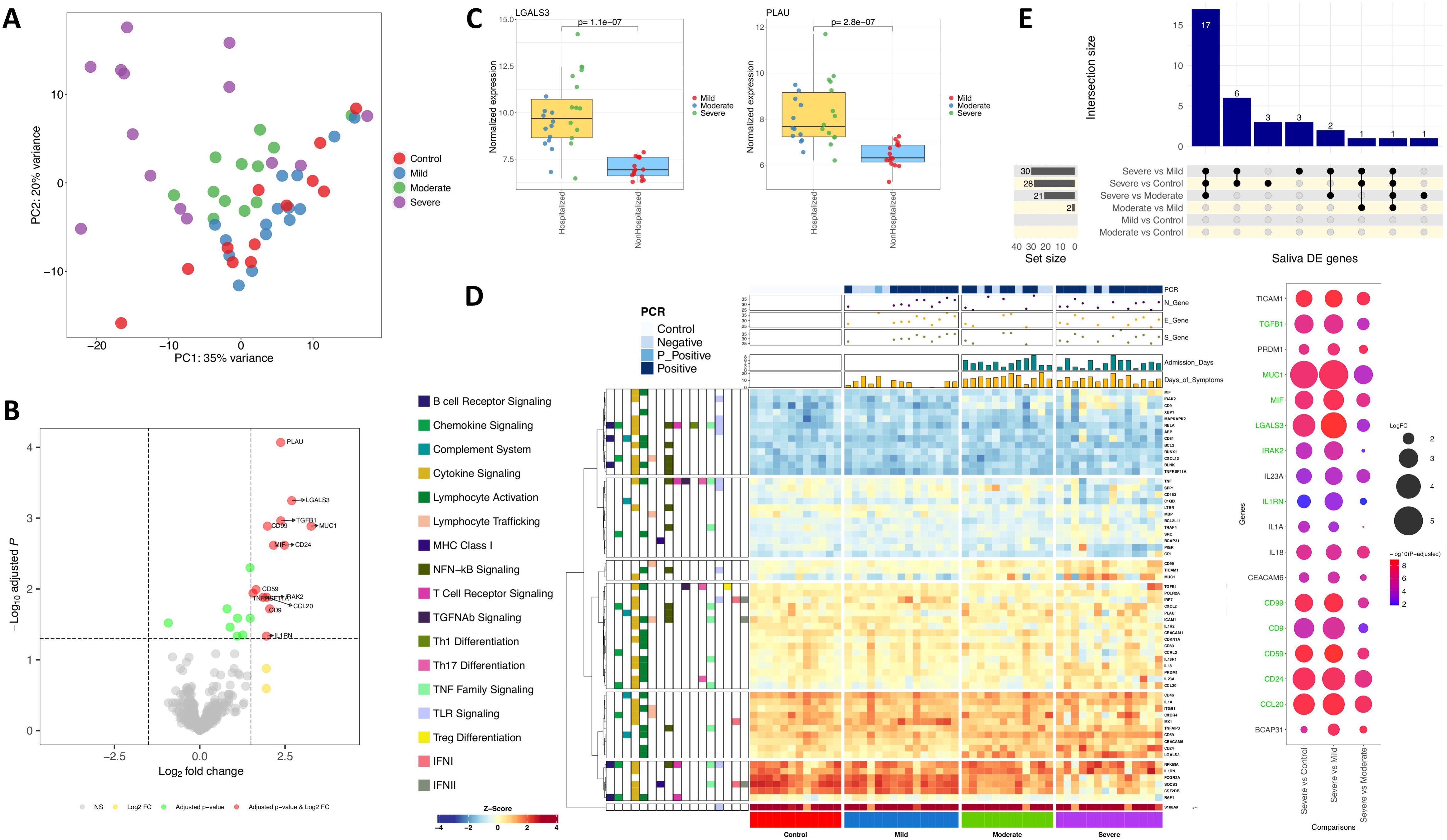
A) PCA of immune transcriptomic data from saliva samples of the different COVID-19 categories analyzed using all genes from the panel detected above the background. B) Volcano plot showing the DEGs between hospitalized and non-hospitalized saliva samples. C) Boxplots of the most relevant genes from the comparison of hospitalized *vs.* non-hospitalized saliva samples and the *P*-value of the statistical test. D) Heatmap and cluster analysis of the DEGs between all categories in saliva samples. Only genes with a log_2_FC |>1| were represented. Gene clusters were generated using k-means partitioning. E) Upsetplot of the common DEGs between categories compared in saliva samples. Bubble plot represents log_2_FC and adjusted *P*-value from the 18 DEGs between severe and all remaining categories. Green color in gene names indicates genes also included in the hospitalized signature.

We found 18 common over-expressed DEGs that differentiate severe cases from the other phenotypic categories (Fig 3E**; S23 Fig in S1 Appendix**). *CCL20* showed the highest log_2_FC value in all comparisons (**S6 Table**). As expected, higher differences were found in severe *vs*. mild and severe *vs*. controls (Fig 3E). Eleven out of the 18 genes were also included in the signature that differentiated hospitalized from both non-hospitalized and healthy controls, respectively (Fig 3E; **S24 Fig in S1 Appendix**).

Pathway analysis in severe *vs*. controls using GSEA found some moderate up-regulation of processes and cell components, but only phagocytic vesicle as under-regulated cellular component (**S25 Fig in S1 Appendix**). Some of these pathways were also highlighted as up-regulated in severe *vs*. mild patients but, in this case, a positive regulation of cell migration and motility was also detected (**S7 Table; S26 Fig in S1 Appendix**). GSEA analysis of severe *vs*. moderate patients provided the highest number of hits from all comparisons, all showing up-regulation in severe cases, but moderate NES values (**S7 Table; S27 Fig in S1 Appendix**). Among the up-regulated pathways in moderate *vs*. mild (**S28 Fig in S1 Appendix**) cases there are pathways related to cell migration, receptor activator activity and chemoattractant activity, of which the latter two were also up-regulated in moderate cases *vs*. controls (**S29 Fig in S1 Appendix**). GSEA also pointed to a strong under-regulation of the IFN signalling in moderate *vs*. mild cases in all the reference databases interrogated (**S7 Table; S28 Fig in S1 Appendix**). Notably, IFN pathway was not detected in severe *vs*. mild cases, however it appears as highly under-regulated in hospitalized *vs*. non-hospitalized patients, although not when comparing to controls.

Finally, comparison of mild cases and controls resulted in several under-regulated processes and cellular components mainly involved in cell junction, cell communication and secretory vesicles (**S7 Table**; **S30 Fig in S1 Appendix**).

### Transversal multi-tissue analysis

We selected a subset of severe and mild patients (*n* = 21) with samples available from all tissues collected at the same time. We explored for interactions between severity and tissue to elucidate immunological features that could illuminate the causes for the differential evolution observed in patients. Only when relaxing the thresholds for this interaction analysis (adjusted *P*-value < 0.05 and a cumulative log_2_FC > |1.4|) we found that the effect of severity for the *CXCL9* gene is significant for all tissue comparisons, being much lower in nasal compared to blood tissue (*P*-adjusted = 1.41E-6; cumulative log_2_FC = -2.94) and moderately lower for both saliva *vs*. blood (*P*-adjusted = 0.03; cumulative log_2_FC = -1.42) and nasal *vs*. saliva (*P*-adjusted = 0.017; cumulative log_2_FC = -1.51) tissues (**S8 Table**). Looking at the *CXCL9* normalized data, we found higher expression in severe than in mild cases for blood, with the opposite pattern in nasal and saliva epithelium samples (**S31 Fig in S1 Appendix**). When we increased the significance thresholds (*P*-adjusted < 10E-6; cumulative log_2_FC > |2.5|) a set of 17 genes were found as DE in all possible comparisons (**S8 Table;** Fig 4B; Fig 4C). Thirteen out of these 17 genes were also detected as DE between severe and mild cases in the longitudinal transcriptomic analysis (Fig 4B). The transversal multi-tissue analysis generates four clusters for these 17 genes according to their cumulative Log_2_FC values (Fig 4C). This analysis highlights that the main specific severity signals in blood were driven by gene cluster 1 represented by *ARG1*, *CEACAM8*, *LTF* and *ITGA2B*, and to a lesser extent by cluster 2 (specially *IRAK3* and *TLR5*). This means that severity has a more marked effect in blood in comparison with nasal and saliva samples (Fig 4C; **S8 Table**; **S31 Fig in S1 Appendix**). Cluster 3 (Fig 4C), represented by *CD22* and *TRAF1*, indicates that severity determines an opposite behavior in nasal (lower expression in mild and higher in severe) than in blood (higher expression in mild and lower in severe (**S31 Fig in S1 Appendix**). Finally, cluster 4 (Fig 4C) represented by seven genes indicates a differential effect of severity in saliva with respect to blood and nasal epithelium (especially remarkable for *MUC1* and *LGALS3*).

**Fig 4.**
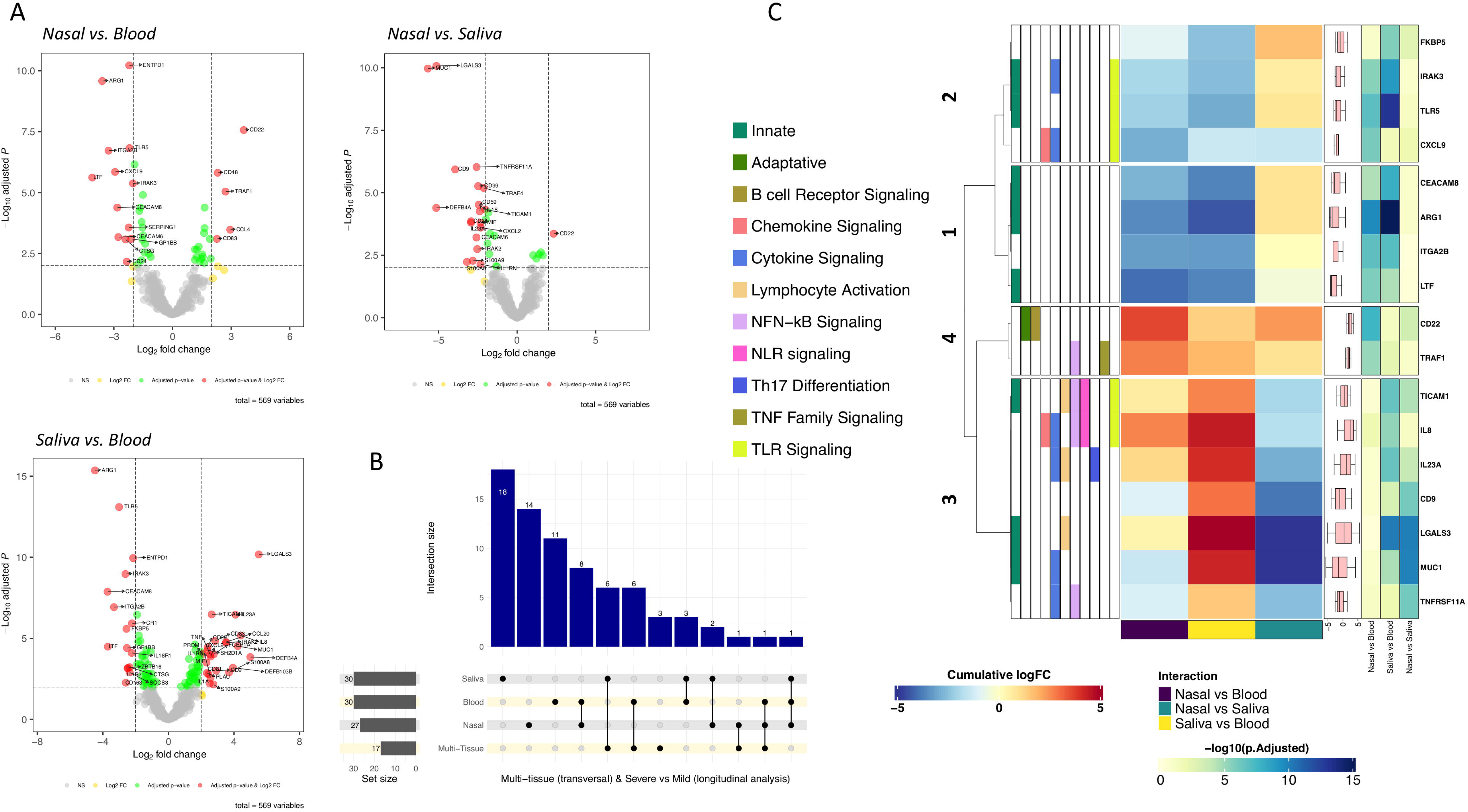
A) Volcano plots showing the DEGs between severities (severe/mild) and different tissue samples (thresholds: *P*-adjusted = 0.01, cumulative log_2_FC = |2|). B) Upsetplot of the DEGs between severe and mild categories in all tissues as well as in the multi-tissue transversal analysis. C) Heatmap and cluster analysis of the DEGs between severities (severe/mild) and different tissue samples (thresholds: *P*-adjusted = 10E-6, cumulative log_2_FC = |2.5|). Gene clusters were generated using k-means partitioning.

## Discussion

Healthcare systems worldwide collapsed during the COVID-19 pandemic due to the high number of hospital admissions, mostly with respiratory complications that, in many cases, ended up being fatal in patients with severe disease. Usually, viral colonization begins in a local tissue environment, with the upper respiratory tract being the main gateway for the virus entrance; however the infection does not always have a significant systemic impact. To explore differential host response to infection, we analyzed the expression of immune response genes in three different tissues from COVID-19 patients affected by different disease severities.

In the analysis of blood samples, we detected specific COVID-19 DEGs, all but one (*FCER1A*) over-expressed, which were subsequently validated in two independent RNA-seq datasets. A few DEGs have comparable expression levels in other viral and bacterial infections, suggesting a common response against severe infections. Moreover, *ARG1* and *FCER1A* show similar expression values independently of the causative pathogen origin (viral or bacterial), but with opposed log_2_FC values (*ARG1*: 1.5 to 3.5; *FCER1A*: -2.5 to -1.5). *ARG1* encodes for the Arginase-1 protein and is expressed in neutrophils, for which a central role in innate immune and inflammatory response has been described, while *FCER1A* is mainly expressed in basophils and mast cells and is involved in the initiation of allergic response. We also observed that *TNFRSF17* is DE in COVID-19 compared to other infections. This gene is expressed in mature B lymphocytes encoding for a member of the TNF-receptor superfamily; it is involved in MAPK8/JNK and NF-kB activation pathways and mediates the development and survival of B-cell lymphocytes to maintain humoral immunity. The number of DEGs in blood increases significantly with severity (**S4 Fig in S1 Appendix**), clearly pointing to a lower immune systemic response in mild and moderate than in severe patients. Moreover, mild and moderate patients over-expressed the interferon type I (IFN-I) pathway in response to viral infection: a timely IFN-I response is critical as it represents a successful front-line protection against viral infections. IFN-I is produced by cells after pathogen-associated molecular patterns (PAMPs) recognition, inducing downstream expression of IFN stimulated genes (ISGs). Furthermore, it has an important role in immune control through the regulation of proinflammatory cytokines, activation and recruitment of different immune cells types, and shaping a successful adaptative immune response (62). SARS-CoV-2 seems to be more susceptible to IFN-I and produces various proteins that antagonize IFN production through different mechanisms, suppressing IFN-I pathway in a more efficient manner than other coronaviruses (63, 64). Interestingly, low IFN-I alpha level has been recently related to a more severe disease outcome, and proposed as a potential therapeutic treatment (65, 66). The IL-1 pathway related genes were detected as over-activated in severe cases. Conversely, moderate and severe patients showed an up-regulation of innate immune response (not observed in mild patients and healthy controls), driven by a different degree of neutrophil response pathways activation. Severe disease was associated with high neutrophil counts and levels of pro-inflammatory cytokines, producing microthrombus formation and organ damage. The release of neutrophil extracellular traps (NETs) could explain (at least partially) COVID-19 vascular complications through forming aggregates that produce vessel occlusion and endothelial damage (67, 68); unbalanced NET production seems to have an important role in viral immunopathology and respiratory disease complications (69). Consistently, we observed a DE of *S100A8*/*S100A9* and *CAMP* genes (both encoding proteins usually identified in NETs) in severe cases *vs*. all other categories. Likewise, the multi-tissue transversal analysis consistently indicated a higher difference in severity between systemic and local immune response with neutrophil related genes (*ARG1*, *LTF*, *CEACAM8*), and other blood coagulation genes that participate in the process of platelet aggregation (*ITGA2B*), which could explain the formation of microthrombus in severe phenotypes. Overall, patterns of exacerbated innate response in hospitalized individuals were accompanied by a global failure of the adaptative immune response in severe cases, with under-regulation of several routes involved in T-cell signalling and maintenance of immune homeostasis, like ZAP-70 signalling. ZAP-70 kinase regulates TCR downstream signaling of antigen presentation, and an adequate regulation of its activity is key for a proper T cell proliferation; furthermore, ZAP protein can restrict SARS-CoV-2 replication (70). Interestingly, routes representing the adaptative immune system were also suppressed in severe cases; this fact could explain the characteristic lymphopenia observed in severe patients. In agreement with an over-activation of the innate and under-activation of adaptative immune response, an unbalanced activation of HLA class I and II antigen presentation was also found in severe cases.

The most prominently over-expressed genes in the nasal epithelium of severe cases were anti-inflammatory IL-1 family receptors IL-1R2 and IL-18R1 (mainly expressed by monocytes-macrophages). Both receptors are involved in the IL-1 and IL-18 cascades, which play a role in the initiation of the inflammation process after PAMPs recognition. Nevertheless, we did not detect DE of the pro-inflammatory cytokines *IL1A*, *IL1B*, *IL1RAP* or *IL18* in severe samples, just a moderate up-regulation of *IL1R1* gene. This imbalance between agonists-antagonists/soluble receptors in severe disease with respect to mild cases was previously noted in broncho-alveolar lavage fluids (BALF) from COVID-19 patients (71), and it could lead to exacerbated inflammatory responses (72). In addition, pathways analysis detected an alteration of NAD+ metabolism in severe cases with respect to those with mild disease, probably pointing to a NAD (and ATP) depletion in host cells from severe patients. It is known that some intracellular pathogens can modulate host cell NAD, impairing several innate immune routes (73). Nasal samples from severe cases also showed similar expression patterns of CXC-chemokines, *GZMB* and antiviral *IFITM1*/*IFIT2* genes compared to controls. Unlike in mild cases, in which main signals came from cell-mediated immunity (Th1, NK and T cytotoxic cells responses) as well as from a robust IFN-II induction, severe cases show an over-expression of genes associated to monocyte-macrophage recruitment; these cells could play a key role in the cytokine storm observed in severe patients (74). For instance, CC pro-inflammatory chemokines genes *CCL3* and *CCL4* act as chemoattractant cytokines in response to tissue injury or inflammation. Some of them have already been highlighted as severity-related biomarkers of COVID-19 in airway epithelial cells (75), and they were found highly expressed in BALF of severe compared to mild cases (71). They are ligands of chemokine receptors CCR1 (CCL3) and CCR5 (CCL3 and CCL4) and have a role in monocyte recruitment. *CD163*, *CLEC4E* and *MARCO* are expressed in macrophages, and the latter two can act as pattern recognition receptors (PRR) for pathogen-associated molecular patterns (PAMPs). The protein encoded by *CLEC4E* can interact with *FCER1G* which, unlike *FCRE1A* gene, is over-expressed in severe cases when compared to controls. Nasal epithelium response in moderate and mild patients was mainly characterized by an over-expression of CXCR3 receptor ligands chemokines *CXCL9*, *CXCL10* and *CXCL11.* These chemokines are highly inducible by IFN gamma, and once joined to the receptor IFN is activated and triggers calcium release to the cytosol, acting as a second messenger to activate downstream signalling pathways (also highlighted in the pathway analysis of nasal samples from mild patients). These chemokines were previously described as biomarkers of respiratory viral infection in nasopharyngeal samples (76). Similarly, other antiviral genes were also over-expressed in mild cases: *IFITM1* has antiviral activity against several enveloped viruses, hampering the viral membrane fusion, and has demonstrated a strong inhibition to SARS-CoV-2 infection by blocking the virus entrance into cells (77); and *IFIT2* is involved in the inhibition of viral replication.

Contrary to the expression profile observed in nasal epithelium samples, there is an absence of appreciable DE in the saliva of mild cases, suggesting a lack of oral immune response against viral infection. Nevertheless, in severe, and to a lesser extent in moderate samples, some up-regulated severity related hits were detected, many of them linked to the viral dynamics. The virus has mechanisms to hijack cellular machinery to their own benefit, altering the levels of several metabolites and cellular processes; one of these molecules are galectins, which have been proposed as biomarkers to monitor viral infections as well as therapeutic targets or antagonists (78). One of the most remarkable over-expressed hits in moderate and severe saliva samples was *LGALS3* (galectin-3), a contributor to pro-inflammatory response against different viral infections (78), enhancing macrophage survival, and positively regulating macrophage recruitment (79), even promoting a more severe course of the infection (80). Elevated levels of galectin-3 were observed in macrophages, monocytes and dendritic cells in severe *vs.* mild COVID-19 patients (81); the regulatory role of galectin-3 in inflammatory response, fibrosis and disease progression, makes it a promising therapeutic target for severe cases (82). We did not observe systemic or nasal epithelium over-expression of this gene. Another important over-expressed gene in saliva was *PLAU* (urokinase-type plasminogen activator), which, together with *CD59* (inhibitor of complement-mediated lysis) and *CD9* (tetraspanin), can facilitate virus-host cell interaction, host immune system viral evasion and, thus, dissemination and viral pathogenesis through buccal epithelium. Urokinase transforms plasminogen to the active form serine protease plasmin; this airway protease (like several others e.g. trypsin, cathepsins, elastase, and *TMPRSS* family members) are required for coronavirus entry enhancement (83). After viral interaction with *ACE2* receptor, furin site in S protein of SARS-CoV-2 is proteolytically cleaved by *TMPRSS2* to enable viral internalization (84). Plasmin has demonstrated to cleave several viral proteins of many respiratory pathogens (85–87) and also S protein in SARS-CoV *in vitro* (88), increasing the ability of the virus to enter host cells. Even though the plasmin protein S cleavage in SARS-CoC-2 has not been demonstrated, there is growing evidence pointing towards a probable role of the plasmin-plasminogen system and fibrinolytic pathway in the COVID-19 severity (89–92). Co-expression and upregulation of *PLAU*, *TMPRSS2*, and *ACE2* was observed in airway epithelial cells from severe/moderate patients and SARS-CoV-2 infected cell lines. This, together with a plasmin-mediated cleavage, could make cells more susceptible to infection (93). Interestingly, we also observed an enhanced expression of *PLAU* in nasal epithelium from severe cases when compared to mild and controls, highlighting the crosslink between both tissues. In the same vein, tetraspanins are molecules associated to specific viruses and seem to play a role in different stages of viral infections (94). Tetraspanin CD9 was involved in early MERS-CoV lung cell entry (95). Recently, it has also been proposed that extracellular vesicles may contribute to viral dissemination by transferring viral receptors, such as CD9 and ACE2, to other host cells making them more susceptible to the infection after exosomes endocytosis (96). Our functional analysis revealed down-regulation of some pathways related to cell communication, adhesion and signal transduction in mild cases, probably as a mechanism to minimize viral spread. Conversely, severe cases showed up-regulation of genes involved in cell junction, adhesion and extracellular exosomes, all processes that could facilitate the viral dissemination. CD59 is a non-IFN inducible protein that acts as inhibitor of complement-mediated lysis by disruption of the membrane attack complex. Its implication in viral complement evasion of many enveloped viruses (97), including coronaviruses (98), has been demonstrated. This escape mechanism can be mediated by incorporating CD59 to the viral envelope, controlling cellular CD59 expression or producing a counterpart protein that mimics host CD59, or even use it as cell entry receptor (99). Overall, immune response in buccal cavity, due to its anatomical location and interplay between upper and lower respiratory tract (LRT), could play a key role in viral spread and transmission not only at inter-individual level (37) but also in intra-individual viral dissemination to LRT, leading to a more severe phenotype.

Transversal analysis of different tissue samples from the same patients showed a tissue-specific behaviour depending on illness severity. Most of the tissue-specific severity signals observed were represented by genes involved in innate immune response and cytokine signaling. Genes that appeared differentially expressed with severities in nasal and blood samples are related to leukocyte chemotaxis, cytokine-cytokine signaling, cell adhesion and extracellular exosomes, with higher expression of severe cases (with respect to those with mild disease) in blood and, in contrast, lower expression in nasal samples; B-cell antigen receptor signaling genes showed an opposite pattern. Specific differential signatures of severity in saliva samples correspond to genes involved in inflammatory and NK activation routes, cell adhesion, and vesicle trafficking.

Our results point to a set of genes and pathways that are tissue-specific and correlated to severity (Fig 5); many of these are involved in the innate immune system and cytokine/chemokine signaling pathways. Disease severity at systemic level might be strongly dependent on local immune response. Thus, the nasal epithelium shows a robust activation of the antiviral immune machinery in mild cases, characterized by IFN pathway involvement, thus triggering the IFN induced genes, chemokine release and transduction signaling in the cells, leading to the activation of downstream cascades related to Th1, NK and T-cytotoxic adaptative immune response. Instead, severe cases showed these local antiviral reactions switched off; they develop an exacerbated innate response represented by an over-expression of genes related to monocyte-macrophage recruitment in nasal epithelium (Fig 5). We did not find signals of immune viral response in buccal mucosa in mild patients, which probably indicates a successful local control of the infection and, therefore, prevention of viral dissemination and systemic colonization. On the contrary, buccal mucosa in moderate and, to a greater extent, severe cases, provided evident signals of viral activity, cell arresting and viral dissemination to the LRT (Fig 5); ultimately generating an exacerbated innate and impaired adaptative systemic immune responses. In addition, the buccal cavity might play a key role in body infection dissemination: a severe phenotype seems to be associated to limited immunological resistance to the virus in the mouth, therefore facilitating the dissemination of the virus towards the LRT. This could make saliva not only a good prognostic proxy of severity, but also a source of therapeutic targets to contain viral dissemination in early stages of the disease.

**Fig 5.**
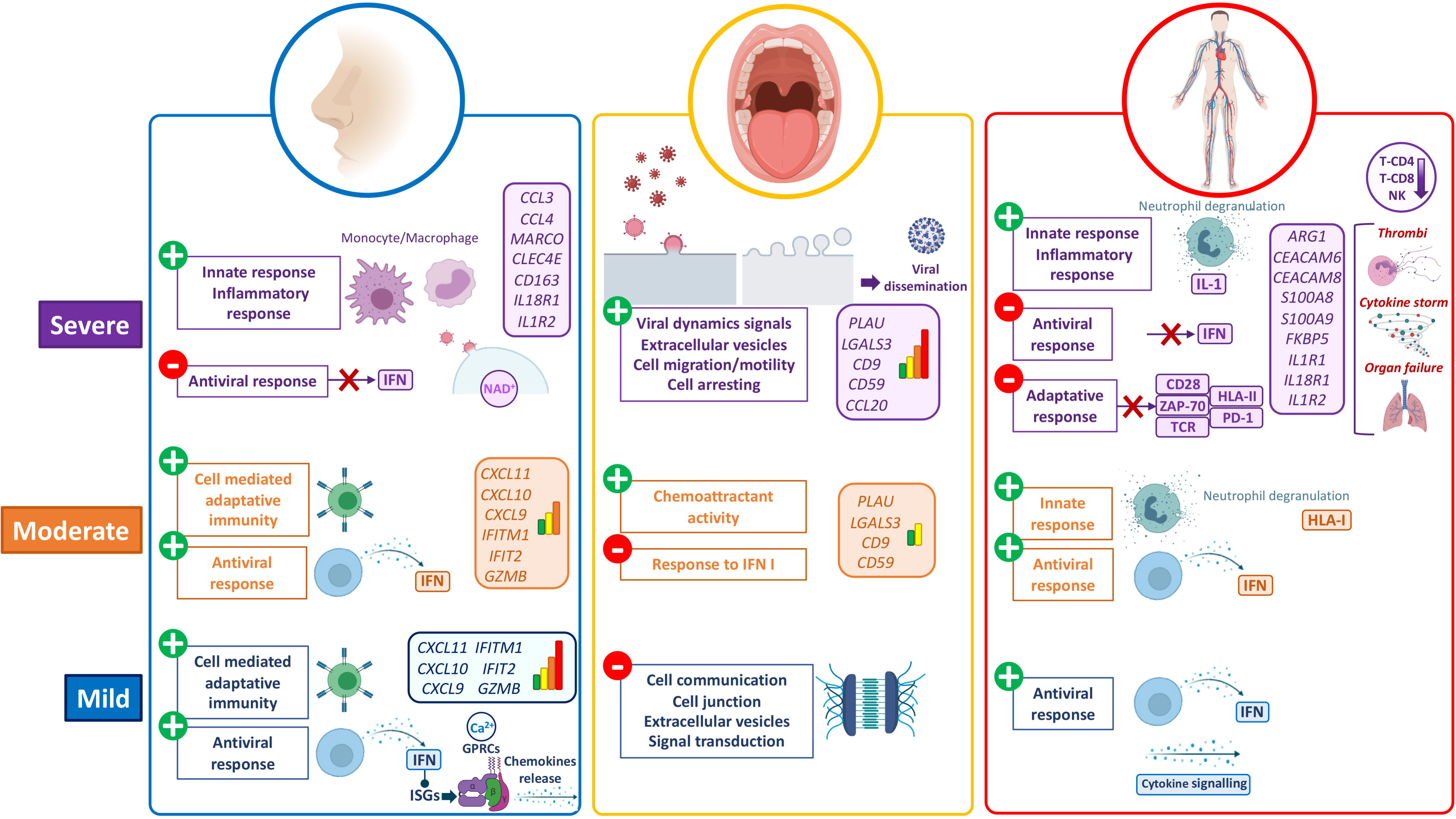
Schematic representation of the main findings in gene expression and pathway analysis of COVID-19 severity in nasal, oral and blood tissues. Fig was built with Biorender (https://biorender.com/).

Overall, our findings include transcript biomarkers and pathways that signal severity at different tissue layers, and might illuminate new diagnostic, prognostic or therapeutic targets in COVID-19. Further validation using additional cohorts should be warranted.

## Acknowledgements

This study received support from projects: GePEM (Instituto de Salud Carlos III(ISCIII)/PI16/01478/Cofinanciado FEDER), DIAVIR (Instituto de Salud Carlos III(ISCIII)/DTS19/00049/Cofinanciado FEDER; Proyecto de Desarrollo Tecnológico en Salud), Resvi-Omics (Instituto de Salud Carlos III(ISCIII)/PI19/01039/Cofinanciado FEDER), BI-BACVIR (PRIS-3; Agencia de Conocimiento en Salud (ACIS)—Servicio Gallego de Salud (SERGAS)—Xunta de Galicia; Spain), Programa Traslaciona Covid-19 (ACIS—Servicio Gallego de Salud (SERGAS)—Xunta de Galicia; Spain) and Axencia Galega de Innovación (GAIN; IN607B 2020/08—Xunta de Galicia; Spain) awarded to A.S.; and projects ReSVinext (Instituto de Salud Carlos III(ISCIII)/PI16/01569/Cofinanciado FEDER), Enterogen (Instituto de Salud Carlos III(ISCIII)/ PI19/01090/Cofinanciado FEDER), and Axencia Galega de Innovación (GAIN; IN845D 2020/23—Xunta de Galicia; Spain) awarded to F.M.-T. We also thank Aida Freire Valls from Nanostring for her support.

## Author Contribution

AS and FMT conceived, designed, and provided financial support to the study. IRC, JGR, CRV, NRN, GBC, HPF, MLC, ADU, CRT, SP, MJCT, SVL and FMT were involved in sample recruitment and analysis of the clinical data. AGC carried out the experimental lab work. AS, AGC, JPS, XB, RBA analyzed the data. AGC and AS wrote the initial draft of the article. All the authors revised and contributed to the final version of the manuscript.

## Appendix S1 figures legends

**S1 Fig.** A) Analysis of qPCR results in nasopharyngeal samples. Correlation analysis of Ct (cycle threshold) values of viral genes and days of symptoms showing Pearson correlation index (R). Boxplots of Ct values of viral genes stratified by disease severity. *P*-values are also shown (p). B) Immune cell fractions estimated from gene expression data.

**S2 Fig.** Boxplots of normalized expression data from DEGs stratified by severity between COVID-19 patients and controls in blood samples. *P*-values of statistical comparisons are also indicated.

**S3 Fig.** nCounter and RNA-seq/microarray data correlation analysis of log_2_FC values obtained from the comparisons. A) COVID-19 cases *vs.* controls; B) COVID-19 cases *vs.* other viral and bacterial infections, and C) severe COVID-19 cases *vs*. controls.

**S4 Fig.** Volcano plots showing the DEGs between different severity comparisons and also healthy controls in blood samples.

**S5 Fig.** Boxplots of normalized expression data stratified by severity from DEGs between severe cases and all remaining categories in blood samples. *P*-values of statistical comparisons are also indicated.

**S6 Fig.** Volcano plots showing the DEGs from the hospitalized *vs.* controls and *vs.* non-hospitalized comparisons in blood samples. Boxplots of normalized expression showing the most relevant genes from hospitalized *vs.* non-hospitalized comparison.

**S7 Fig.** Dotplot and concept network plot from GSEA pathway analysis showing results of severe cases *vs*. healthy controls in blood samples using Reactome and GO database as reference.

**S8 Fig.** Dotplot and concept network plot from GSEA pathway analysis showing results of severe *vs*. moderate cases in blood samples using Reactome and GO database as reference.

**S9 Fig.** Dotplot and concept network plot from GSEA pathway analysis showing results of severe *vs*. mild cases in blood samples using Reactome and GO database as reference.

**S10 Fig.** Dotplot, GSEA plots and concept network plots from GSEA pathway analysis showing results of mild *vs*. controls, moderate *vs.* controls and moderate *vs.* mild cases in blood samples using Reactome and GO database as reference.

**S11 Fig.** Dotplots, GSEA plots and concept network plots from GSEA and ORA pathway analysis showing results of hospitalized vs. non-hospitalized cases in blood samples using Reactome and GO database as reference.

**S12 Fig.** Boxplots of normalized expression data from DEGs stratified by severity between COVID-19 cases and controls and hospitalized *vs*. non-hospitalized patients in nasal epithelium samples. *P*-values of statistical comparisons are also indicated.

**S13 Fig.** Volcano plots showing the DEGs between different severity comparisons, and healthy controls in nasal epithelium samples.

**S14 Fig.** Boxplots of normalized expression data from DEGs overlapping between moderate and mild cases *vs*. controls independently in nasal epithelium samples.

**S15 Fig.** Boxplots of normalized expression data from seven out of the eight DEGs overlapping between comparisons of severe cases and all remaining categories independently in nasal epithelium samples.

**S16 Fig.** Dotplots and concept network plots from GSEA and ORA pathway analysis showing results of moderate cases *vs.* controls in nasal epithelium samples using GO database as reference.

**S17 Fig.** Dotplots and concept network plots from ORA pathway analysis showing results of mild cases *vs.* controls in nasal epithelium samples using GO database as reference.

**S18 Fig.** Dotplots, concept network plots and ridge plot from GSEA and ORA pathway analysis showing results of severe *vs.* mild cases in nasal epithelium samples using Reactome and GO database as reference.

**S19 Fig.** Dotplots and GSEA plots from GSEA pathway analysis showing results of severe vs. moderate cases in in nasal epithelium samples using Reactome and GO database as reference.

**S20 Fig.** Dotplot, concept network plots and ridge plot from GSEA and ORA pathway analysis showing results of severe *vs.* mild cases in nasal epithelium samples using GO database as reference.

**S21 Fig.** Boxplots of normalized expression data from DEGs between hospitalized *vs*. non-hospitalized cases in saliva samples. *P*-values of statistical comparisons are also indicated.

**S22 Fig.** Boxplots of normalized expression data stratified by severity from DEGs between hospitalized *vs*. non-hospitalized cases in saliva samples. *P*-values of statistical comparisons are also indicated.

**S23 Fig.** Volcano plots showing the DEGs between different severity comparisons, and healthy controls in saliva samples.

**S24 Fig.** Boxplots of normalized expression data stratified by severity showing the most relevant DEGs in severe cases with respect to all other categories in saliva samples. *P*-values of statistical comparisons are also indicated.

**S25 Fig.** Concept network plots and ridge plot from GSEA and ORA pathway analysis results of severe cases *vs.* controls in saliva samples using GO database as reference.

**S26 Fig.** Ridge plots from GSEA pathway analysis results of severe *vs.* mild cases in saliva samples using GO database as reference.

**S27 Fig.** Dotplots from GSEA pathway analysis results of severe *vs.* moderate cases in saliva samples using GO database as reference.

**S28 Fig.** Dotplots and concept network plots from GSEA pathway analysis showing results of severe *vs.* mild cases in saliva samples using Reactome and GO databases as reference.

**S29 Fig.** Concept network plots from GSEA pathway analysis showing results of moderate cases *vs.* controls in saliva samples using GO database as reference.

**S30 Fig.** Concept network plots from GSEA pathway analysis showing results of mild cases *vs.* controls in saliva samples using GO database as reference.

**S31 Fig.** Heatmap and clustering analysis of the differentially expressed between severities (severe/mild) and different tissue samples obtained from transversal analysis (thresholds: *P*-adjusted = 0.01, cumulative log_2_FC = |2|). Gene clusters were generated using k-means partitioning. Boxplots of normalized expression data from the most relevant DE between severities and tissues. *P*-values of statistical comparisons are also indicated.

## Supplementary tables legends

**S1 Table.** Description of the samples from different tissues used in the multi-source analysis.

**S2 Table.** DE analysis of immune genes between different COVID categories in blood samples

**S3 Table.** Pathway analysis results obtained from GSEA and ORA in blood samples.

**S4 Table.** DE analysis of immune genes between different COVID categories in nasal epithelium samples

**S5 Table.** Pathway analysis results obtained from GSEA and ORA in nasal epithelium samples.

**S6 Table.** DE analysis of immune genes between different COVID categories in saliva samples

**S7 Table.** Pathway analysis results obtained from GSEA and ORA in saliva samples.

**S8 Table**. Multi-tissue transversal expression analysis of immune genes testing the differential effect of severity comparing different tissues collected in the same patients (severe/mild).

## References

1. Gómez-Carballa A, Bello X, Pardo-Seco J, Martinón-Torres F, Salas A. Mapping genome variation of SARS-CoV-2 worldwide highlights the impact of COVID-19 super-spreaders. Genome Research. 2020;30(10):1434–48.

2. Pipes L, Wang H, Huelsenbeck JP, Nielsen R. Assessing uncertainty in the rooting of the SARS-CoV-2 phylogeny. Mol Biol Evol. 2020.

3. Davies NG, Abbott S, Barnard RC, Jarvis CI, Kucharski AJ, Munday JD, et al. Estimated transmissibility and impact of SARS-CoV-2 lineage B.1.1.7 in England. Science. 2021;372(6538).

4. Gómez-Carballa A, Bello X, Pardo-Seco J, Del Molino MLP, Martinón-Torres F, Salas A. Phylogeography of SARS-CoV-2 pandemic in Spain: a story of multiple introductions, micro-geographic stratification, founder effects, and super-spreaders. Zoological research. 2020;41(6):605.

5. Althouse BM, Wenger EA, Miller JC, Scarpino SV, Allard A, Hebert-Dufresne L, et al. Superspreading events in the transmission dynamics of SARS-CoV-2: Opportunities for interventions and control. PLoS Biol. 2020;18(11):e3000897.

6. Lemieux JE, Siddle KJ, Shaw BM, Loreth C, Schaffner SF, Gladden-Young A, et al. Phylogenetic analysis of SARS-CoV-2 in Boston highlights the impact of superspreading events. Science. 2020.

7. Sungnak W, Huang N, Bécavin C, Berg M, Queen R, Litvinukova M, et al. SARS-CoV-2 entry factors are highly expressed in nasal epithelial cells together with innate immune genes. Nat Med. 2020;26(5):681–7.

8. He X, Lau EHY, Wu P, Deng X, Wang J, Hao X, et al. Temporal dynamics in viral shedding and transmissibility of COVID-19. Nat Med. 2020;26(5):672–5.

9. Zou L, Ruan F, Huang M, Liang L, Huang H, Hong Z, et al. SARS-CoV-2 Viral Load in Upper Respiratory Specimens of Infected Patients. N Engl J Med. 2020;382(12):1177–9.

10. Jin X, Lian JS, Hu JH, Gao J, Zheng L, Zhang YM, et al. Epidemiological, clinical and virological characteristics of 74 cases of coronavirus-infected disease 2019 (COVID-19) with gastrointestinal symptoms. Gut. 2020;69(6):1002–9.

11. Lamers MM, Beumer J, van der Vaart J, Knoops K, Puschhof J, Breugem TI, et al. SARS-CoV-2 productively infects human gut enterocytes. Science. 2020;369(6499):50–4.

12. Huang C, Wang Y, Li X, Ren L, Zhao J, Hu Y, et al. Clinical features of patients infected with 2019 novel coronavirus in Wuhan, China. Lancet. 2020;395(10223):497–506.

13. Qin C, Zhou L, Hu Z, Zhang S, Yang S, Tao Y, et al. Dysregulation of Immune Response in Patients With Coronavirus 2019 (COVID-19) in Wuhan, China. Clin Infect Dis. 2020;71(15):762–8.

14. Yang X, Yu Y, Xu J, Shu H, Xia J, Liu H, et al. Clinical course and outcomes of critically ill patients with SARS-CoV-2 pneumonia in Wuhan, China: a single-centered, retrospective, observational study. Lancet Respir Med. 2020;8(5):475–81.

15. Zheng HY, Zhang M, Yang CX, Zhang N, Wang XC, Yang XP, et al. Elevated exhaustion levels and reduced functional diversity of T cells in peripheral blood may predict severe progression in COVID-19 patients. Cell Mol Immunol. 2020;17(5):541–3.

16. Gomez-Rial J, Rivero-Calle I, Salas A, Martinon-Torres F. Role of Monocytes/Macrophages in Covid-19 Pathogenesis: Implications for Therapy. Infect Drug Resist. 2020;13:2485–93.

17. Gómez-Rial J, Currás-Tuala MJ, Rivero-Calle I, Gómez-Carballa A, Cebey-López M, Rodríguez-Tenreiro C, et al. Increased Serum Levels of sCD14 and sCD163 Indicate a Preponderant Role for Monocytes in COVID-19 Immunopathology. Frontiers in Immunology. 2020;11:2436.

18. Zhang C, Wu Z, Li JW, Zhao H, Wang GQ. Cytokine release syndrome in severe COVID-19: interleukin-6 receptor antagonist tocilizumab may be the key to reduce mortality. Int J Antimicrob Agents. 2020;55(5):105954.

19. McGonagle D, Sharif K, O’Regan A, Bridgewood C. The Role of Cytokines including Interleukin-6 in COVID-19 induced Pneumonia and Macrophage Activation Syndrome-Like Disease. Autoimmun Rev. 2020;19(6):102537.

20. Zhou Y, Zhang J, Wang D, Guan W, Qin J, Xu X, et al. Profiling of the immune repertoire in COVID-19 patients with mild, severe, convalescent, or retesting-positive status. J Autoimmun. 2021;118:102596.

21. Zheng HY, Xu M, Yang CX, Tian RR, Zhang M, Li JJ, et al. Longitudinal transcriptome analyses show robust T cell immunity during recovery from COVID-19. Signal Transduct Target Ther. 2020;5(1):294.

22. Ong EZ, Chan YFZ, Leong WY, Lee NMY, Kalimuddin S, Haja Mohideen SM, et al. A Dynamic Immune Response Shapes COVID-19 Progression. Cell Host Microbe. 2020;27(6):879–82.e2.

23. Thair SA, He YD, Hasin-Brumshtein Y, Sakaram S, Pandya R, Toh J, et al. Transcriptomic similarities and differences in host response between SARS-CoV-2 and other viral infections. iScience. 2021;24(1):101947.

24. Aschenbrenner AC, Mouktaroudi M, Krämer B, Oestreich M, Antonakos N, Nuesch-Germano M, et al. Disease severity-specific neutrophil signatures in blood transcriptomes stratify COVID-19 patients. Genome Med. 2021;13(1):7.

25. Alfi O, Yakirevitch A, Wald O, Wandel O, Izhar U, Oiknine-Djian E, et al. Human nasal and lung tissues infected *ex vivo* with SARS-CoV-2 provide insights into differential tissue-specific and virus-specific innate immune responses in the upper and lower respiratory tract. J Virol. 2021.

26. Gamage AM, Tan KS, Chan WOY, Liu J, Tan CW, Ong YK, et al. Infection of human Nasal Epithelial Cells with SARS-CoV-2 and a 382-nt deletion isolate lacking ORF8 reveals similar viral kinetics and host transcriptional profiles. PLoS Pathog. 2020;16(12):e1009130.

27. Sajuthi SP, DeFord P, Li Y, Jackson ND, Montgomery MT, Everman JL, et al. Type 2 and interferon inflammation regulate SARS-CoV-2 entry factor expression in the airway epithelium. Nat Commun. 2020;11(1):5139.

28. Lieberman NAP, Peddu V, Xie H, Shrestha L, Huang ML, Mears MC, et al. In vivo antiviral host transcriptional response to SARS-CoV-2 by viral load, sex, and age. PLoS Biol. 2020;18(9):e3000849.

29. Pierce CA, Sy S, Galen B, Goldstein DY, Orner E, Keller MJ, et al. Natural mucosal barriers and COVID-19 in children. JCI Insight. 2021;6(9).

30. Jain R, Ramaswamy S, Harilal D, Uddin M, Loney T, Nowotny N, et al. Host transcriptomic profiling of COVID-19 patients with mild, moderate, and severe clinical outcomes. Comput Struct Biotechnol J. 2021;19:153–60.

31. Pasomsub E, Watcharananan SP, Boonyawat K, Janchompoo P, Wongtabtim G, Suksuwan W, et al. Saliva sample as a non-invasive specimen for the diagnosis of coronavirus disease 2019: a cross-sectional study. Clin Microbiol Infect. 2020.

32. To KK, Tsang OT, Yip CC, Chan KH, Wu TC, Chan JM, et al. Consistent Detection of 2019 Novel Coronavirus in Saliva. Clin Infect Dis. 2020;71(15):841–3.

33. Wyllie AL, Fournier J, Casanovas-Massana A, Campbell M, Tokuyama M, Vijayakumar P, et al. Saliva or Nasopharyngeal Swab Specimens for Detection of SARS-CoV-2. N Engl J Med. 2020;383(13):1283–6.

34. Teo AKJ, Choudhury Y, Tan IB, Cher CY, Chew SH, Wan ZY, et al. Saliva is more sensitive than nasopharyngeal or nasal swabs for diagnosis of asymptomatic and mild COVID-19 infection. Sci Rep. 2021;11(1):3134.

35. Zupin L, Pascolo L, Crovella S. Is *FURIN* gene expression in salivary glands related to SARS-CoV-2 infectivity through saliva? J Clin Pathol. 2020.

36. Chen L, Zhao J, Peng J, Li X, Deng X, Geng Z, et al. Detection of SARS-CoV-2 in saliva and characterization of oral symptoms in COVID-19 patients. Cell Prolif. 2020;53(12):e12923.

37. Huang N, Pérez P, Kato T, Mikami Y, Okuda K, Gilmore RC, et al. SARS-CoV-2 infection of the oral cavity and saliva. Nat Med. 2021.

38. Vandesompele J, De Preter K, Pattyn F, Poppe B, Van Roy N, De Paepe A, et al. Accurate normalization of real-time quantitative RT-PCR data by geometric averaging of multiple internal control genes. Genome biology. 2002;3(7):RESEARCH0034.

39. Zhong S. ctrlGene: Assess the Stability of Candidate Housekeeping Genes. version 101, R package. 2019.

40. Love MI, Huber W, Anders S. Moderated estimation of fold change and dispersion for RNA-seq data with DESeq2. Genome Biol. 2014;15(12):550.

41. Risso D, Ngai J, Speed TP, Dudoit S. Normalization of RNA-seq data using factor analysis of control genes or samples. Nat Biotechnol. 2014;32(9):896–902.

42. Bhattacharya A, Hamilton AM, Furberg H, Pietzak E, Purdue MP, Troester MA, et al. An approach for normalization and quality control for NanoString RNA expression data. Brief Bioinform. 2020.

43. Blighe K, Rana S, Lewis M. EnhancedVolcano: Publication-ready volcano plots with enhanced colouring and labeling. version 180, R package. 2020.

44. Gu Z, Eils R, Schlesner M. Complex heatmaps reveal patterns and correlations in multidimensional genomic data. Bioinformatics. 2016;32(18):2847–9.

45. Wickham H. ggplot2: Elegant Graphics for Data Analysis: Springer-Verlag New York; 2016.

46. Subramanian A, Tamayo P, Mootha VK, Mukherjee S, Ebert BL, Gillette MA, et al. Gene set enrichment analysis: a knowledge-based approach for interpreting genome-wide expression profiles. Proc Natl Acad Sci U S A. 2005;102(43):15545–50.

47. Yu G, Wang LG, Han Y, He QY. clusterProfiler: an R package for comparing biological themes among gene clusters. Omics : a journal of integrative biology. 2012;16(5):284–7.

48. Yu G. enrichplot: Visualization of Functional Enrichment Result. 2021.

49. Mahajan P, Kuppermann N, Mejias A, Suarez N, Chaussabel D, Casper TC, et al. Association of RNA Biosignatures With Bacterial Infections in Febrile Infants Aged 60 Days or Younger. JAMA. 2016;316(8):846–57.

50. Parnell GP, McLean AS, Booth DR, Armstrong NJ, Nalos M, Huang SJ, et al. A distinct influenza infection signature in the blood transcriptome of patients with severe community-acquired pneumonia. Crit Care. 2012;16(4):R157.

51. Herberg JA, Kaforou M, Gormley S, Sumner ER, Patel S, Jones KD, et al. Transcriptomic profiling in childhood H1N1/09 influenza reveals reduced expression of protein synthesis genes. The Journal of infectious diseases. 2013;208(10):1664–8.

52. Suarez NM, Bunsow E, Falsey AR, Walsh EE, Mejias A, Ramilo O. Superiority of transcriptional profiling over procalcitonin for distinguishing bacterial from viral lower respiratory tract infections in hospitalized adults. J Infect Dis. 2015;212(2):213–22.

53. Herberg JA, Kaforou M, Wright VJ, Shailes H, Eleftherohorinou H, Hoggart CJ, et al. Diagnostic test accuracy of a 2-transcript host RNA signature for discriminating bacterial vs viral infection in febrile children. JAMA. 2016;316(8):835–45.

54. DeBerg HA, Zaidi MB, Altman MC, Khaenam P, Gersuk VH, Campos FD, et al. Shared and organism-specific host responses to childhood diarrheal diseases revealed by whole blood transcript profiling. PLoS One. 2018;13(1):e0192082.

55. Du P, Kibbe WA, Lin SM. lumi: a pipeline for processing Illumina microarray. Bioinformatics. 2008;24(13):1547–8.

56. Carvalho BS, Irizarry RA. A framework for oligonucleotide microarray preprocessing. Bioinformatics. 2010;26(19):2363–7.

57. Barral-Arca R, Pardo-Seco J, Martinón-Torres F, Salas A. A 2-transcript host cell signature distinguishes viral from bacterial diarrhea and it is influenced by the severity of symptoms. Sci Rep. 2018;8(1):8043.

58. Sweeney TE, Wong HR, Khatri P. Robust classification of bacterial and viral infections via integrated host gene expression diagnostics. Sci Transl Med. 2016;8(346):346ra91.

59. Ritchie ME, Phipson B, Wu D, Hu Y, Law CW, Shi W, et al. limma powers differential expression analyses for RNA-sequencing and microarray studies. Nucleic Acids Res. 2015;43(7):e47.

60. Newman AM, Steen CB, Liu CL, Gentles AJ, Chaudhuri AA, Scherer F, et al. Determining cell type abundance and expression from bulk tissues with digital cytometry. Nat Biotechnol. 2019;37(7):773–82.

61. Newman AM, Liu CL, Green MR, Gentles AJ, Feng W, Xu Y, et al. Robust enumeration of cell subsets from tissue expression profiles. Nat Methods. 2015;12(5):453–7.

62. Crouse J, Kalinke U, Oxenius A. Regulation of antiviral T cell responses by type I interferons. Nat Rev Immunol. 2015;15(4):231–42.

63. Lokugamage KG, Hage A, de Vries M, Valero-Jimenez AM, Schindewolf C, Dittmann M, et al. Type I Interferon Susceptibility Distinguishes SARS-CoV-2 from SARS-CoV. J Virol. 2020;94(23).

64. Mantlo E, Bukreyeva N, Maruyama J, Paessler S, Huang C. Antiviral activities of type I interferons to SARS-CoV-2 infection. Antiviral Res. 2020;179:104811.

65. Contoli M, Papi A, Tomassetti L, Rizzo P, Vieceli Dalla Sega F, Fortini F, et al. Blood Interferon-α Levels and Severity, Outcomes, and Inflammatory Profiles in Hospitalized COVID-19 Patients. Front Immunol. 2021;12:648004.

66. Hadjadj J, Yatim N, Barnabei L, Corneau A, Boussier J, Smith N, et al. Impaired type I interferon activity and inflammatory responses in severe COVID-19 patients. Science. 2020;369(6504):718–24.

67. Leppkes M, Knopf J, Naschberger E, Lindemann A, Singh J, Herrmann I, et al. Vascular occlusion by neutrophil extracellular traps in COVID-19. EBioMedicine. 2020;58:102925.

68. Veras FP, Pontelli MC, Silva CM, Toller-Kawahisa JE, de Lima M, Nascimento DC, et al. SARS-CoV-2-triggered neutrophil extracellular traps mediate COVID-19 pathology. J Exp Med. 2020;217(12).

69. Twaddell SH, Baines KJ, Grainge C, Gibson PG. The Emerging Role of Neutrophil Extracellular Traps in Respiratory Disease. Chest. 2019;156(4):774–82.

70. Nchioua R, Kmiec D, Müller JA, Conzelmann C, Groß R, Swanson CM, et al. SARS-CoV-2 Is Restricted by Zinc Finger Antiviral Protein despite Preadaptation to the Low-CpG Environment in Humans. mBio. 2020;11(5).

71. Xu G, Qi F, Li H, Yang Q, Wang H, Wang X, et al. The differential immune responses to COVID-19 in peripheral and lung revealed by single-cell RNA sequencing. Cell Discov. 2020;6:73.

72. Italiani P, Mosca E, Della Camera G, Melillo D, Migliorini P, Milanesi L, et al. Profiling the Course of Resolving vs. Persistent Inflammation in Human Monocytes: The Role of IL-1 Family Molecules. Front Immunol. 2020;11:1426.

73. Mesquita I, Varela P, Belinha A, Gaifem J, Laforge M, Vergnes B, et al. Exploring NAD+ metabolism in host-pathogen interactions. Cell Mol Life Sci. 2016;73(6):1225–36.

74. Merad M, Martin JC. Pathological inflammation in patients with COVID-19: a key role for monocytes and macrophages. Nat Rev Immunol. 2020;20(6):355–62.

75. Chua RL, Lukassen S, Trump S, Hennig BP, Wendisch D, Pott F, et al. COVID-19 severity correlates with airway epithelium-immune cell interactions identified by single-cell analysis. Nat Biotechnol. 2020;38(8):970–9.

76. Landry ML, Foxman EF. Antiviral Response in the Nasopharynx Identifies Patients With Respiratory Virus Infection. J Infect Dis. 2018;217(6):897–905.

77. Buchrieser J, Dufloo J, Hubert M, Monel B, Planas D, Rajah MM, et al. Syncytia formation by SARS-CoV-2-infected cells. EMBO J. 2020;39(23):e106267.

78. Wang WH, Lin CY, Chang MR, Urbina AN, Assavalapsakul W, Thitithanyanont A, et al. The role of galectins in virus infection - A systemic literature review. J Microbiol Immunol Infect. 2020;53(6):925–35.

79. Liu FT, Hsu DK. The role of galectin-3 in promotion of the inflammatory response. Drug News Perspect. 2007;20(7):455–60.

80. Chen YJ, Wang SF, Weng IC, Hong MH, Lo TH, Jan JT, et al. Galectin-3 Enhances Avian H5N1 Influenza A Virus-Induced Pulmonary Inflammation by Promoting NLRP3 Inflammasome Activation. Am J Pathol. 2018;188(4):1031–42.

81. Caniglia JL, Asuthkar S, Tsung AJ, Guda MR, Velpula KK. Immunopathology of galectin-3: an increasingly promising target in COVID-19. F1000Res. 2020;9:1078.

82. Garcia-Revilla J, Deierborg T, Venero JL, Boza-Serrano A. Hyperinflammation and Fibrosis in Severe COVID-19 Patients: Galectin-3, a Target Molecule to Consider. Front Immunol. 2020;11:2069.

83. Millet JK, Whittaker GR. Host cell proteases: Critical determinants of coronavirus tropism and pathogenesis. Virus Res. 2015;202:120–34.

84. Devaux CA, Rolain JM, Raoult D. ACE2 receptor polymorphism: Susceptibility to SARS-CoV-2, hypertension, multi-organ failure, and COVID-19 disease outcome. J Microbiol Immunol Infect. 2020;53(3):425–35.

85. Berri F, Rimmelzwaan GF, Hanss M, Albina E, Foucault-Grunenwald ML, Lê VB, et al. Plasminogen controls inflammation and pathogenesis of influenza virus infections via fibrinolysis. PLoS Pathog. 2013;9(3):e1003229.

86. Goto H, Wells K, Takada A, Kawaoka Y. Plasminogen-binding activity of neuraminidase determines the pathogenicity of influenza A virus. J Virol. 2001;75(19):9297–301.

87. Dubovi EJ, Geratz JD, Tidwell RR. Enhancement of respiratory syncytial virus-induced cytopathology by trypsin, thrombin, and plasmin. Infect Immun. 1983;40(1):351–8.

88. Kam YW, Okumura Y, Kido H, Ng LF, Bruzzone R, Altmeyer R. Cleavage of the SARS coronavirus spike glycoprotein by airway proteases enhances virus entry into human bronchial epithelial cells in vitro. PLoS One. 2009;4(11):e7870.

89. Ji HL, Zhao R, Matalon S, Matthay MA. Elevated Plasmin(ogen) as a Common Risk Factor for COVID-19 Susceptibility. Physiol Rev. 2020;100(3):1065–75.

90. Gralinski LE, Bankhead A, Jeng S, Menachery VD, Proll S, Belisle SE, et al. Mechanisms of severe acute respiratory syndrome coronavirus-induced acute lung injury. mBio. 2013;4(4).

91. D’Alonzo D, De Fenza M, Pavone V. COVID-19 and pneumonia: a role for the uPA/uPAR system. Drug Discov Today. 2020;25(8):1528–34.

92. Kwaan HC, Lindholm PF. The Central Role of Fibrinolytic Response in COVID-19-A Hematologist’s Perspective. Int J Mol Sci. 2021;22(3).

93. Hou Y, Ding Y, Nie H, Ji HL. Fibrinolysis influences SARS-CoV-2 infection in ciliated cells. bioRxiv. 2021.

94. Martin F, Roth DM, Jans DA, Pouton CW, Partridge LJ, Monk PN, et al. Tetraspanins in viral infections: a fundamental role in viral biology? J Virol. 2005;79(17):10839–51.

95. Earnest JT, Hantak MP, Li K, McCray PB, Perlman S, Gallagher T. The tetraspanin CD9 facilitates MERS-coronavirus entry by scaffolding host cell receptors and proteases. PLoS Pathog. 2017;13(7):e1006546.

96. Hassanpour M, Rezaie J, Nouri M, Panahi Y. The role of extracellular vesicles in COVID-19 virus infection. Infect Genet Evol. 2020;85:104422.

97. Li Y, Parks GD. Relative Contribution of Cellular Complement Inhibitors CD59, CD46, and CD55 to Parainfluenza Virus 5 Inhibition of Complement-Mediated Neutralization. Viruses. 2018;10(5).

98. Wei Y, Ji Y, Guo H, Zhi X, Han S, Zhang Y, et al. CD59 association with infectious bronchitis virus particles protects against antibody-dependent complement-mediated lysis. J Gen Virol. 2017;98(11):2725–30.

99. Maloney BE, Perera KD, Saunders DRD, Shadipeni N, Fleming SD. Interactions of viruses and the humoral innate immune response. Clin Immunol. 2020;212:108351.

