## Supplementary material for "A multi-tissue study of immune gene expression profiling highlights the key role of the nasal epithelium in COVID-19 severity": S1 Appendix.pdf

#### Supplementary figures

**Figure S1.** A) Analysis of qPCR results in nasopharyngeal samples. Correlation analysis of Ct (cycle threshold) values of viral genes and days of symptoms showing Pearson correlation index (R). Boxplots of Ct values of viral genes stratified by disease severity. *P*-values are also shown (p). B) Immune cell fractions estimated from gene expression data.

**Figure S2.** Boxplots of normalized expression data from DEGs stratified by severity between COVID-19 patients and controls in blood samples. *P*-values of statistical comparisons are also indicated.

**Figure S3.** nCounter and RNA-seq/microarray data correlation analysis of $log_2FC$ values obtained from the comparisons. A) COVID-19 cases *vs.* controls; B) COVID-19 cases *vs.* other viral and bacterial infections, and C) severe COVID-19 cases *vs*. controls.

**Figure S4.** Volcano plots showing the DEGs between different severity comparisons and also healthy controls in blood samples.

**Figure S5.** Boxplots of normalized expression data stratified by severity from DEGs between severe cases and all remaining categories in blood samples. *P*-values of statistical comparisons are also indicated.

**Figure S6.** Volcano plots showing the DEGs from the hospitalized *vs.* controls and *vs.* non-hospitalized comparisons in blood samples. Boxplots of normalized expression showing the most relevant genes from hospitalized *vs.* non-hospitalized comparison.

**Figure S7.** Dotplot and concept network plot from GSEA pathway analysis showing results of severe cases *vs*. healthy controls in blood samples using Reactome and GO database as reference.

**Figure S8.** Dotplot and concept network plot from GSEA pathway analysis showing results of severe *vs*. moderate cases in blood samples using Reactome and GO database as reference.

**Figure S9.** Dotplot and concept network plot from GSEA pathway analysis showing results of severe *vs*. mild cases in blood samples using Reactome and GO database as reference.

**Figure S10.** Dotplot, GSEA plots and concept network plots from GSEA pathway analysis showing results of mild *vs*. controls, moderate *vs.* controls and moderate *vs.* mild cases in blood samples using Reactome and GO database as reference.

**Figure S11.** Dotplots, GSEA plots and concept network plots from GSEA and ORA pathway analysis showing results of hospitalized vs. non-hospitalized cases in blood samples using Reactome and GO database as reference.

**Figure S12.** Boxplots of normalized expression data from DEGs stratified by severity between COVID-19 cases and controls and hospitalized *vs*. non-hospitalized patients in nasal epithelium samples. *P*-values of statistical comparisons are also indicated.

**Figure S13.** Volcano plots showing the DEGs between different severity comparisons, and healthy controls in nasal epithelium samples.

**Figure S14.** Boxplots of normalized expression data from DEGs overlapping between moderate and mild cases *vs*. controls independently in nasal epithelium samples.

**Figure S15.** Boxplots of normalized expression data from seven out of the eight DEGs overlapping between comparisons of severe cases and all remaining categories independently in nasal epithelium samples.

**Figure S16.** Dotplots and concept network plots from GSEA and ORA pathway analysis showing results of moderate cases *vs.* controls in nasal epithelium samples using GO database as reference.

**Figure S17.** Dotplots and concept network plots from ORA pathway analysis showing results of mild cases *vs.* controls in nasal epithelium samples using GO database as reference.

**Figure S18.** Dotplots, concept network plots and ridge plot from GSEA and ORA pathway analysis showing results of severe *vs.* mild cases in nasal epithelium samples using Reactome and GO database as reference.

**Figure S19.** Dotplots and GSEA plots from GSEA pathway analysis showing results of severe vs. moderate cases in in nasal epithelium samples using Reactome and GO database as reference.

**Figure S20.** Dotplot, concept network plots and ridge plot from GSEA and ORA pathway analysis showing results of severe *vs.* mild cases in nasal epithelium samples using GO database as reference.

**Figure S21.** Boxplots of normalized expression data from DEGs between hospitalized *vs*. non-hospitalized cases in saliva samples. *P*-values of statistical comparisons are also indicated.

**Figure S22.** Boxplots of normalized expression data stratified by severity from DEGs between hospitalized *vs*. non-hospitalized cases in saliva samples. *P*-values of statistical comparisons are also indicated.

**Figure S23.** Volcano plots showing the DEGs between different severity comparisons, and healthy controls in saliva samples.

**Figure S24.** Boxplots of normalized expression data stratified by severity showing the most relevant DEGs in severe cases with respect to all other categories in saliva samples. *P*-values of statistical comparisons are also indicated.

**Figure S25.** Concept network plots and ridge plot from GSEA and ORA pathway analysis results of severe cases *vs.* controls in saliva samples using GO database as reference.

**Figure S26.** Ridge plots from GSEA pathway analysis results of severe *vs.* mild cases in saliva samples using GO database as reference.

**Figure S27.** Dotplots from GSEA pathway analysis results of severe *vs.* moderate cases in saliva samples using GO database as reference.

**Figure S28.** Dotplots and concept network plots from GSEA pathway analysis showing results of severe *vs.* mild cases in saliva samples using Reactome and GO databases as reference.

**Figure S29.** Concept network plots from GSEA pathway analysis showing results of moderate cases *vs.* controls in saliva samples using GO database as reference.

**Figure S30.** Concept network plots from GSEA pathway analysis showing results of mild cases *vs.* controls in saliva samples using GO database as reference.

**Figure S31.** Heatmap and clustering analysis of the differentially expressed between severities (severe/mild) and different tissue samples obtained from transversal analysis (thresholds: *P*-adjusted = 0.01, cumulative $log_2FC = |2|$). Gene clusters were generated using k-means partitioning. Boxplots of normalized expression data from the most relevant DE between severities and tissues. *P*-values of statistical comparisons are also indicated.

#### Supplementary tables

**Table S1.** Description of the samples from different tissues used in the multi-source analysis.

**Table S2.** DE analysis of immune genes between different COVID categories in blood samples

**Table S3.** Pathway analysis results obtained from GSEA and ORA in blood samples.

**Table S4.** DE analysis of immune genes between different COVID categories in nasal epithelium samples

**Table S5.** Pathway analysis results obtained from GSEA and ORA in nasal epithelium samples.

**Table S6.** DE analysis of immune genes between different COVID categories in saliva samples

**Table S7.** Pathway analysis results obtained from GSEA and ORA in saliva samples.

**Table S8**. Multi-tissue transversal expression analysis of immune genes testing the differential effect of severity comparing different tissues collected in the same patients (severe/mild).

### Figure S1.

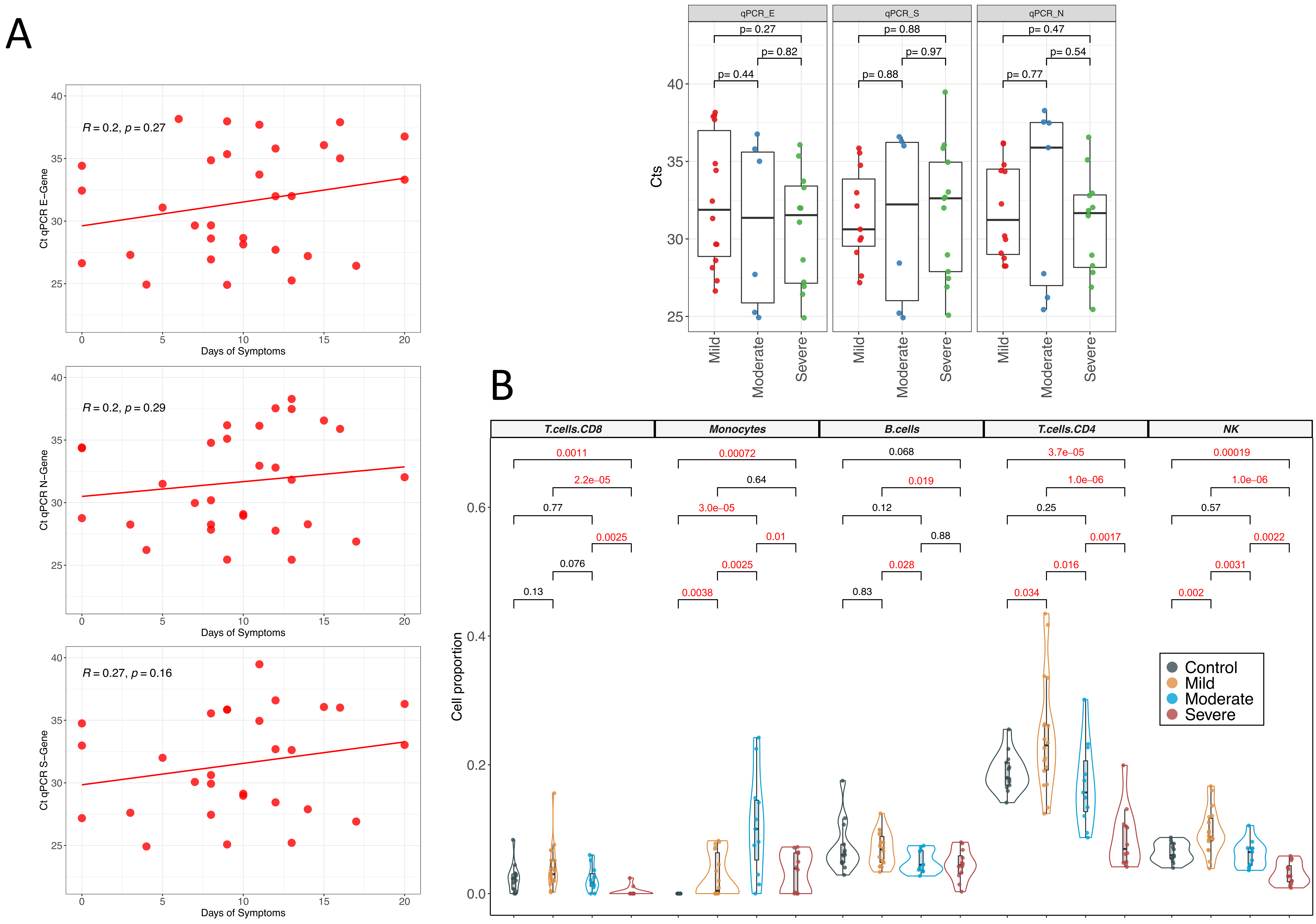
A
R = 0.2, p = 0.27
Ct qPCR E−Gene
Days of Symptoms
R = 0.2, p = 0.29
Ct qPCR N−Gene
Days of Symptoms
R = 0.27, p = 0.16
Ct qPCR S−Gene
Days of Symptoms
qPCR_E
qPCR_S
qPCR_N
p= 0.27
p= 0.82
p= 0.44
p= 0.88
p= 0.97
p= 0.88
p= 0.47
p= 0.54
p= 0.77
Cts
Mild
Moderate
Severe
B
T.cells.CD8
Monocytes
B.cells
T.cells.CD4
NK
Cell proportion
Control
Mild
Moderate
Severe

### Figure S2.

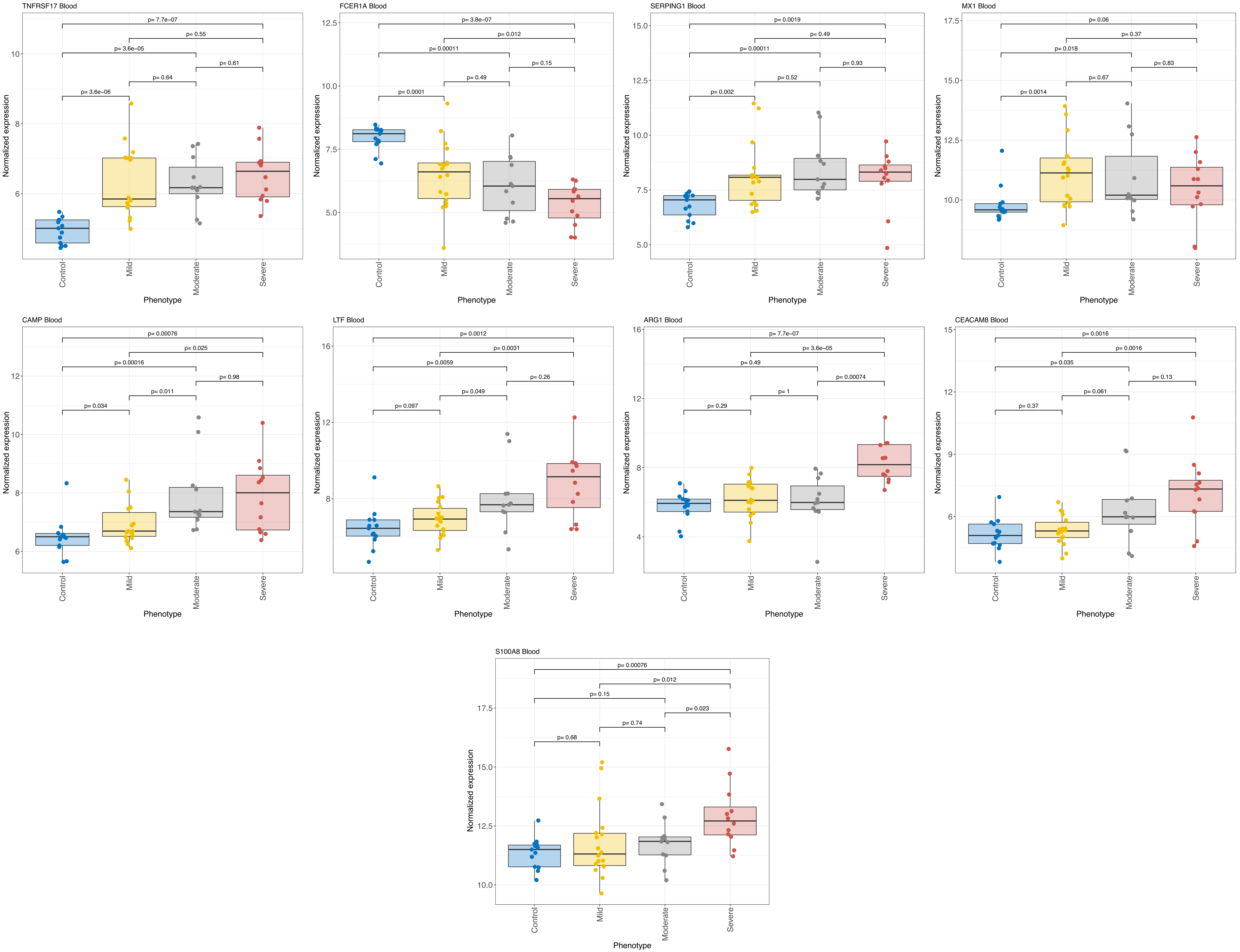

TNFRSF17 Blood
p= 7.7e−07
p= 0.55
p= 3.6e−05
p= 0.61
p= 0.64
p= 3.6e−06
FCER1A Blood
p= 3.8e−07
p= 0.012
p= 0.00011
p= 0.15
p= 0.49
p= 0.0001
SERPING1 Blood
p= 0.0019
p= 0.49
p= 0.00011
p= 0.93
p= 0.52
p= 0.002
MX1 Blood
p= 0.06
p= 0.37
p= 0.018
p= 0.83
p= 0.67
p= 0.0014
CAMP Blood
p= 0.00076
p= 0.025
p= 0.00016
p= 0.98
p= 0.011
p= 0.034
LTF Blood
p= 0.0012
p= 0.0031
p= 0.0059
p= 0.26
p= 0.049
p= 0.097
ARG1 Blood
p= 7.7e−07
p= 3.6e−05
p= 0.49
p= 0.00074
p= 1
p= 0.29
CEACAM8 Blood
p= 0.0016
p= 0.0016
p= 0.035
p= 0.13
p= 0.061
p= 0.37
S100A8 Blood
p= 0.00076
p= 0.012
p= 0.15
p= 0.023
p= 0.74
p= 0.68
Normalized expression
Phenotype
Control
Mild
Moderate
Severe

### Figure S3.

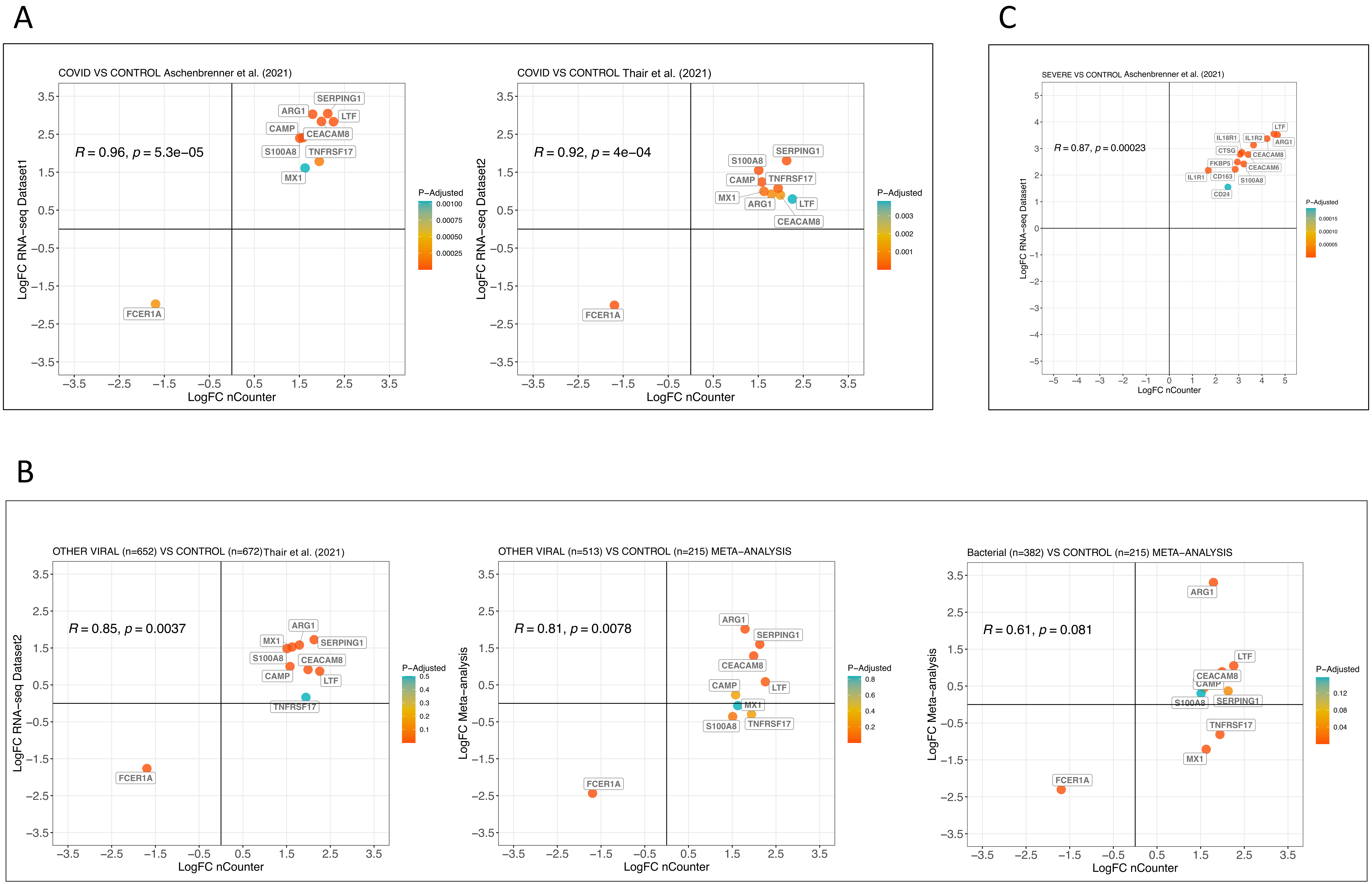

A
COVID VS CONTROL Aschenbrenner et al. (2021)
R = 0.96, p = 5.3e−05
SERPING1
ARG1
LTF
CAMP
CEACAM8
S100A8
TNFRSF17
MX1
FCER1A
LogFC RNA−seq Dataset1
LogFC nCounter
P−Adjusted
0.00100
0.00075
0.00050
0.00025
COVID VS CONTROL Thair et al. (2021)
R = 0.92, p = 4e−04
SERPING1
S100A8
CAMP
TNFRSF17
MX1
ARG1
LTF
CEACAM8
FCER1A
LogFC RNA−seq Dataset2
LogFC nCounter
P−Adjusted
0.003
0.002
0.001
C
SEVERE VS CONTROL Aschenbrenner et al. (2021)
R = 0.87, p = 0.00023
LTF
IL18R1
IL1R2
ARG1
CTSG
CEACAM8
FKBP5
CEACAM6
IL1R1
CD163
S100A8
CD24
LogFC RNA−seq Dataset1
LogFC nCounter
P−Adjusted
0.00015
0.00010
0.00005
B
OTHER VIRAL (n=652) VS CONTROL (n=672)Thair et al. (2021)
R = 0.85, p = 0.0037
ARG1
MX1
SERPING1
S100A8
CEACAM8
CAMP
LTF
TNFRSF17
FCER1A
LogFC RNA−seq Dataset2
LogFC nCounter
P−Adjusted
0.5
0.4
0.3
0.2
0.1
OTHER VIRAL (n=513) VS CONTROL (n=215) META−ANALYSIS
R = 0.81, p = 0.0078
ARG1
SERPING1
CEACAM8
CAMP
LTF
MX1
S100A8
TNFRSF17
FCER1A
LogFC Meta−analysis
LogFC nCounter
P−Adjusted
0.8
0.6
0.4
0.2
Bacterial (n=382) VS CONTROL (n=215) META−ANALYSIS
R = 0.61, p = 0.081
ARG1
LTF
CEACAM8
CAMP
S100A8
SERPING1
TNFRSF17
MX1
FCER1A
LogFC Meta−analysis
LogFC nCounter
P−Adjusted
0.12
0.08
0.04

### Figure S4.

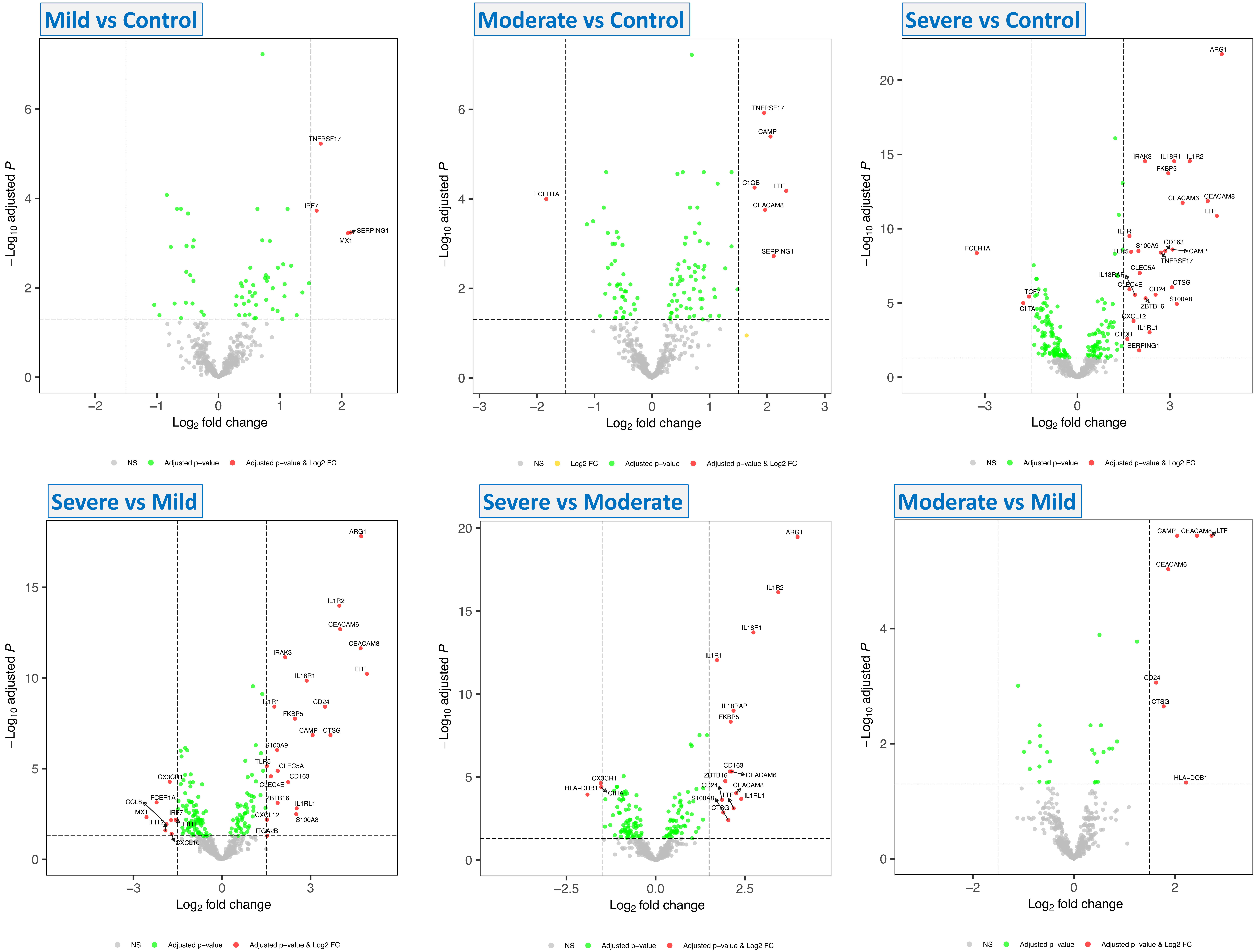
Mild vs Control
Moderate vs Control
Severe vs Control
Severe vs Mild
Severe vs Moderate
Moderate vs Mild
−Log10 adjusted P
Log2 fold change
NS
Log2 FC
Adjusted p−value
Adjusted p−value & Log2 FC
TNFRSF17
IRF7
SERPING1
MX1
FCER1A
CAMP
C1QB
LTF
CEACAM8
ARG1
IRAK3
IL18R1
IL1R2
FKBP5
CEACAM6
IL1R1
S100A9
CD163
TLR5
TCF7
CIITA
IL18RAP
CLEC5A
CLEC4E
CD24
CTSG
ZBTB16
S100A8
CXCL12
IL1RL1
CX3CR1
CCL8
MX1
IFIT2
IFIH1
CXCL10
ITGA2B
HLA−DRB1
HLA−DQB1

### Figure S5.

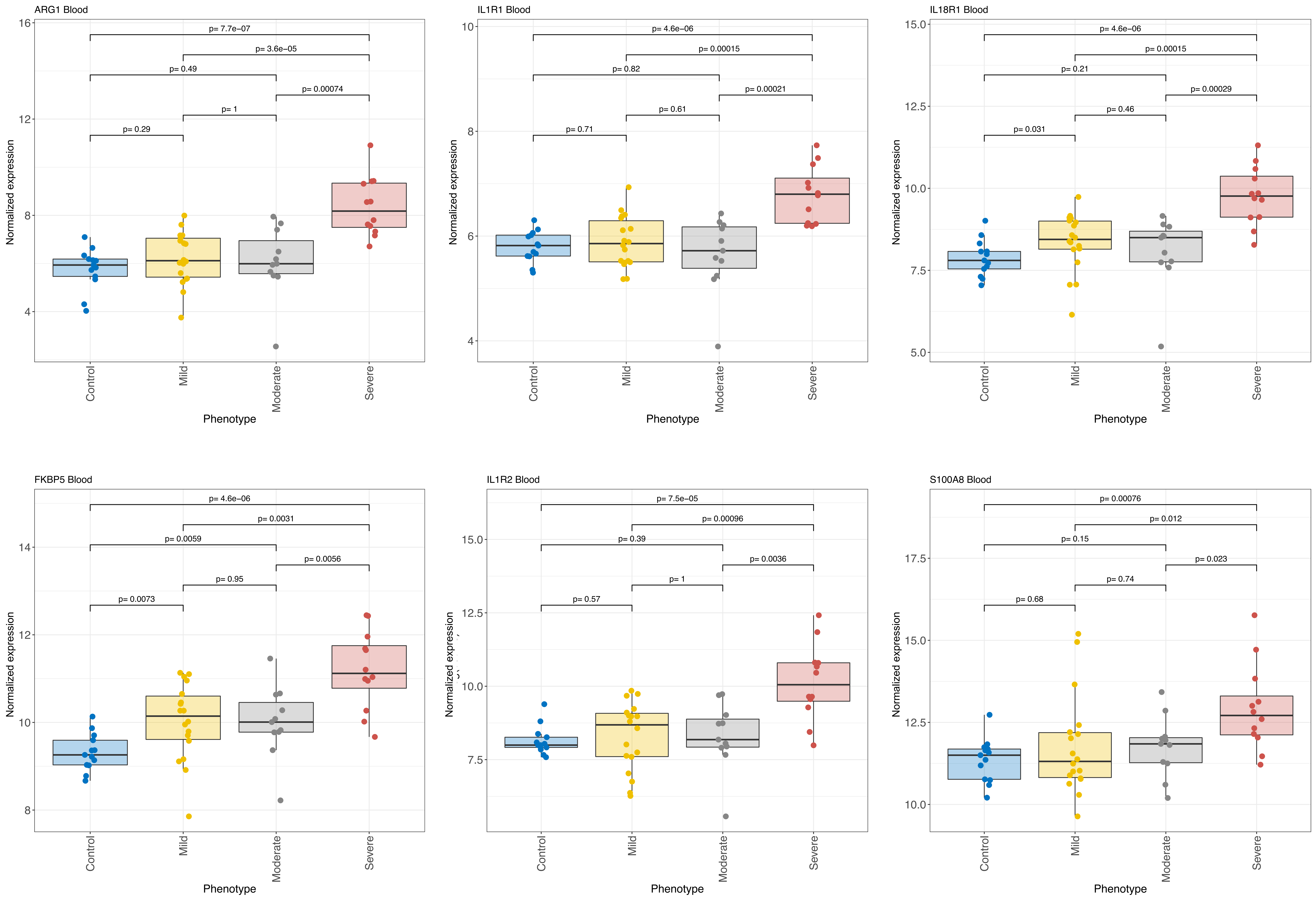
ARG1 Blood
IL1R1 Blood
IL18R1 Blood
FKBP5 Blood
IL1R2 Blood
S100A8 Blood
Normalized expression
Phenotype
Control
Mild
Moderate
Severe
p= 7.7e−07
p= 3.6e−05
p= 0.49
p= 0.00074
p= 1
p= 0.29
p= 4.6e−06
p= 0.00015
p= 0.82
p= 0.00021
p= 0.61
p= 0.71
p= 4.6e−06
p= 0.00015
p= 0.21
p= 0.00029
p= 0.46
p= 0.031
p= 4.6e−06
p= 0.0031
p= 0.0059
p= 0.0056
p= 0.95
p= 0.0073
p= 7.5e−05
p= 0.00096
p= 0.39
p= 0.0036
p= 1
p= 0.57
p= 0.00076
p= 0.012
p= 0.15
p= 0.023
p= 0.74
p= 0.68

### Figure S6.

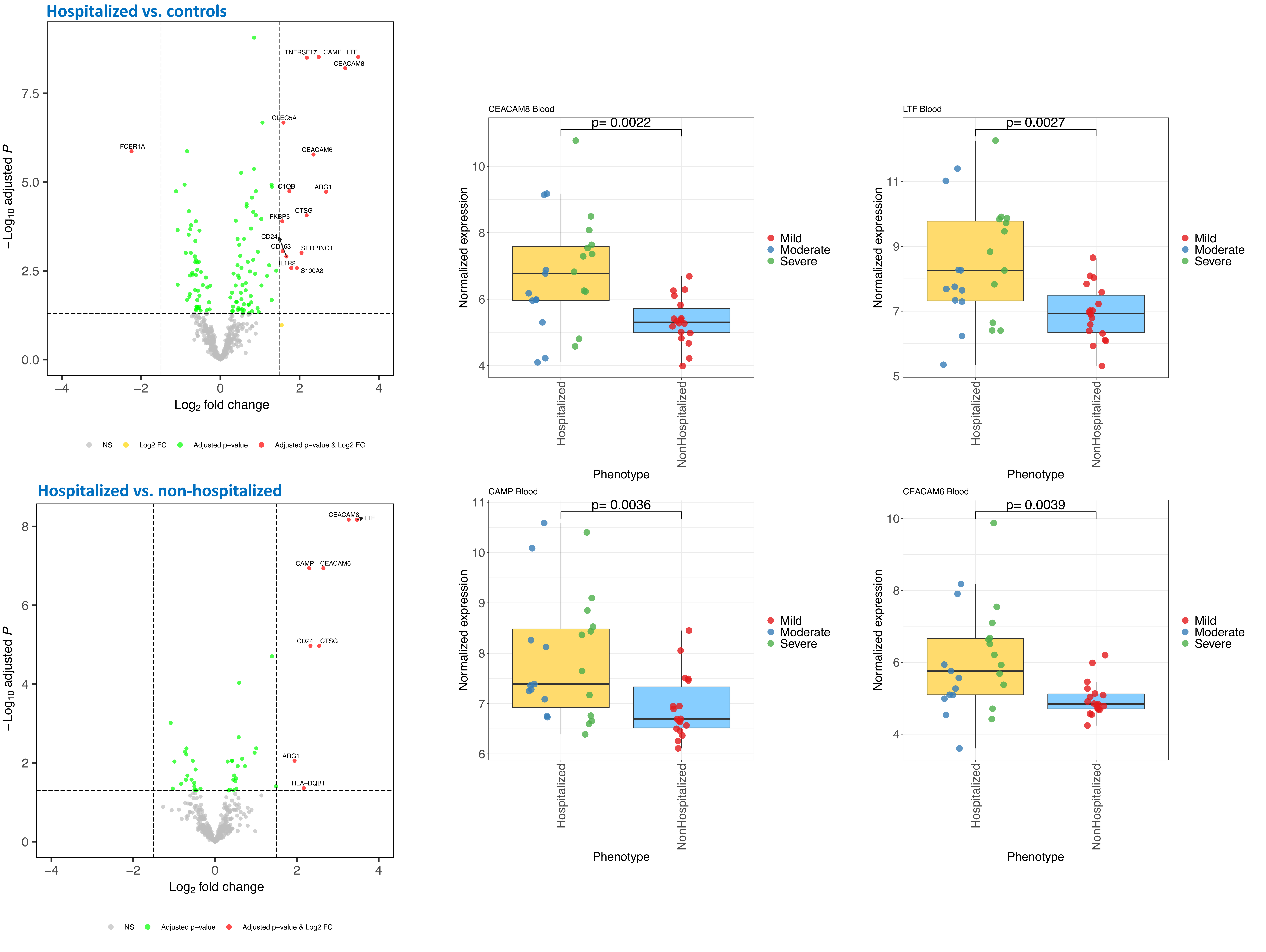

Hospitalized vs. controls
TNFRSF17
CAMP
LTF
CEACAM8
CLEC5A
FCER1A
CEACAM6
C1QB
ARG1
CTSG
FKBP5
CD24
CD163
SERPING1
IL1R2
S100A8
−Log10 adjusted P
Log2 fold change
NS
Log2 FC
Adjusted p−value
Adjusted p−value & Log2 FC
CEACAM8 Blood
p= 0.0022
Normalized expression
Hospitalized
NonHospitalized
Phenotype
Mild
Moderate
Severe
LTF Blood
p= 0.0027
Normalized expression
Hospitalized
NonHospitalized
Phenotype
Mild
Moderate
Severe
Hospitalized vs. non-hospitalized
CEACAM8
LTF
CAMP
CEACAM6
CD24
CTSG
ARG1
HLA−DQB1
−Log10 adjusted P
Log2 fold change
NS
Adjusted p−value
Adjusted p−value & Log2 FC
CAMP Blood
p= 0.0036
Normalized expression
Hospitalized
NonHospitalized
Phenotype
Mild
Moderate
Severe
CEACAM6 Blood
p= 0.0039
Normalized expression
Hospitalized
NonHospitalized
Phenotype
Mild
Moderate
Severe

### Figure S7. Severe vs Control

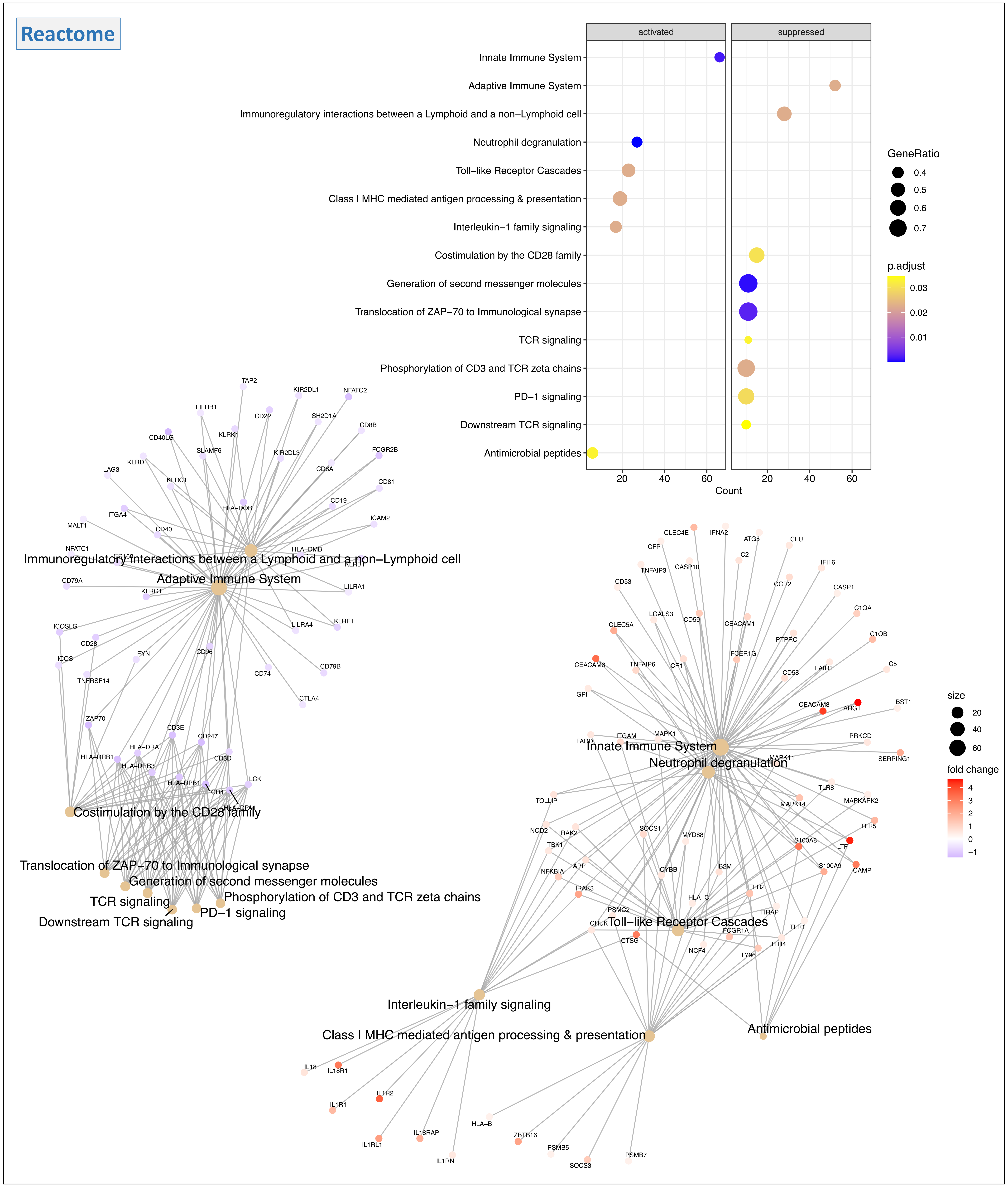


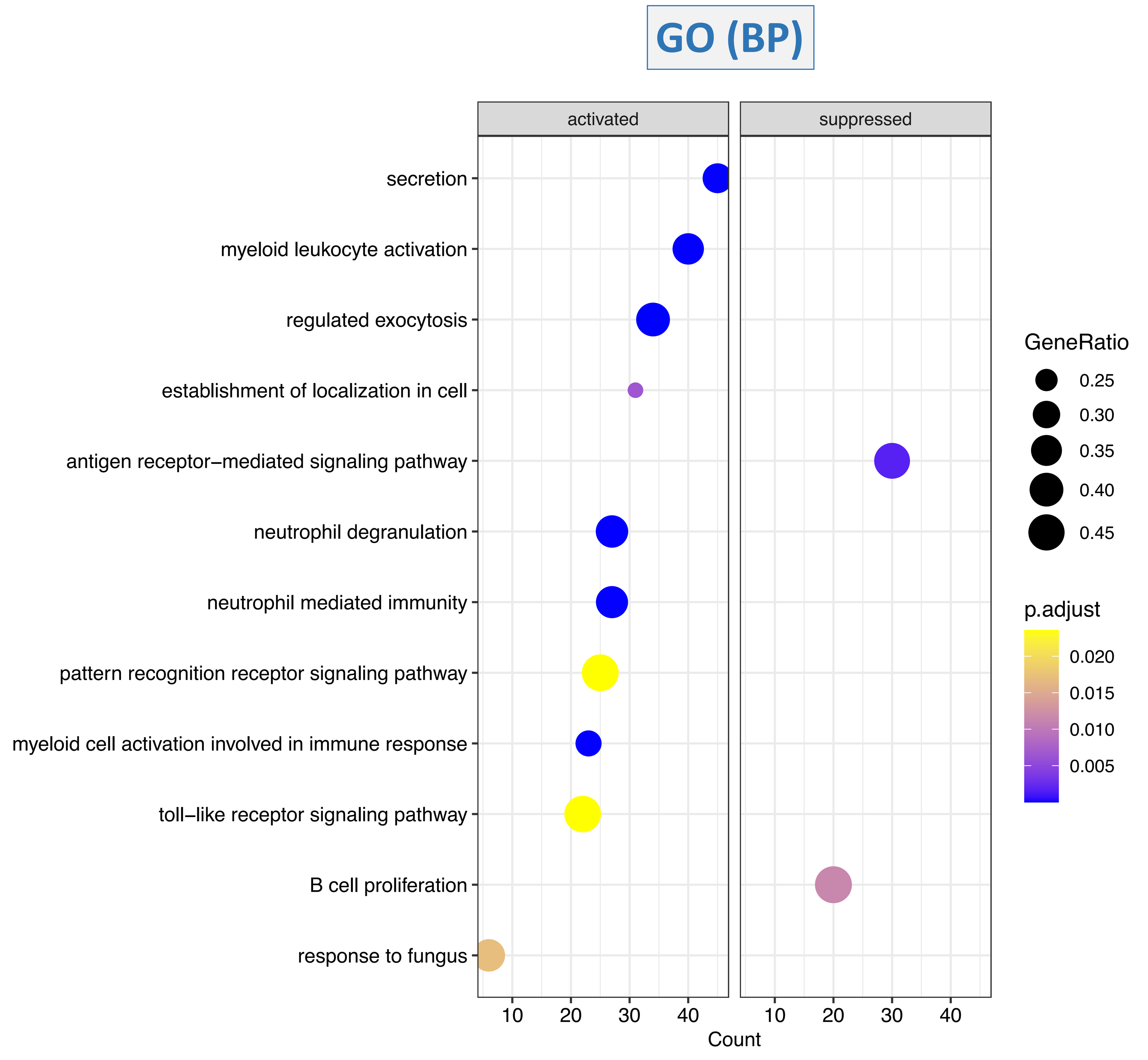

### Figure S8. Severe vs Moderate

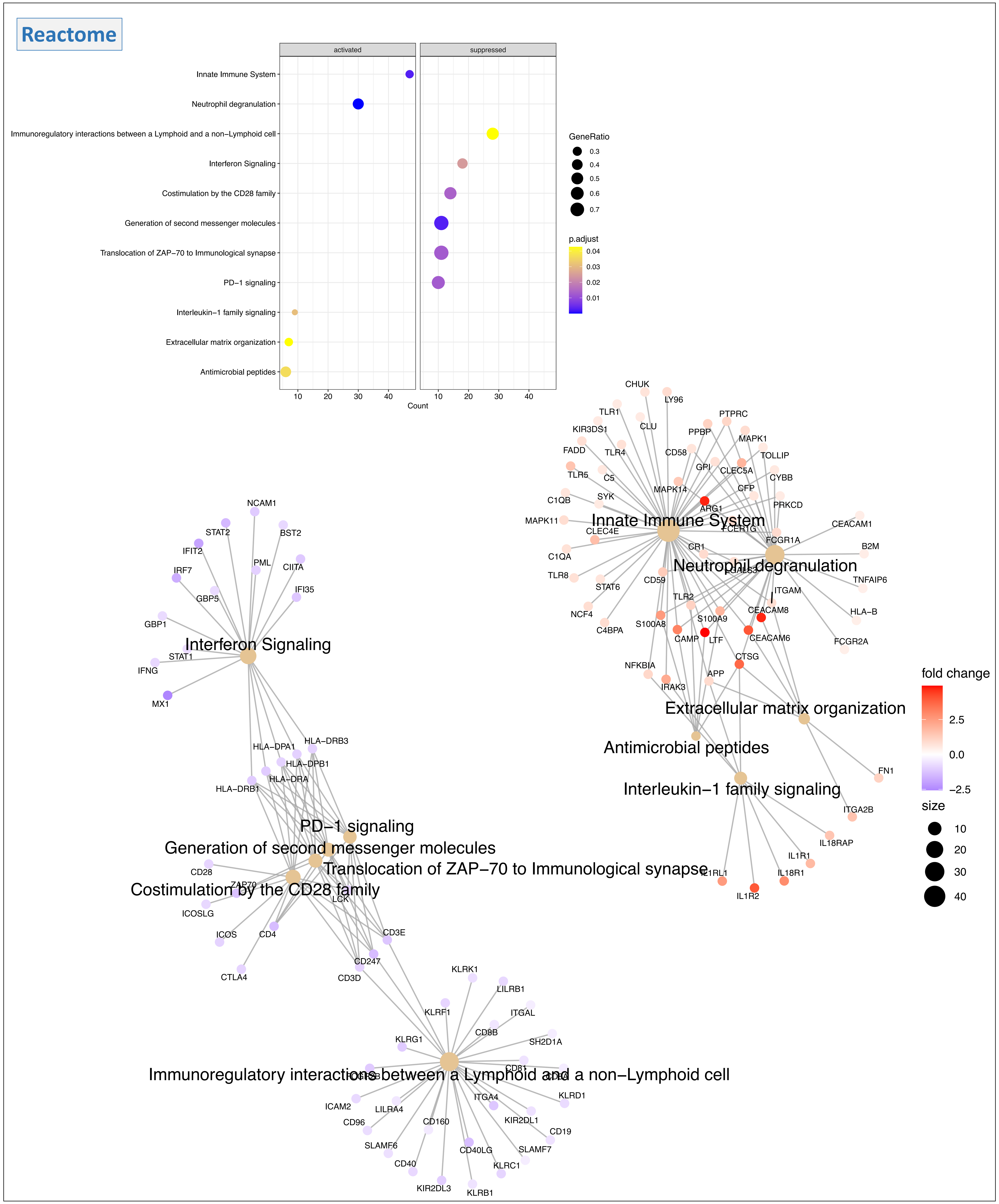


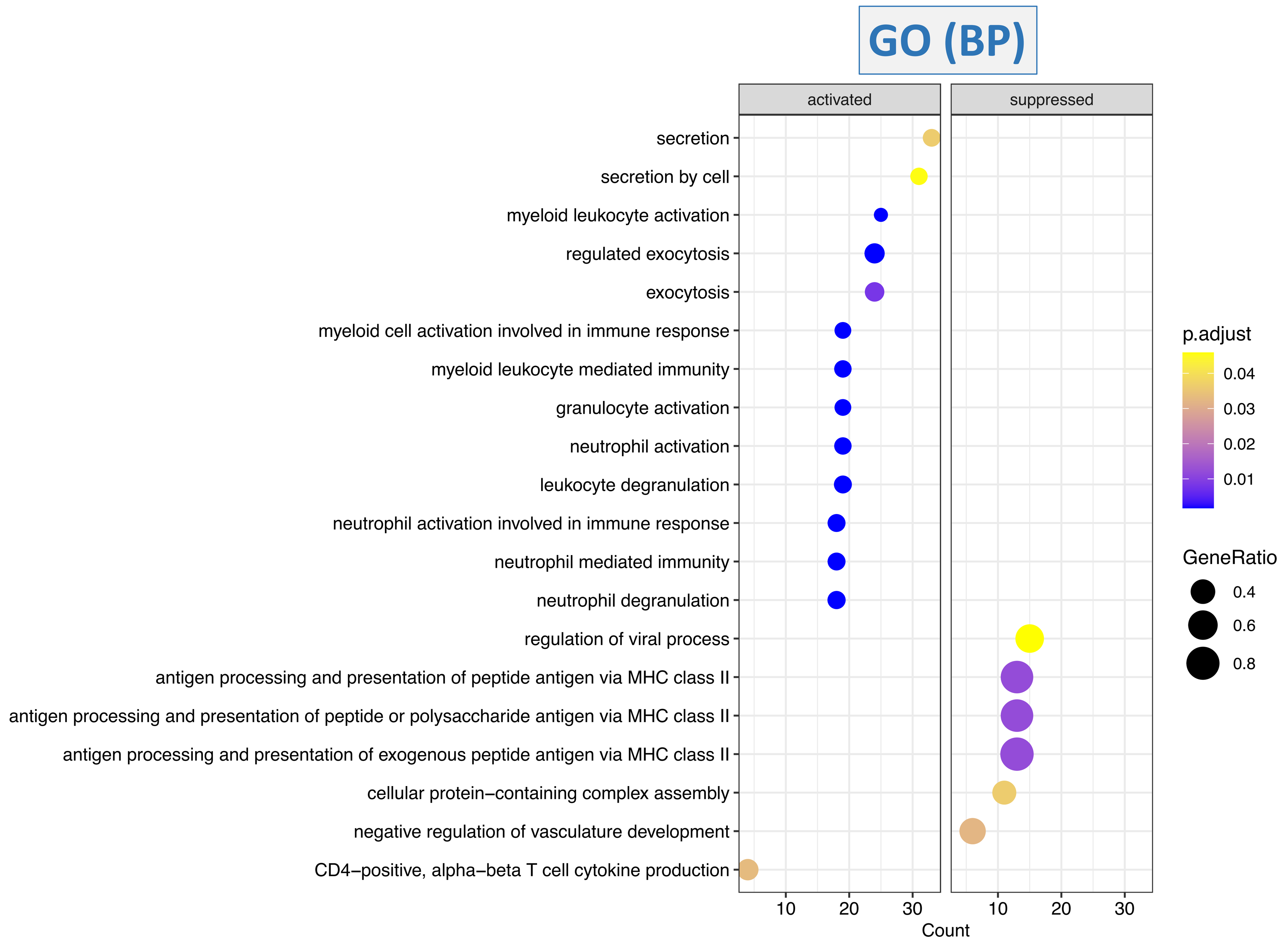

### Figure S9. Severe vs Mild

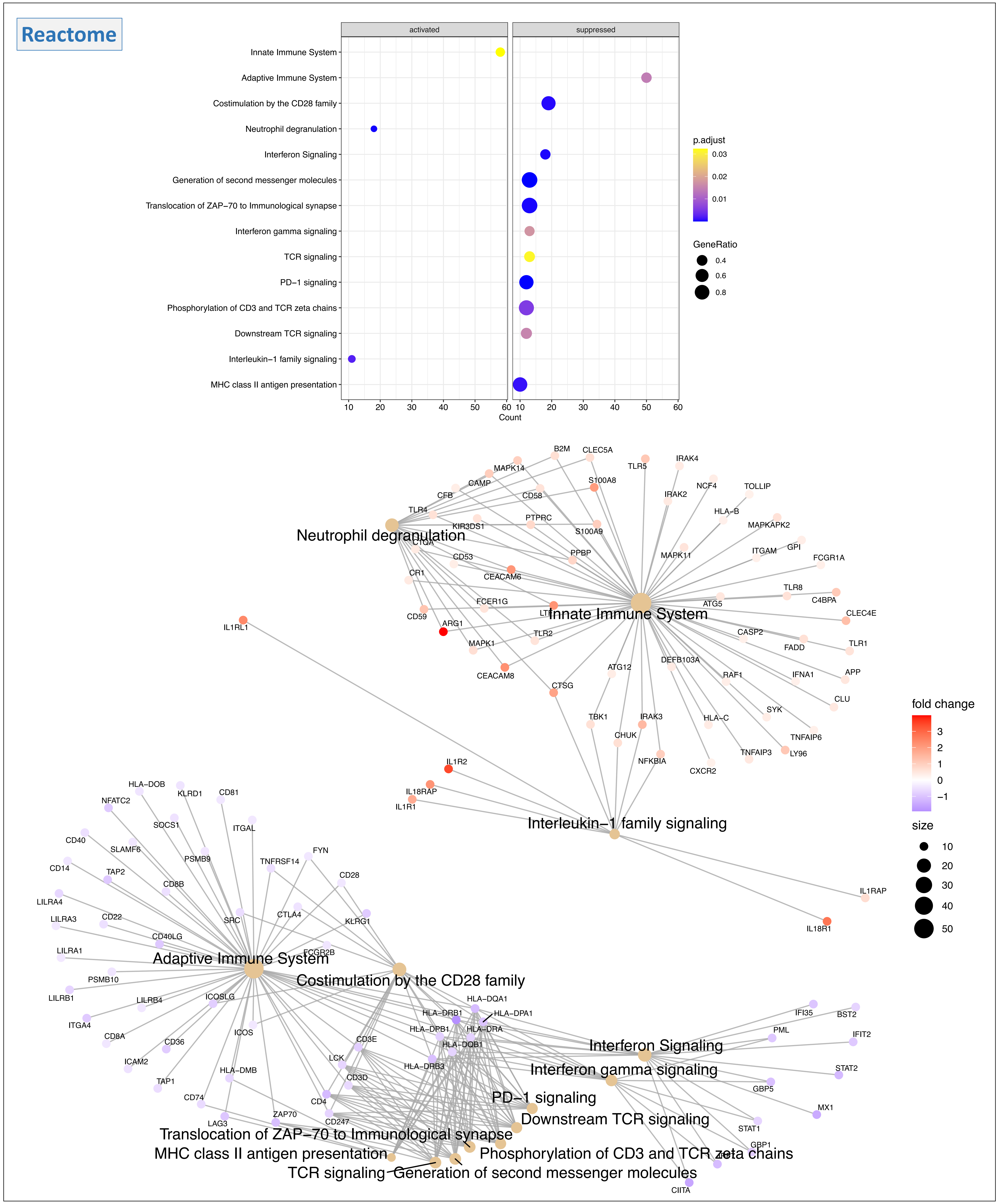


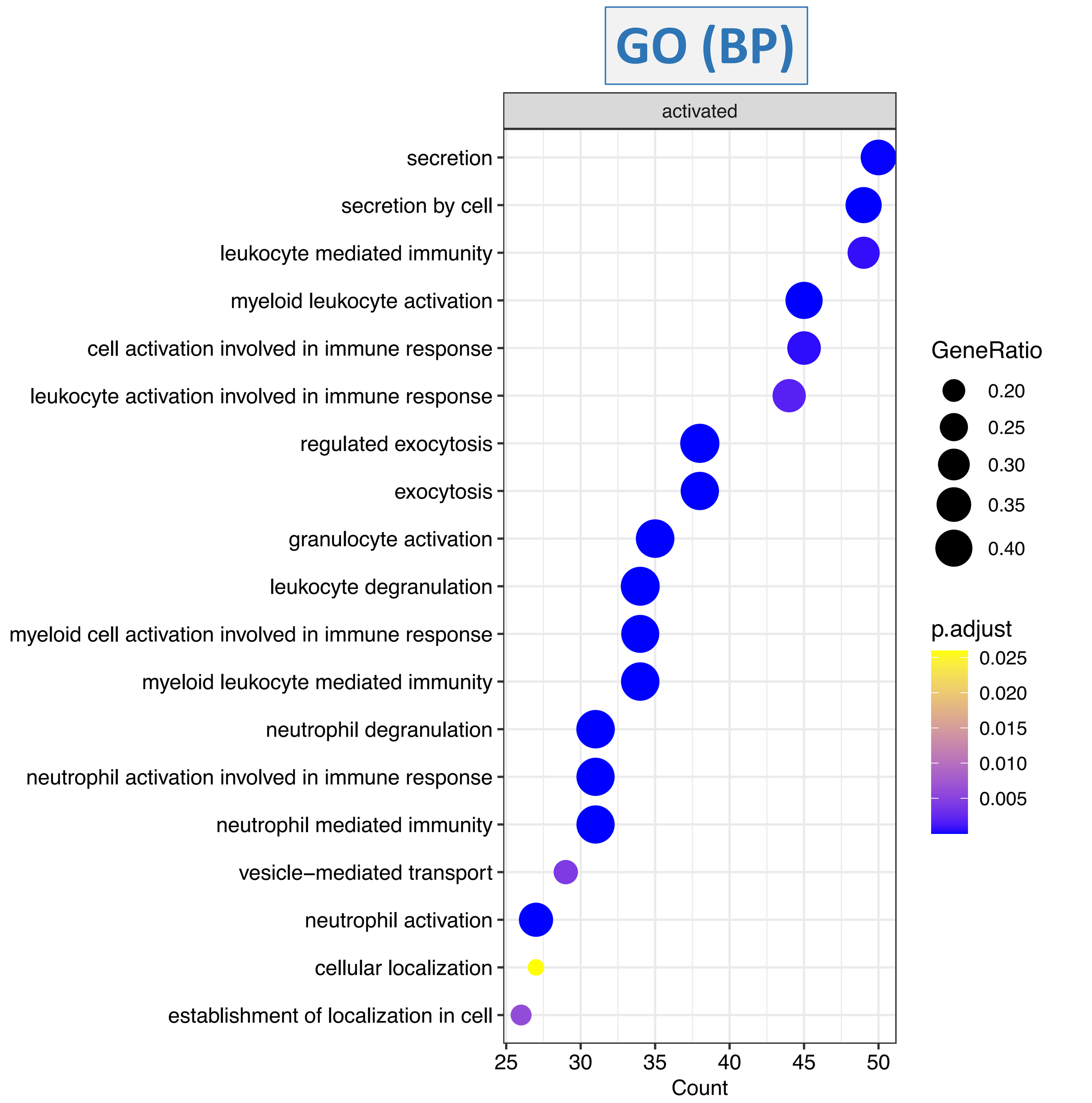

**Figure S10.**

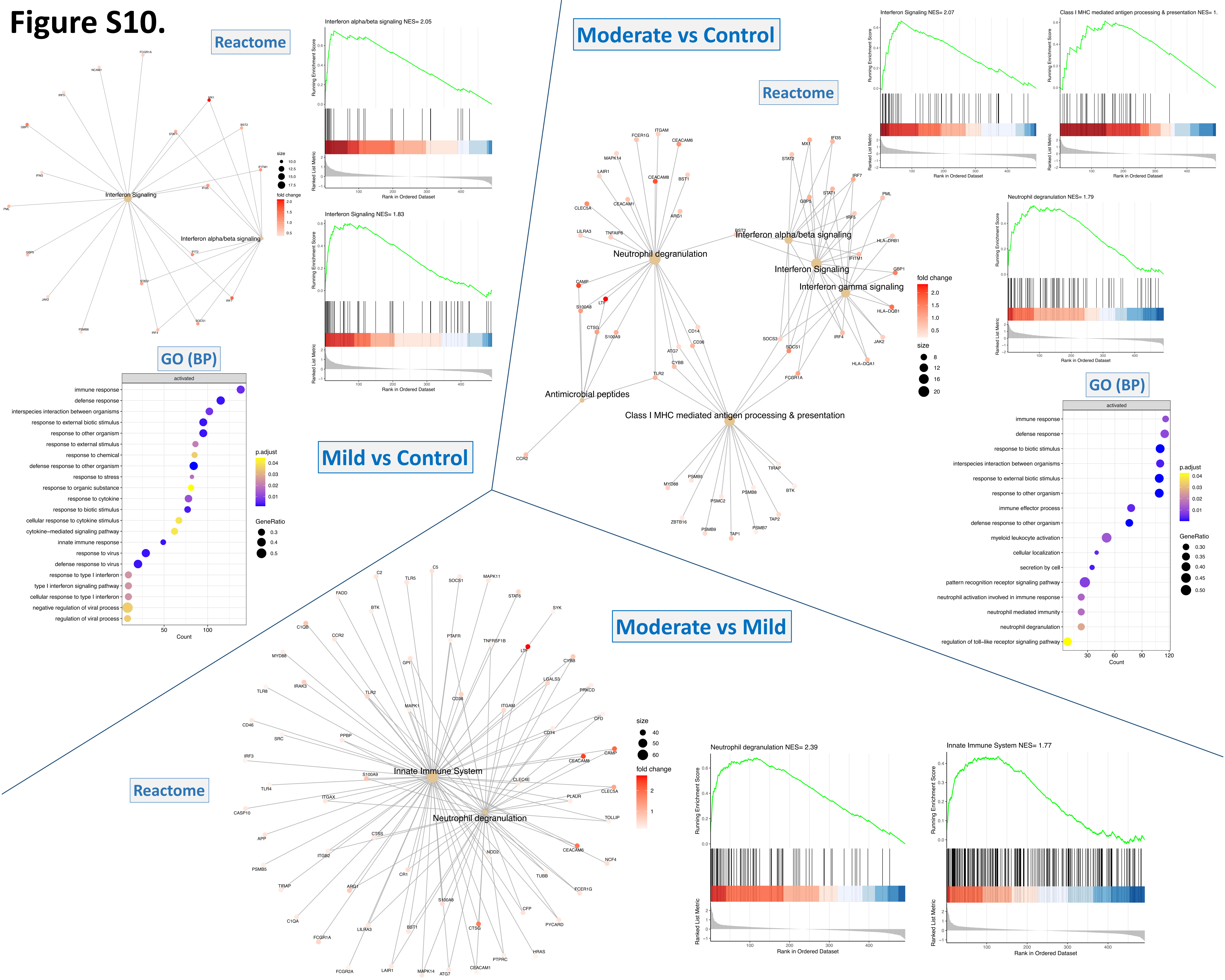

Mild vs Control
Reactome
Interferon Signaling
Interferon alpha/beta signaling
size
fold change
Interferon alpha/beta signaling NES= 2.05
Interferon Signaling NES= 1.83
Running Enrichment Score
Ranked List Metric
Rank in Ordered Dataset
GO (BP)
activated
immune response
defense response
interspecies interaction between organisms
response to external biotic stimulus
response to other organism
response to external stimulus
response to chemical
defense response to other organism
response to stress
response to organic substance
response to cytokine
response to biotic stimulus
cellular response to cytokine stimulus
cytokine−mediated signaling pathway
innate immune response
response to virus
defense response to virus
response to type I interferon
type I interferon signaling pathway
cellular response to type I interferon
negative regulation of viral process
regulation of viral process
Count
p.adjust
GeneRatio
Moderate vs Control
Reactome
Neutrophil degranulation
Antimicrobial peptides
Class I MHC mediated antigen processing & presentation
Interferon gamma signaling
Interferon Signaling NES= 2.07
Class I MHC mediated antigen processing & presentation NES= 1.
Neutrophil degranulation NES= 1.79
GO (BP)
activated
immune response
defense response
response to biotic stimulus
interspecies interaction between organisms
response to external biotic stimulus
response to other organism
immune effector process
defense response to other organism
myeloid leukocyte activation
cellular localization
secretion by cell
pattern recognition receptor signaling pathway
neutrophil activation involved in immune response
neutrophil mediated immunity
neutrophil degranulation
regulation of toll−like receptor signaling pathway
Count
p.adjust
GeneRatio
Moderate vs Mild
Reactome
Innate Immune System
Neutrophil degranulation
size
fold change
Neutrophil degranulation NES= 2.39
Innate Immune System NES= 1.77
Running Enrichment Score
Ranked List Metric
Rank in Ordered Dataset

### FIGURE S11.

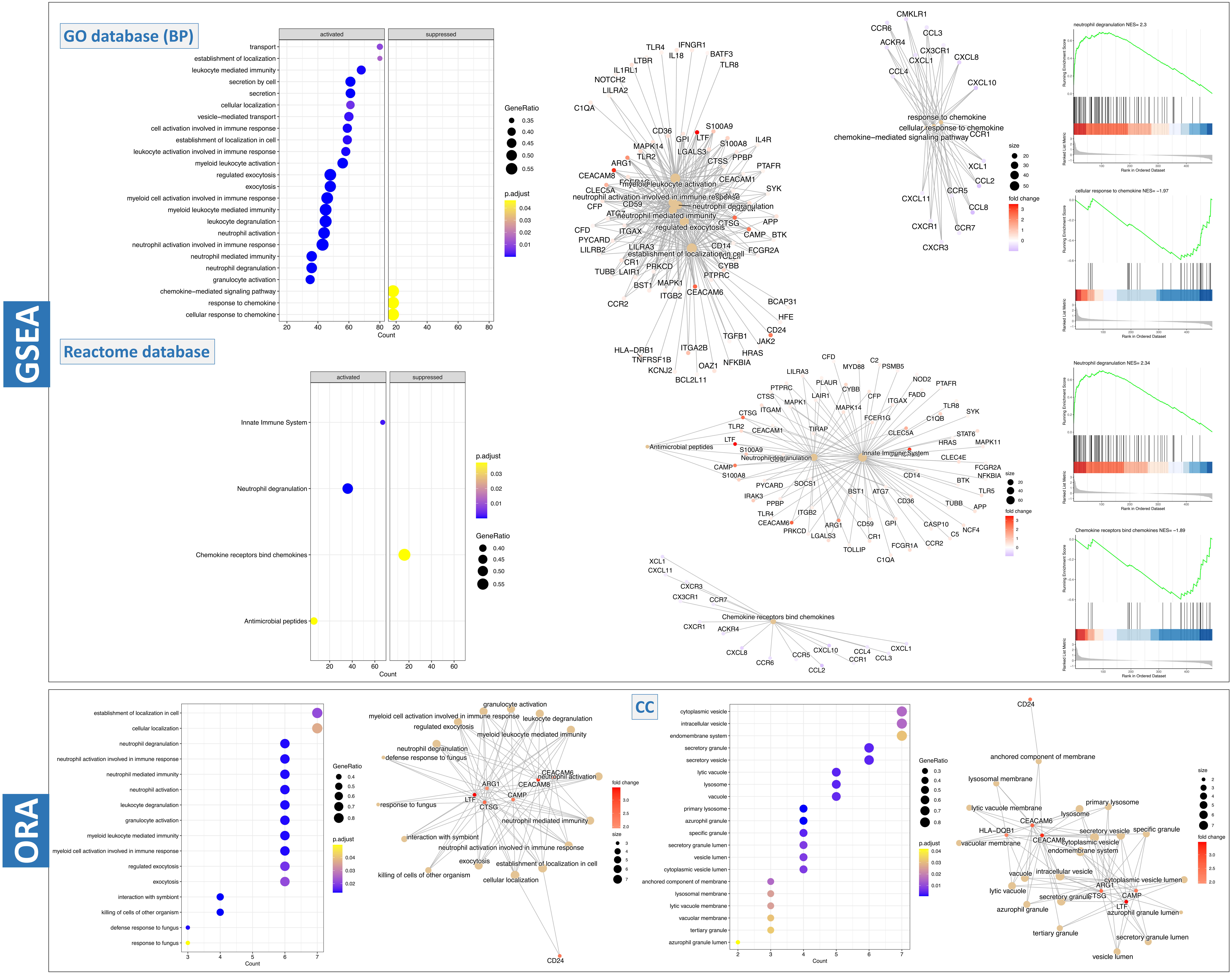
GSEA
GO database (BP)
activated
suppressed
transport
establishment of localization
leukocyte mediated immunity
secretion by cell
secretion
cellular localization
vesicle−mediated transport
cell activation involved in immune response
establishment of localization in cell
leukocyte activation involved in immune response
myeloid leukocyte activation
regulated exocytosis
exocytosis
myeloid cell activation involved in immune response
myeloid leukocyte mediated immunity
leukocyte degranulation
neutrophil activation
neutrophil activation involved in immune response
neutrophil mediated immunity
neutrophil degranulation
granulocyte activation
chemokine−mediated signaling pathway
response to chemokine
cellular response to chemokine
Count
GeneRatio
p.adjust
size
fold change
neutrophil degranulation NES= 2.3
cellular response to chemokine NES= −1.97
Running Enrichment Score
Ranked List Metric
Rank in Ordered Dataset
Reactome database
Innate Immune System
Neutrophil degranulation
Chemokine receptors bind chemokines
Antimicrobial peptides
Neutrophil degranulation NES= 2.34
Chemokine receptors bind chemokines NES= −1.89
ORA
establishment of localization in cell
cellular localization
neutrophil degranulation
neutrophil activation involved in immune response
neutrophil mediated immunity
neutrophil activation
leukocyte degranulation
granulocyte activation
myeloid leukocyte mediated immunity
myeloid cell activation involved in immune response
regulated exocytosis
exocytosis
interaction with symbiont
killing of cells of other organism
defense response to fungus
response to fungus
CC
cytoplasmic vesicle
intracellular vesicle
endomembrane system
secretory granule
secretory vesicle
lytic vacuole
lysosome
vacuole
primary lysosome
azurophil granule
specific granule
secretory granule lumen
vesicle lumen
cytoplasmic vesicle lumen
anchored component of membrane
lysosomal membrane
lytic vacuole membrane
vacuolar membrane
tertiary granule
azurophil granule lumen

### Figure S12.

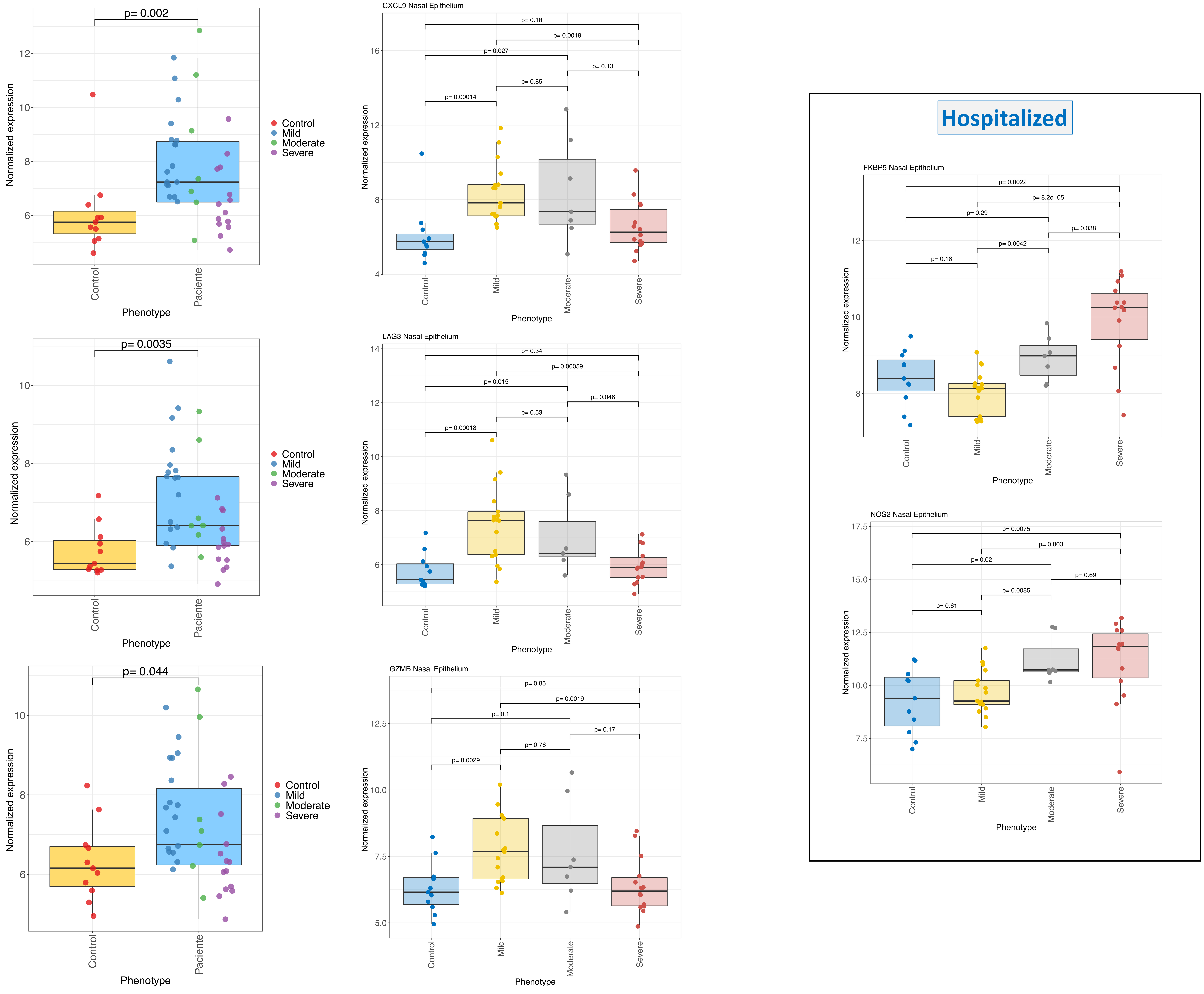

p= 0.002
Normalized expression
Control
Paciente
Phenotype
Control
Mild
Moderate
Severe
p= 0.0035
p= 0.044
CXCL9 Nasal Epithelium
p= 0.18
p= 0.0019
p= 0.027
p= 0.13
p= 0.85
p= 0.00014
LAG3 Nasal Epithelium
p= 0.34
p= 0.00059
p= 0.015
p= 0.046
p= 0.53
p= 0.00018
GZMB Nasal Epithelium
p= 0.85
p= 0.0019
p= 0.1
p= 0.17
p= 0.76
p= 0.0029
Hospitalized
FKBP5 Nasal Epithelium
p= 0.0022
p= 8.2e−05
p= 0.29
p= 0.038
p= 0.0042
p= 0.16
NOS2 Nasal Epithelium
p= 0.0075
p= 0.003
p= 0.02
p= 0.69
p= 0.0085
p= 0.61

### Figure S13.

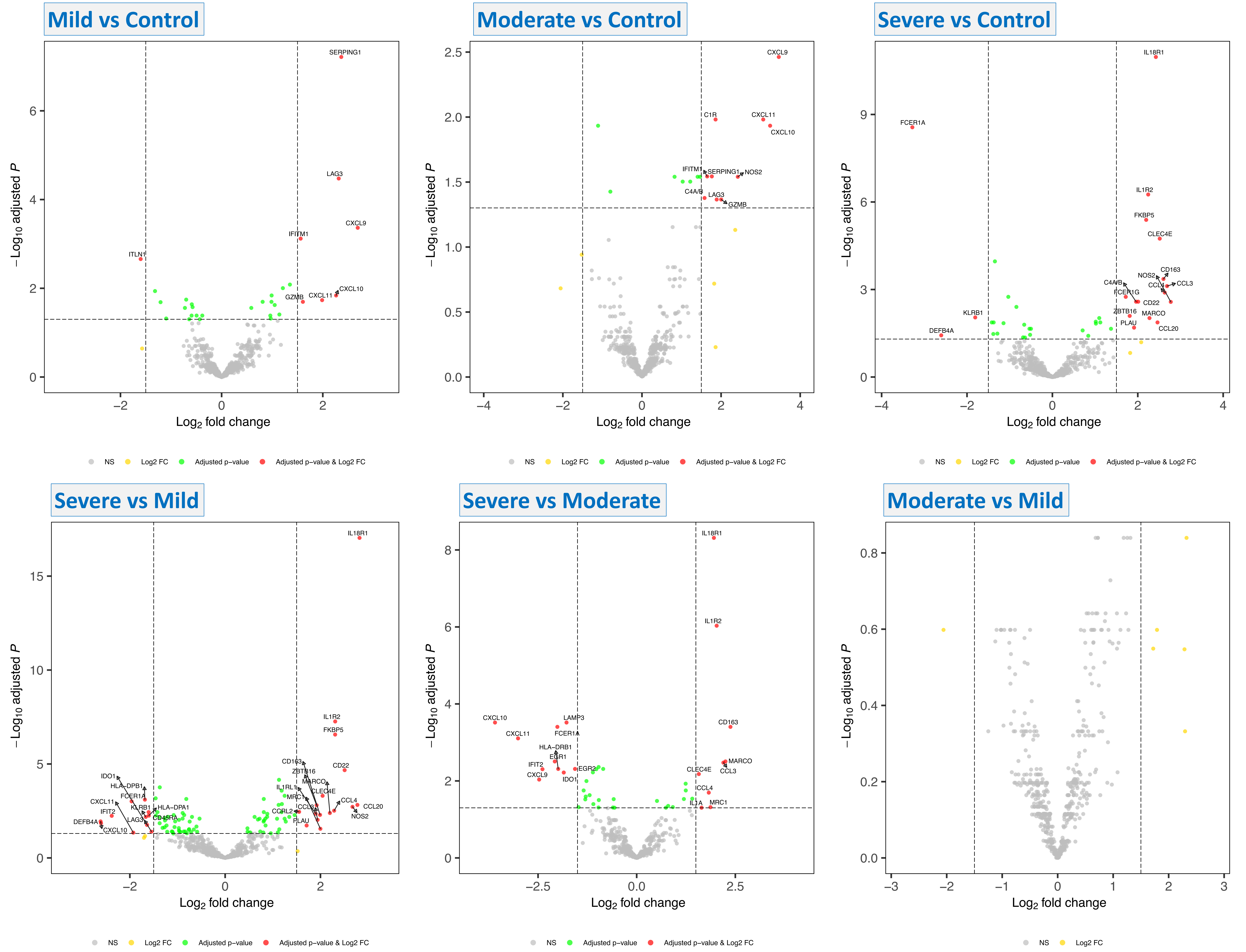
Mild vs Control
SERPING1
LAG3
CXCL9
IFITM1
ITLN1
CXCL10
CXCL11
GZMB
−Log10 adjusted P
Log2 fold change
NS
Log2 FC
Adjusted p−value
Adjusted p−value & Log2 FC
Moderate vs Control
CXCL9
C1R
CXCL11
CXCL10
IFITM1
SERPING1
NOS2
C4A/B
LAG3
GZMB
−Log10 adjusted P
Log2 fold change
NS
Log2 FC
Adjusted p−value
Adjusted p−value & Log2 FC
Severe vs Control
IL18R1
FCER1A
IL1R2
FKBP5
CLEC4E
CD163
NOS2
CCL3
CCL4
C4A/B
FCER1G
CD22
KLRB1
ZBTB16
MARCO
PLAU
DEFB4A
CCL20
−Log10 adjusted P
Log2 fold change
NS
Log2 FC
Adjusted p−value
Adjusted p−value & Log2 FC
Severe vs Mild
IL18R1
IL1R2
FKBP5
CD163
CD22
ZBTB16
IDO1
HLA−DPB1
MARCO
IL1RL1
CLEC4E
FCER1A
MRC1
CXCL11
CCL4
KLRB1
HLA−DPA1
CCL3
CCL20
IFIT2
CCRL2
NOS2
CD45RA
LAG3
PLAU
DEFB4A
CXCL10
−Log10 adjusted P
Log2 fold change
NS
Log2 FC
Adjusted p−value
Adjusted p−value & Log2 FC
Severe vs Moderate
IL18R1
IL1R2
CXCL10
LAMP3
CD163
CXCL11
FCER1A
HLA−DRB1
EGR1
MARCO
IFIT2
CCL3
EGR2
CLEC4E
CXCL9
IDO1
CCL4
IL1A
MRC1
−Log10 adjusted P
Log2 fold change
NS
Adjusted p−value
Adjusted p−value & Log2 FC
Moderate vs Mild
−Log10 adjusted P
Log2 fold change
NS
Log2 FC

### Figure S14.

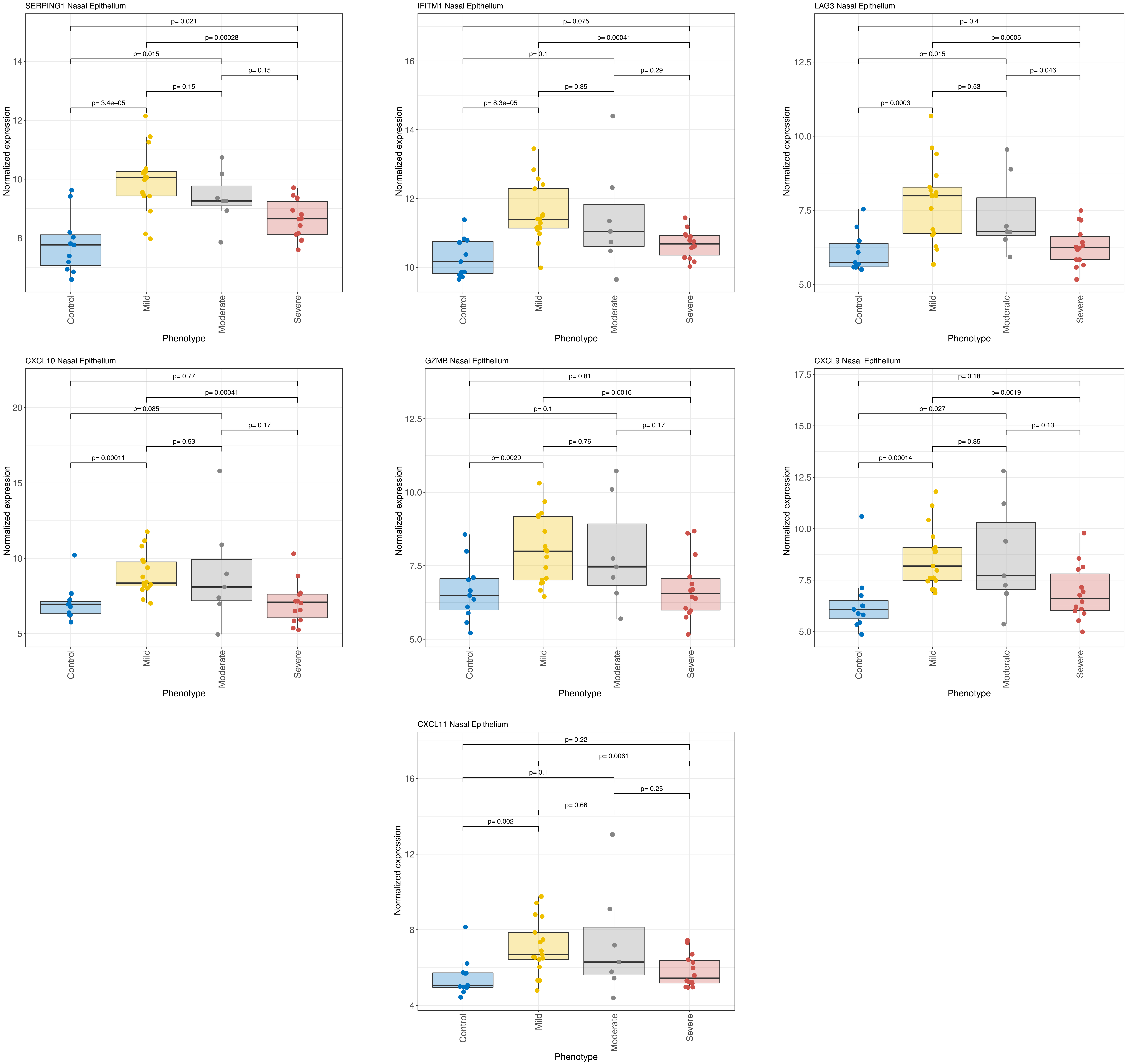
SERPING1 Nasal Epithelium
p= 0.021
p= 0.00028
p= 0.015
p= 0.15
p= 0.15
p= 3.4e−05
IFITM1 Nasal Epithelium
p= 0.075
p= 0.00041
p= 0.1
p= 0.29
p= 0.35
p= 8.3e−05
LAG3 Nasal Epithelium
p= 0.4
p= 0.0005
p= 0.015
p= 0.046
p= 0.53
p= 0.0003
CXCL10 Nasal Epithelium
p= 0.77
p= 0.00041
p= 0.085
p= 0.17
p= 0.53
p= 0.00011
GZMB Nasal Epithelium
p= 0.81
p= 0.0016
p= 0.1
p= 0.17
p= 0.76
p= 0.0029
CXCL9 Nasal Epithelium
p= 0.18
p= 0.0019
p= 0.027
p= 0.13
p= 0.85
p= 0.00014
CXCL11 Nasal Epithelium
p= 0.22
p= 0.0061
p= 0.1
p= 0.25
p= 0.66
p= 0.002
Normalized expression
Phenotype
Control
Mild
Moderate
Severe

### Figure S15.

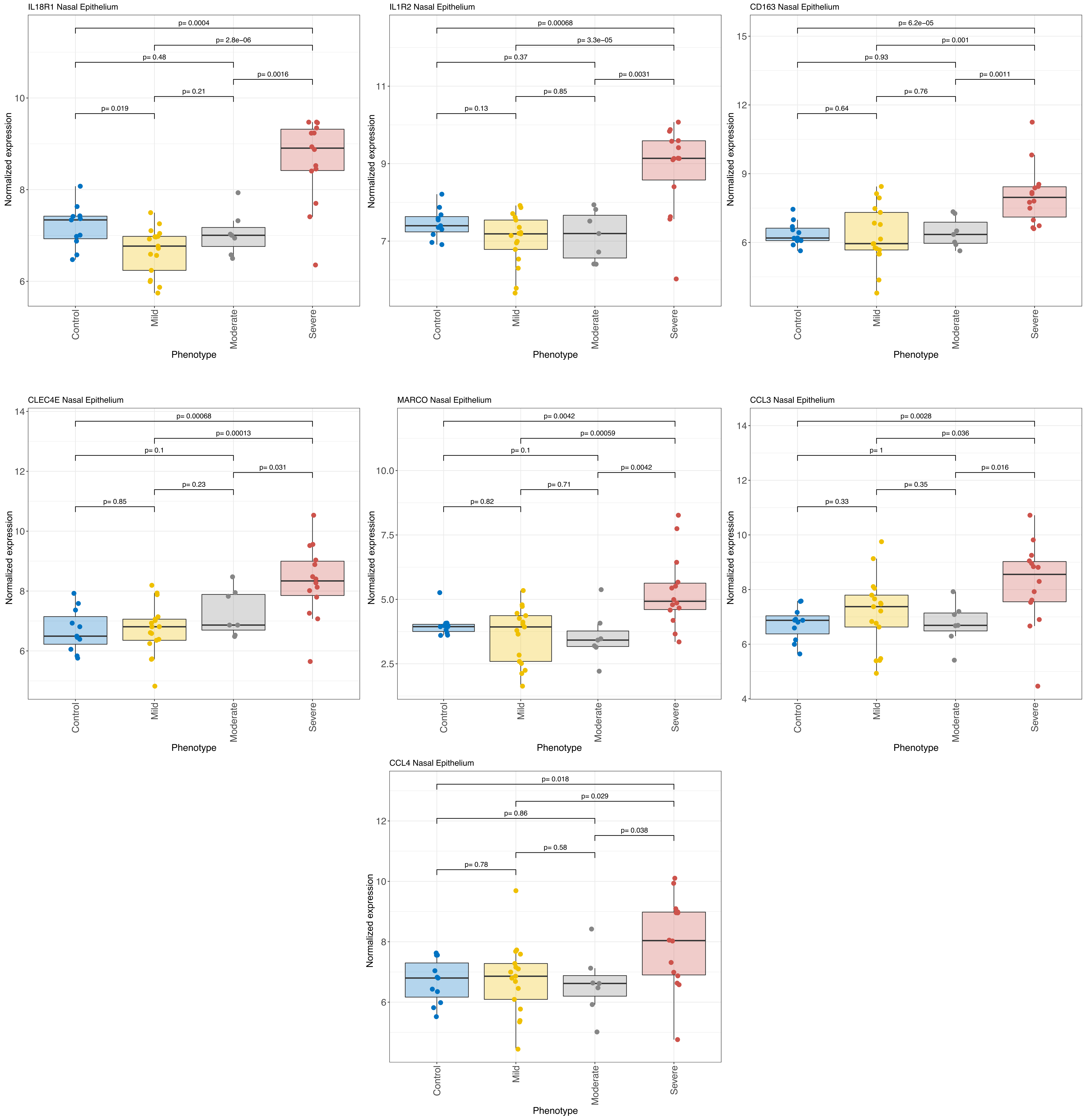

IL18R1 Nasal Epithelium
p= 0.0004
p= 2.8e−06
p= 0.48
p= 0.0016
p= 0.21
p= 0.019
IL1R2 Nasal Epithelium
p= 0.00068
p= 3.3e−05
p= 0.37
p= 0.0031
p= 0.85
p= 0.13
CD163 Nasal Epithelium
p= 6.2e−05
p= 0.001
p= 0.93
p= 0.0011
p= 0.76
p= 0.64
CLEC4E Nasal Epithelium
p= 0.00068
p= 0.00013
p= 0.1
p= 0.031
p= 0.23
p= 0.85
MARCO Nasal Epithelium
p= 0.0042
p= 0.00059
p= 0.1
p= 0.0042
p= 0.71
p= 0.82
CCL3 Nasal Epithelium
p= 0.0028
p= 0.036
p= 1
p= 0.016
p= 0.35
p= 0.33
CCL4 Nasal Epithelium
p= 0.018
p= 0.029
p= 0.86
p= 0.038
p= 0.58
p= 0.78
Normalized expression
Control
Mild
Moderate
Severe
Phenotype

### Figure S16.

#### Moderate vs Control

**GSEA** **GO (BP)**

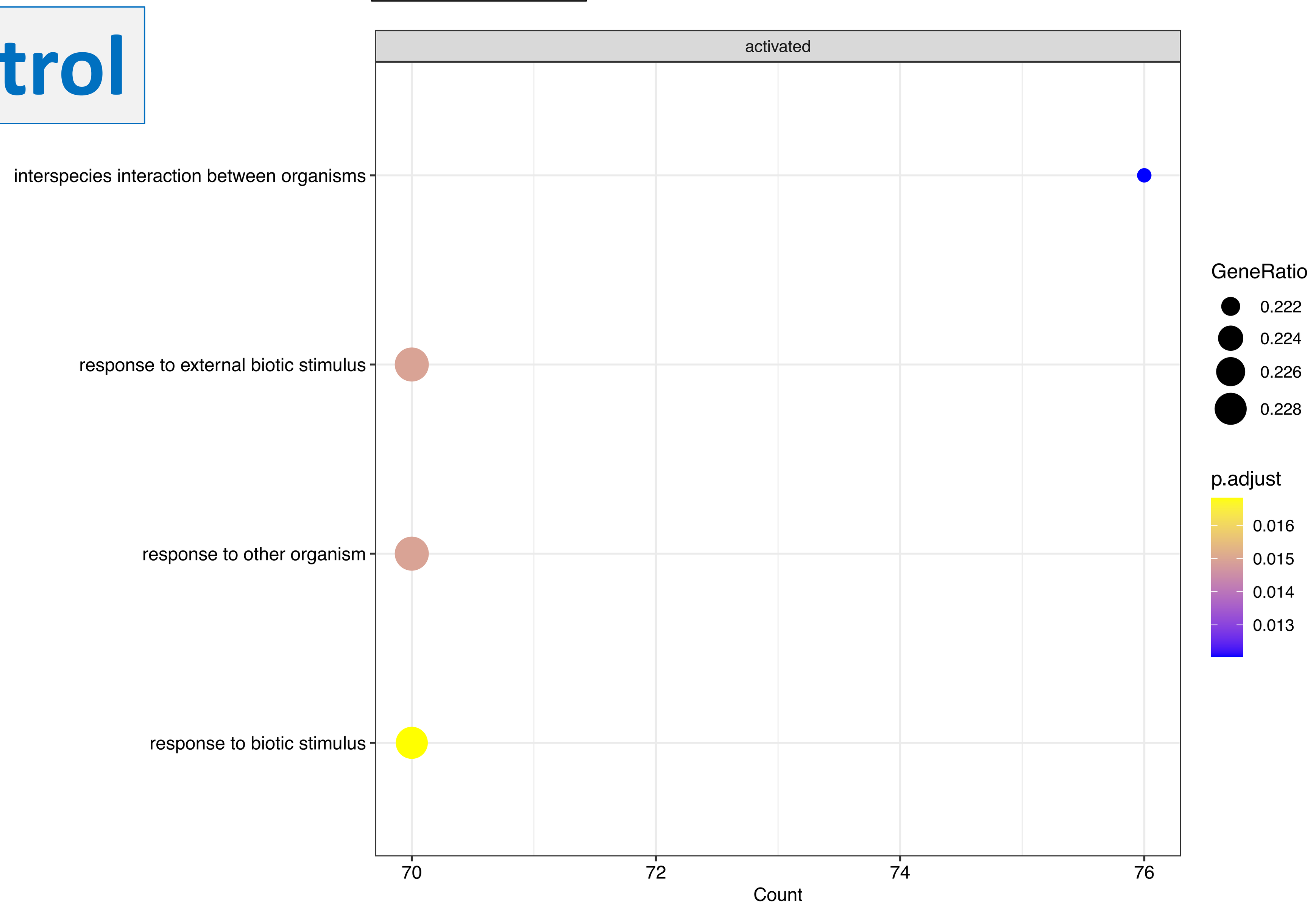


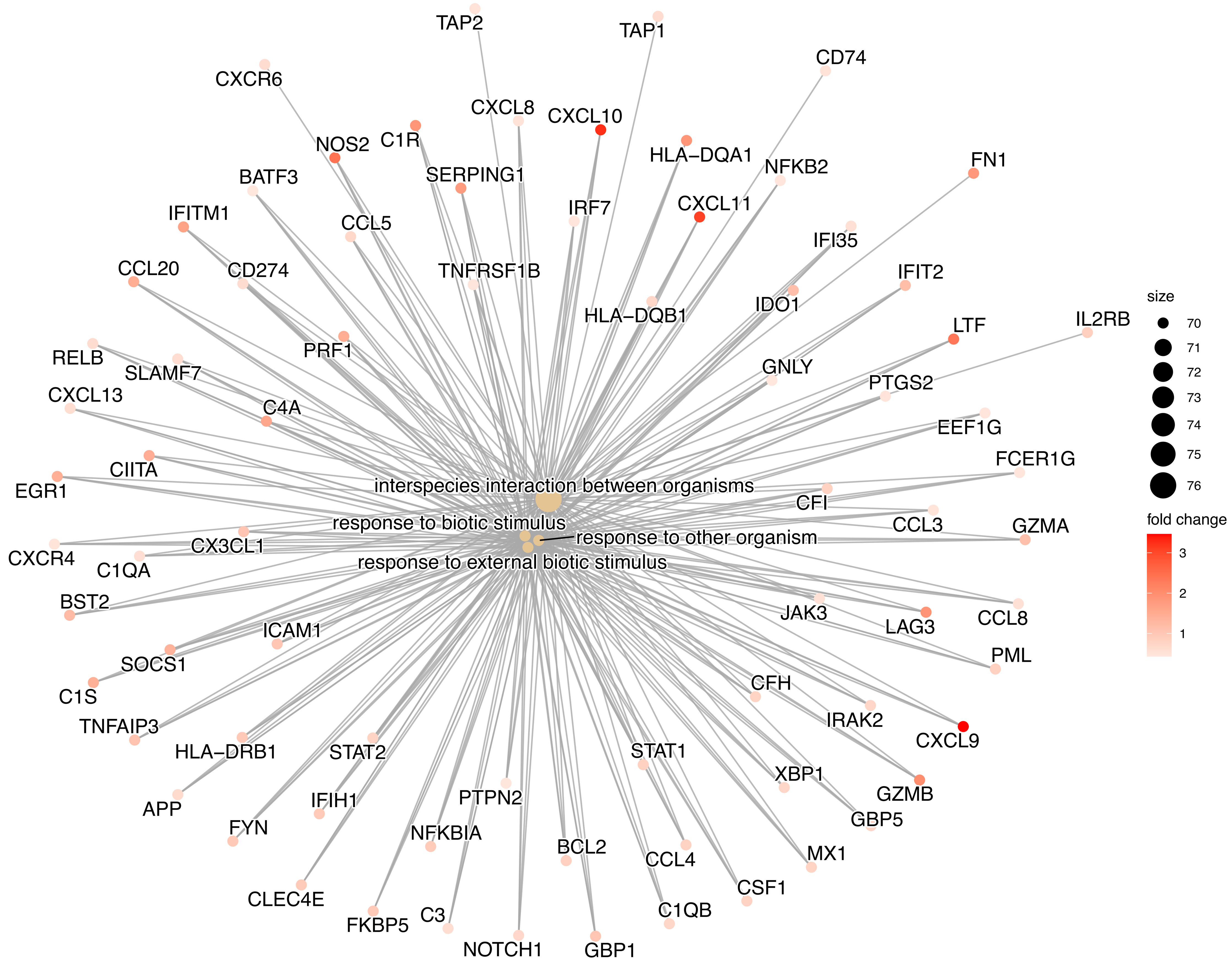


**ORA** **GO (MF)**

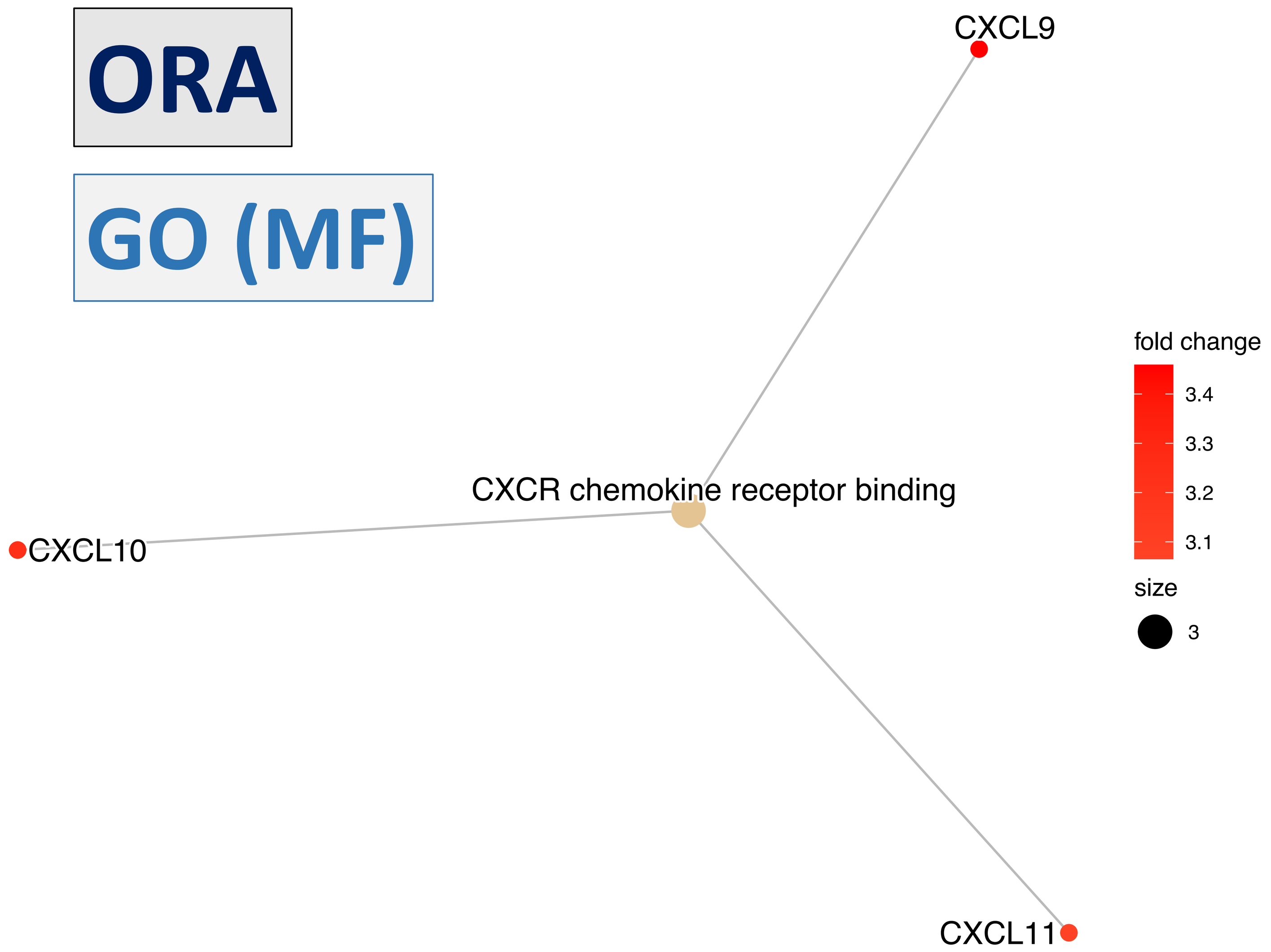


**GSEA** **GO (MF)**

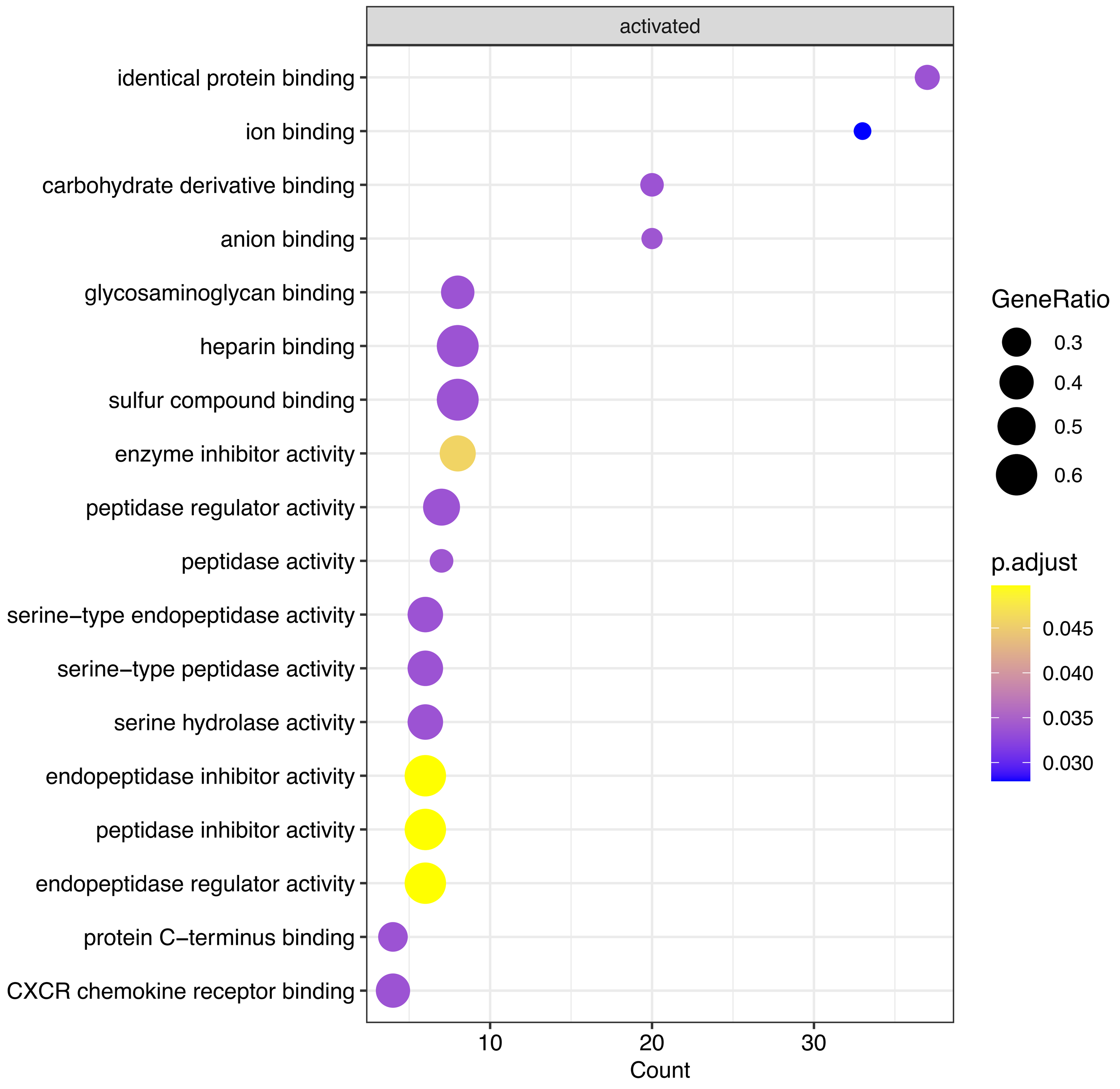


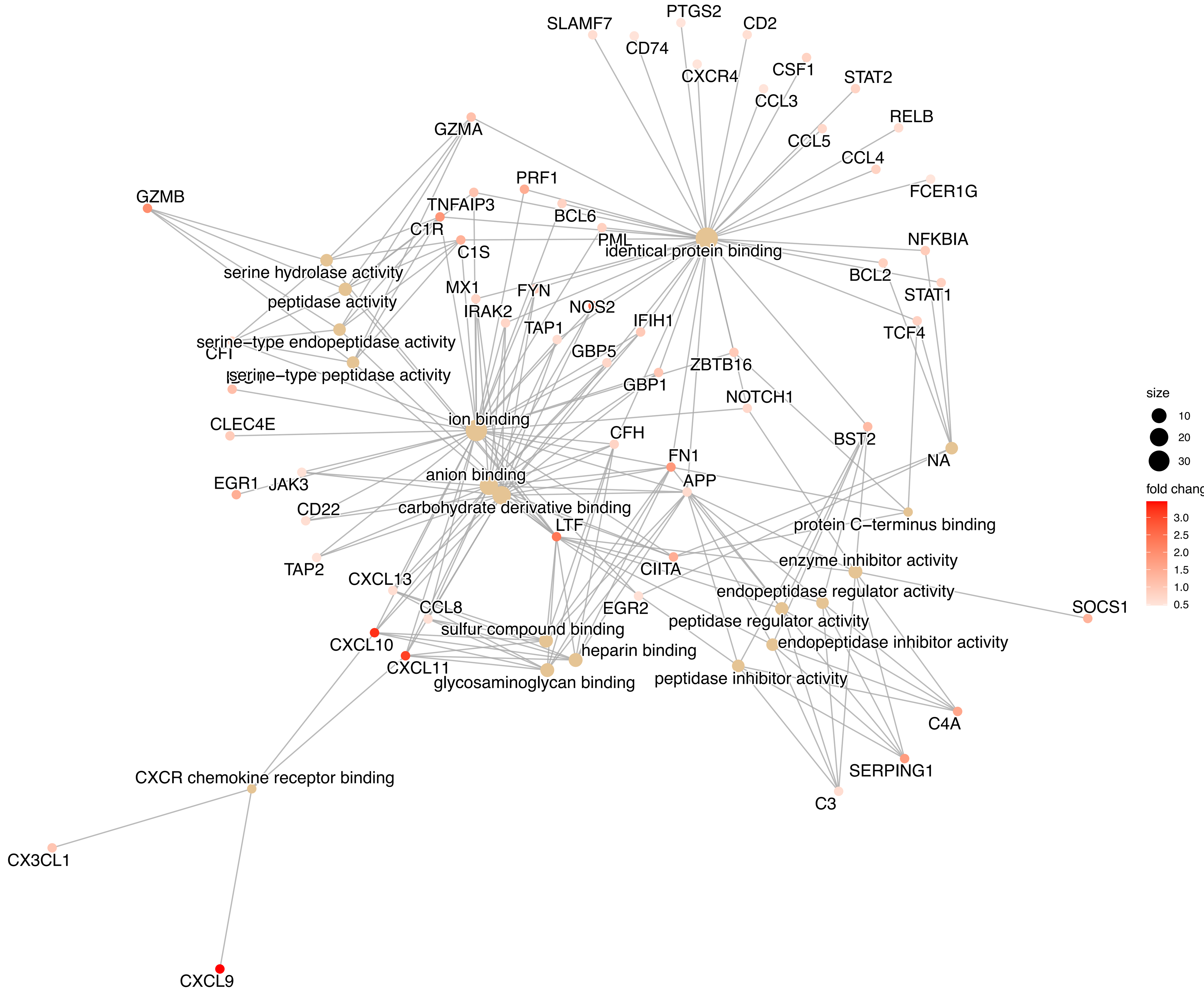

Figure S17.

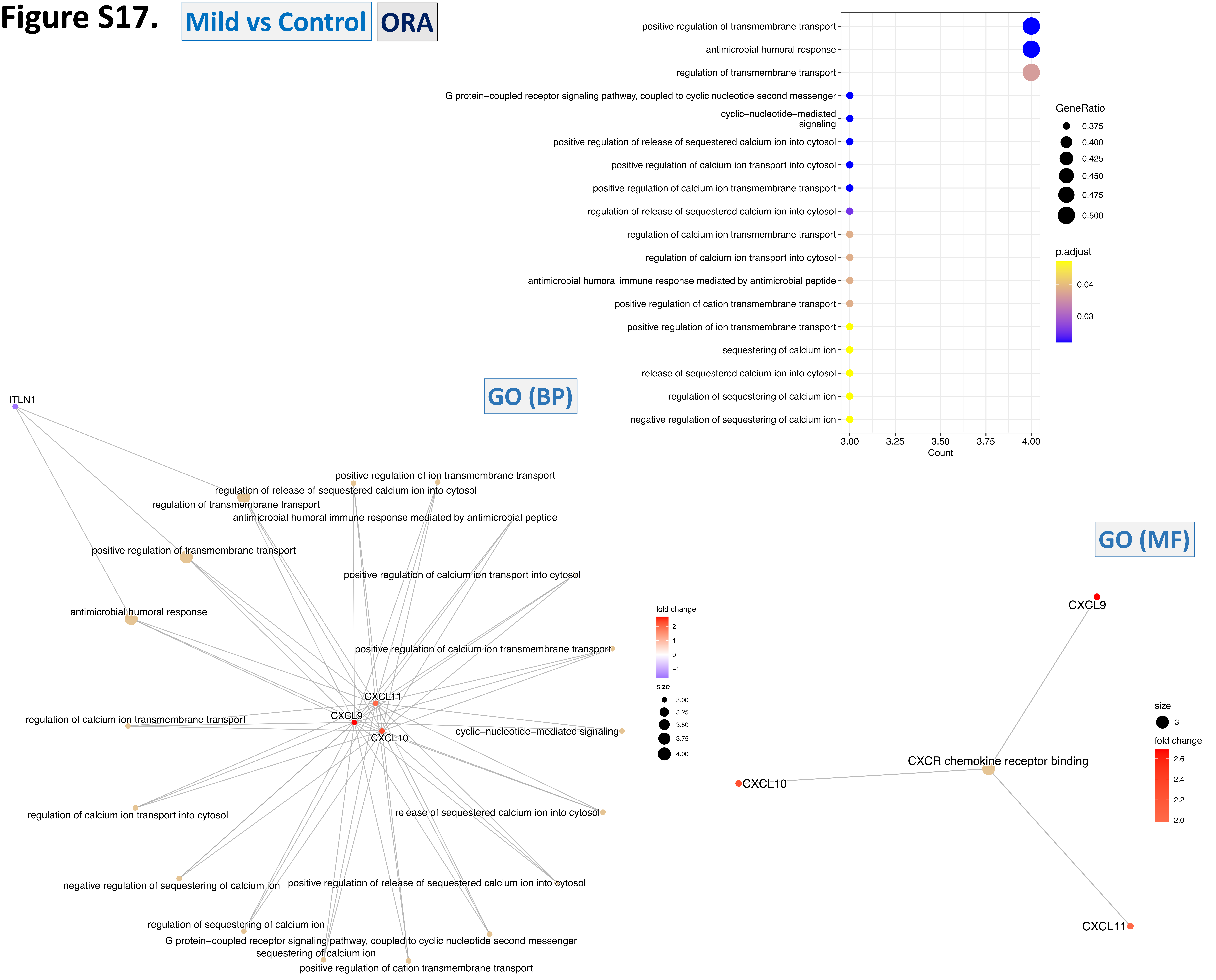
Mild vs Control
ORA
positive regulation of transmembrane transport
antimicrobial humoral response
regulation of transmembrane transport
G protein−coupled receptor signaling pathway, coupled to cyclic nucleotide second messenger
cyclic−nucleotide−mediated signaling
positive regulation of release of sequestered calcium ion into cytosol
positive regulation of calcium ion transport into cytosol
positive regulation of calcium ion transmembrane transport
regulation of release of sequestered calcium ion into cytosol
regulation of calcium ion transmembrane transport
regulation of calcium ion transport into cytosol
antimicrobial humoral immune response mediated by antimicrobial peptide
positive regulation of cation transmembrane transport
positive regulation of ion transmembrane transport
sequestering of calcium ion
release of sequestered calcium ion into cytosol
regulation of sequestering of calcium ion
negative regulation of sequestering of calcium ion
3.00
3.25
3.50
3.75
4.00
Count
GeneRatio
0.375
0.400
0.425
0.450
0.475
0.500
p.adjust
0.04
0.03
GO (BP)
ITLN1
CXCL11
CXCL9
CXCL10
fold change
2
1
0
−1
size
3.00
3.25
3.50
3.75
4.00
GO (MF)
CXCL9
CXCR chemokine receptor binding
CXCL10
CXCL11
size
3
fold change
2.6
2.4
2.2
2.0

**Figure S18.** Severe vs Mild

GSEA Reactome

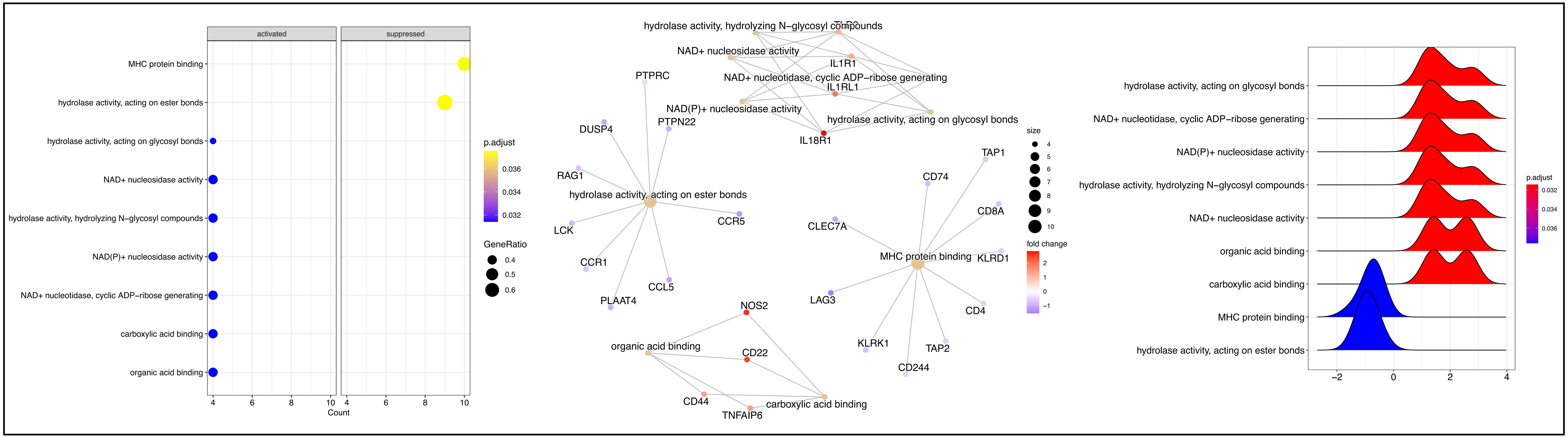


ORA

GO (MF)

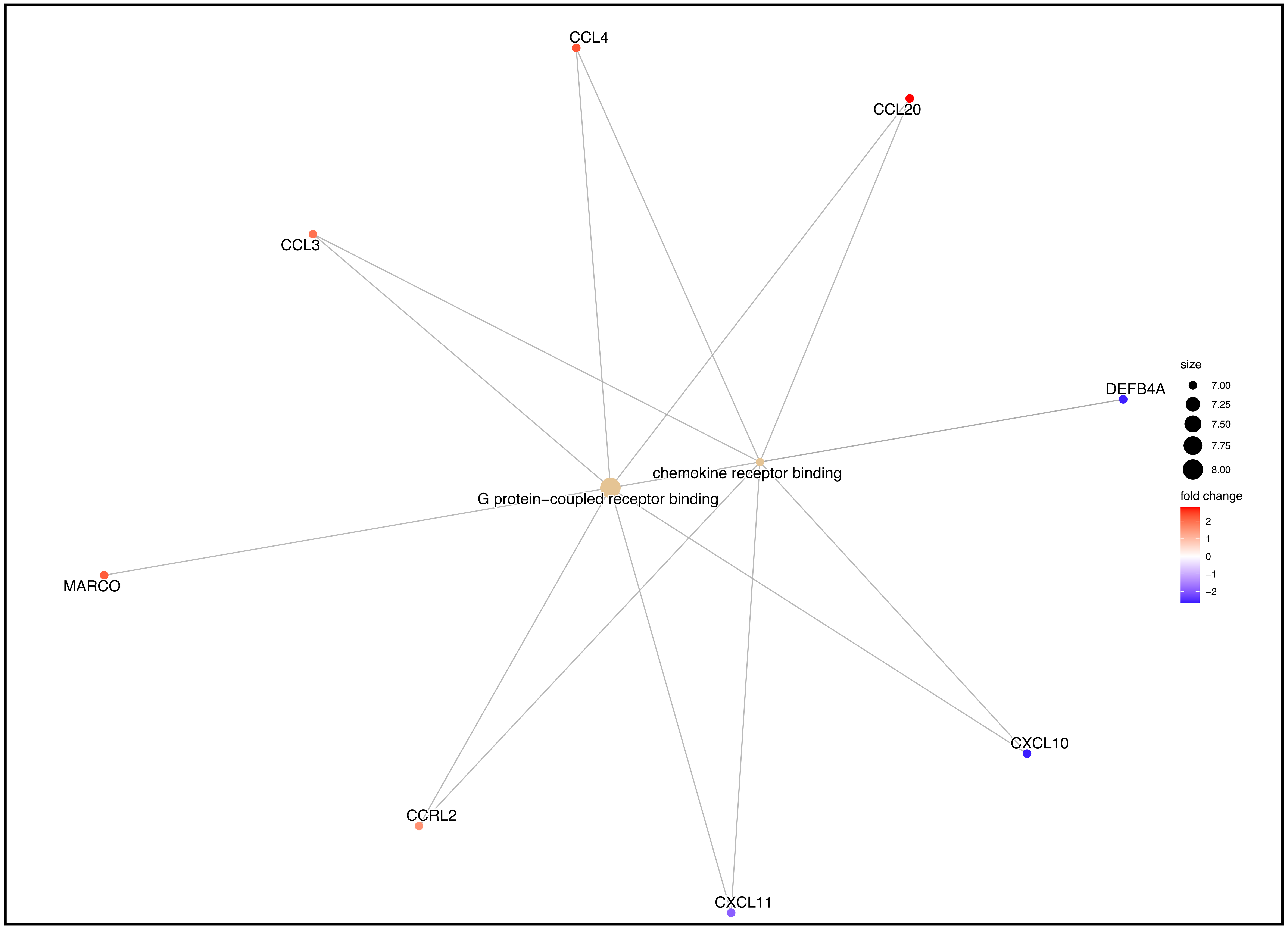

### Figure S19. Severe vs Moderate GSEA

## GO (BP)

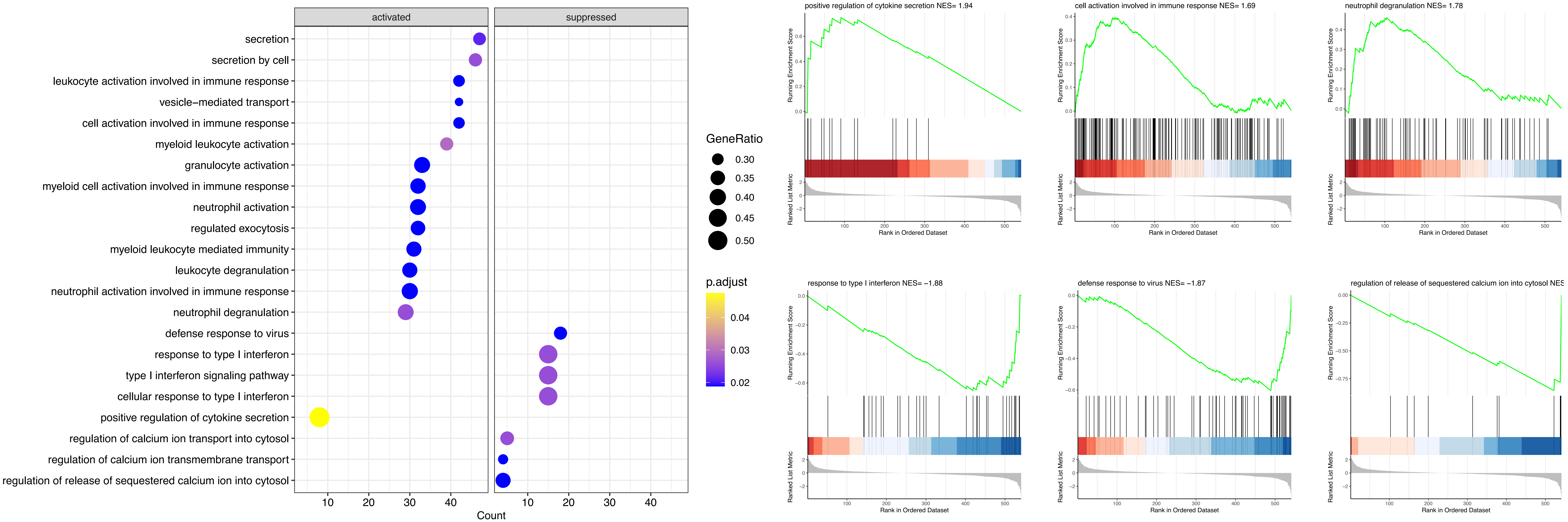


#### Reactome

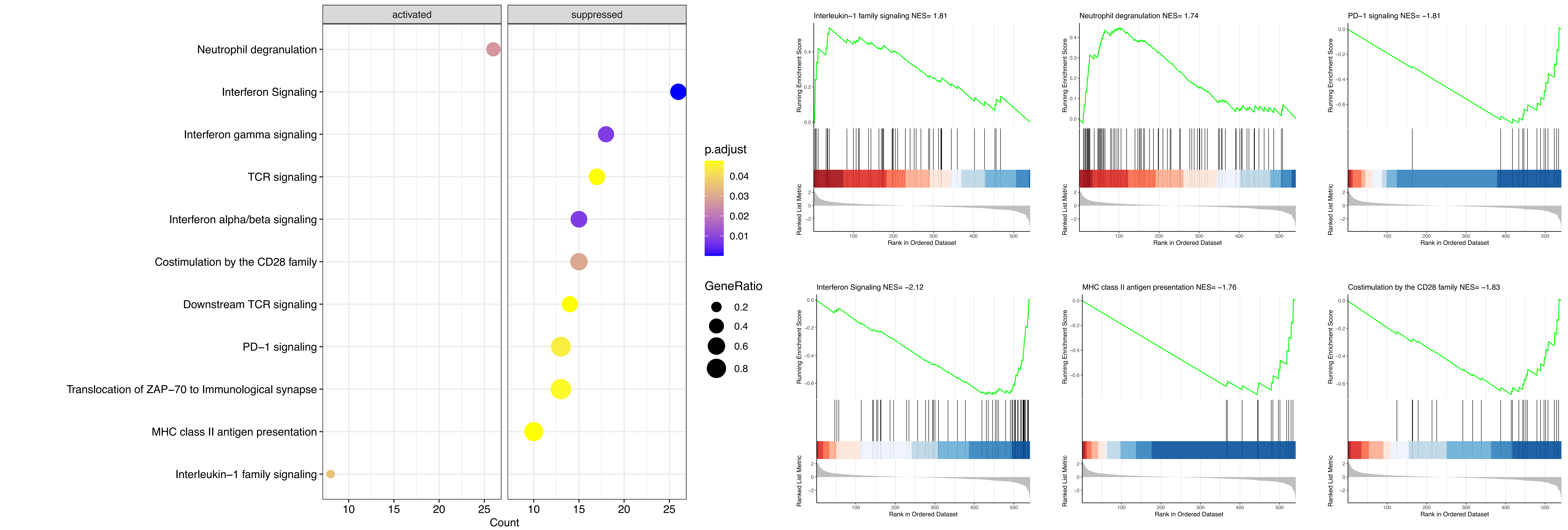

### Figure S20. Moderate vs Mild GSEA

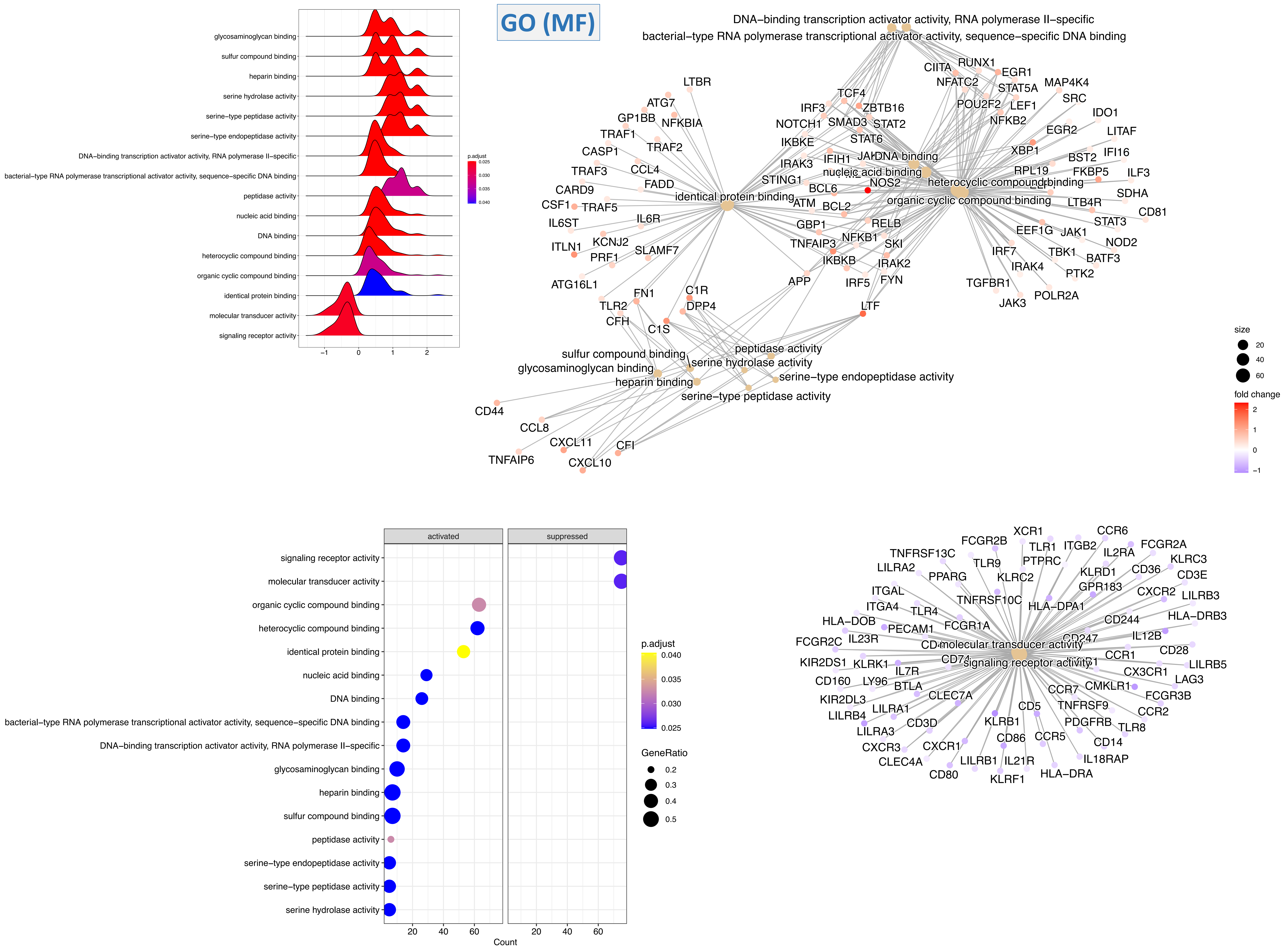

### Figure S21.

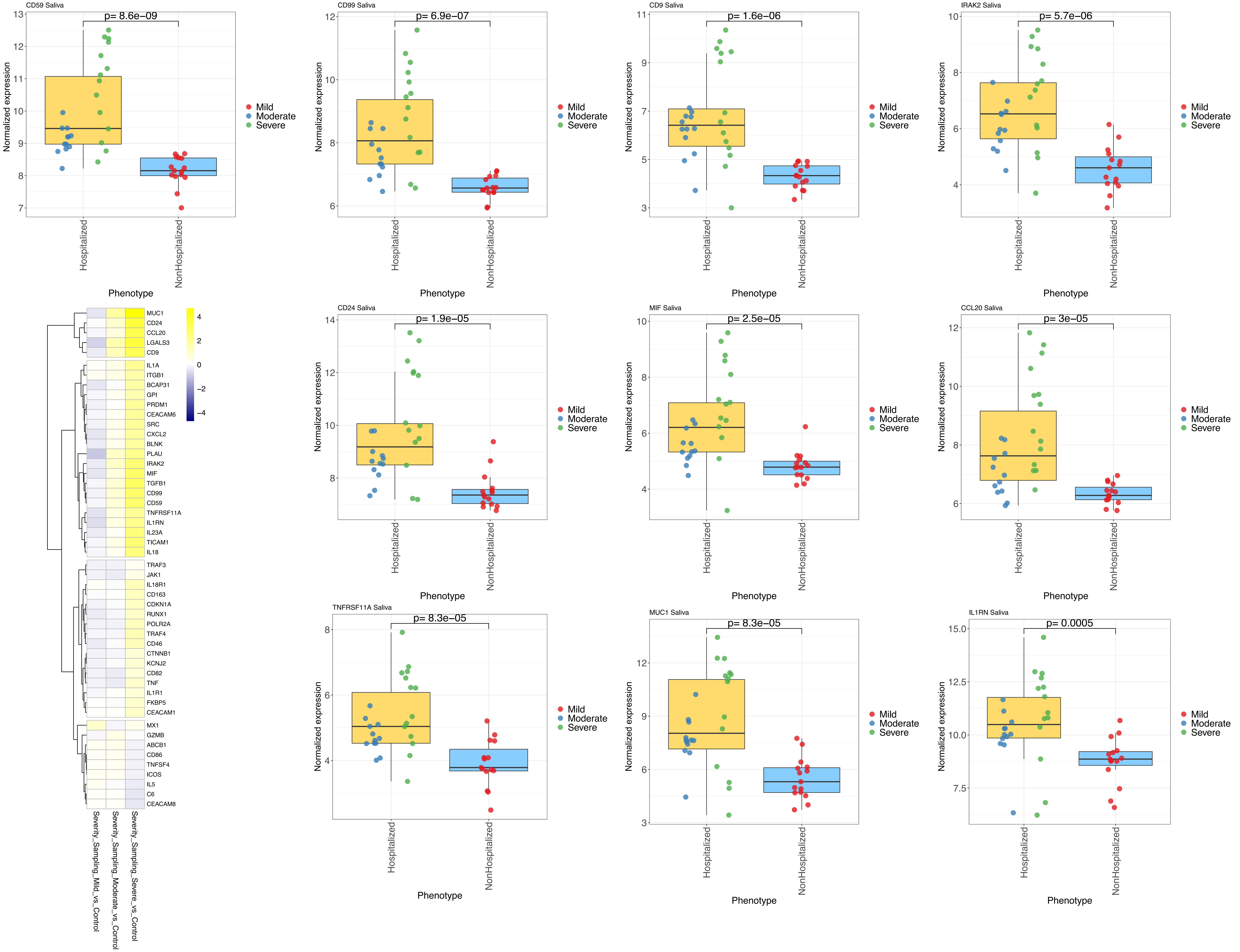
CD59 Saliva
p= 8.6e−09
CD99 Saliva
p= 6.9e−07
CD9 Saliva
p= 1.6e−06
IRAK2 Saliva
p= 5.7e−06
CD24 Saliva
p= 1.9e−05
MIF Saliva
p= 2.5e−05
CCL20 Saliva
p= 3e−05
TNFRSF11A Saliva
p= 8.3e−05
MUC1 Saliva
p= 8.3e−05
IL1RN Saliva
p= 0.0005
Normalized expression
Phenotype
Hospitalized
NonHospitalized
Mild
Moderate
Severe
MUC1
CD24
CCL20
LGALS3
CD9
IL1A
ITGB1
BCAP31
GPI
PRDM1
CEACAM6
SRC
CXCL2
BLNK
PLAU
IRAK2
MIF
TGFB1
CD99
CD59
TNFRSF11A
IL1RN
IL23A
TICAM1
IL18
TRAF3
JAK1
IL18R1
CD163
CDKN1A
RUNX1
POLR2A
TRAF4
CD46
CTNNB1
KCNJ2
CD82
TNF
IL1R1
FKBP5
CEACAM1
MX1
GZMB
ABCB1
CD86
TNFSF4
ICOS
IL5
C6
CEACAM8
Severity_Sampling_Mild_vs_Control
Severity_Sampling_Moderate_vs_Control
Severity_Sampling_Severe_vs_Control

### Figure S22.

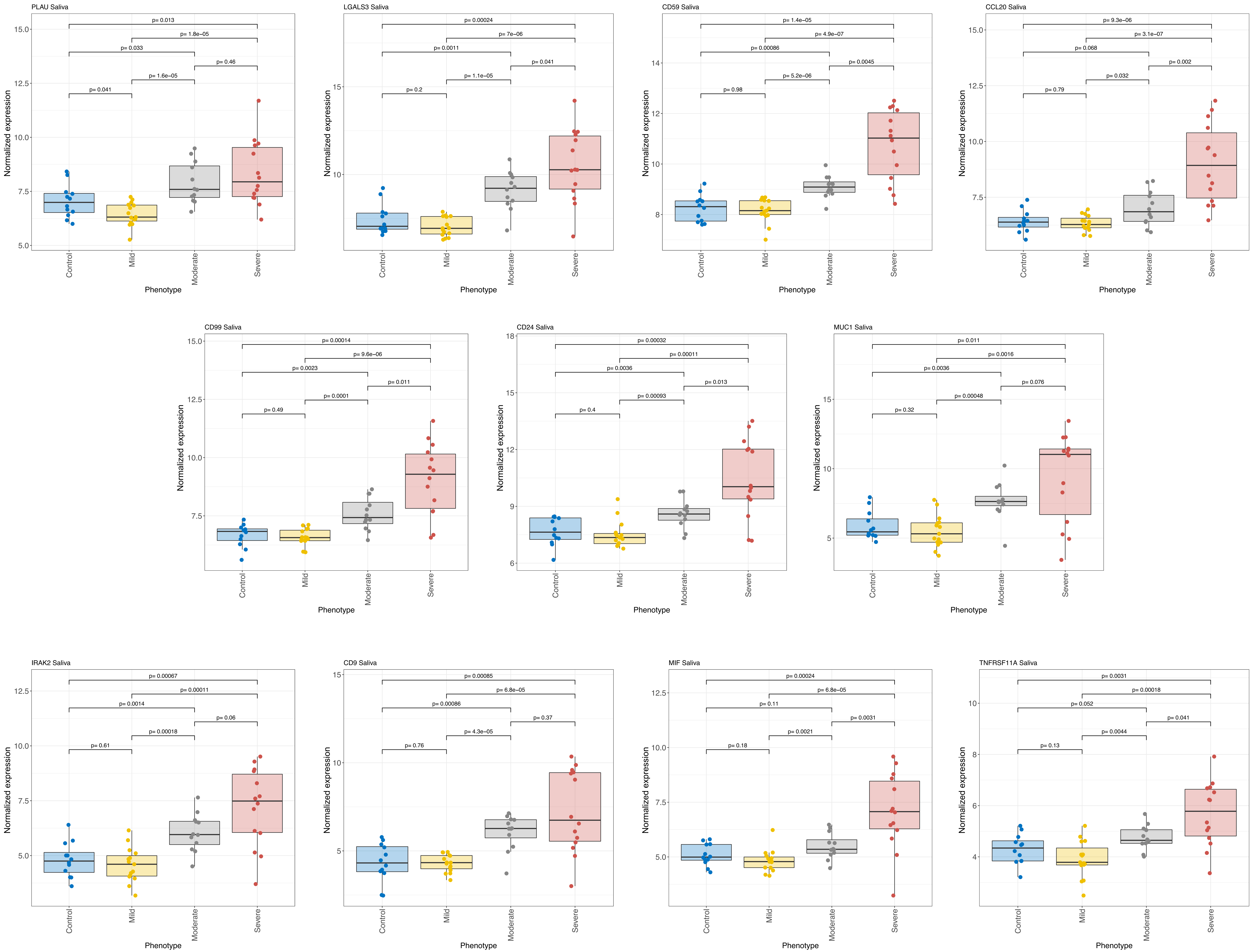

PLAU Saliva
p= 0.013
p= 1.8e−05
p= 0.033
p= 0.46
p= 1.6e−05
p= 0.041
LGALS3 Saliva
p= 0.00024
p= 7e−06
p= 0.0011
p= 0.041
p= 1.1e−05
p= 0.2
CD59 Saliva
p= 1.4e−05
p= 4.9e−07
p= 0.00086
p= 0.0045
p= 5.2e−06
p= 0.98
CCL20 Saliva
p= 9.3e−06
p= 3.1e−07
p= 0.068
p= 0.002
p= 0.032
p= 0.79
CD99 Saliva
p= 0.00014
p= 9.6e−06
p= 0.0023
p= 0.011
p= 0.0001
p= 0.49
CD24 Saliva
p= 0.00032
p= 0.00011
p= 0.0036
p= 0.013
p= 0.00093
p= 0.4
MUC1 Saliva
p= 0.011
p= 0.0016
p= 0.0036
p= 0.076
p= 0.00048
p= 0.32
IRAK2 Saliva
p= 0.00067
p= 0.00011
p= 0.0014
p= 0.06
p= 0.00018
p= 0.61
CD9 Saliva
p= 0.00085
p= 6.8e−05
p= 0.00086
p= 0.37
p= 4.3e−05
p= 0.76
MIF Saliva
p= 0.00024
p= 6.8e−05
p= 0.11
p= 0.0031
p= 0.0021
p= 0.18
TNFRSF11A Saliva
p= 0.0031
p= 0.00018
p= 0.052
p= 0.041
p= 0.0044
p= 0.13
Normalized expression
Control
Mild
Moderate
Severe
Phenotype

### Figure S23.

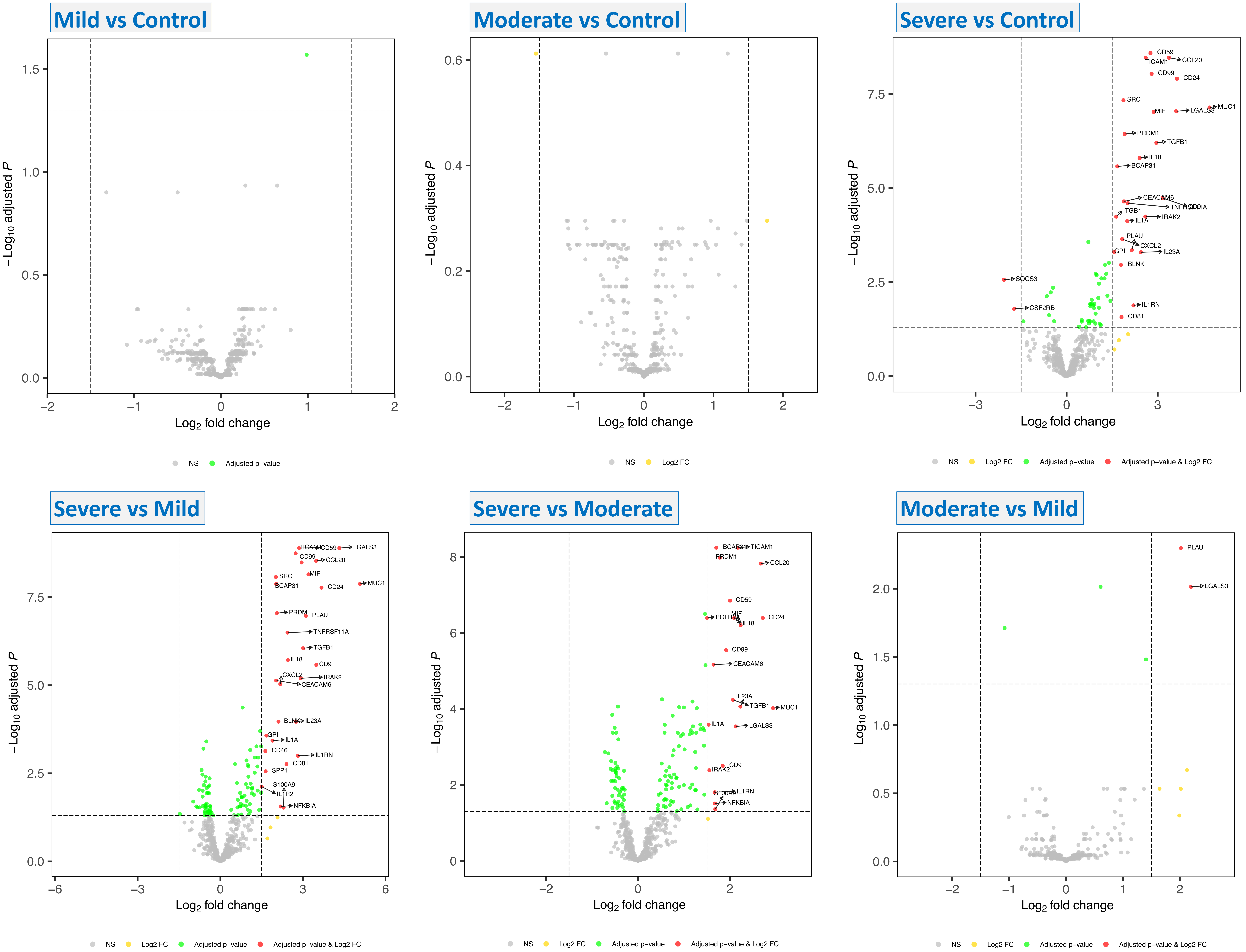
Mild vs Control
Moderate vs Control
Severe vs Control
Severe vs Mild
Severe vs Moderate
Moderate vs Mild
−Log10 adjusted P
Log2 fold change
NS
Log2 FC
Adjusted p−value
Adjusted p−value & Log2 FC
CD59
TICAM1
CCL20
CD99
CD24
SRC
MIF
LGALS3
MUC1
PRDM1
TGFB1
IL18
BCAP31
CEACAM6
TNFRSF11A
CD9
ITGB1
IL1A
IRAK2
PLAU
CXCL2
GPI
IL23A
BLNK
SOCS3
CSF2RB
IL1RN
CD81
CD46
SPP1
S100A9
IL1R2
NFKBIA
POLR2A

### Figure S24.

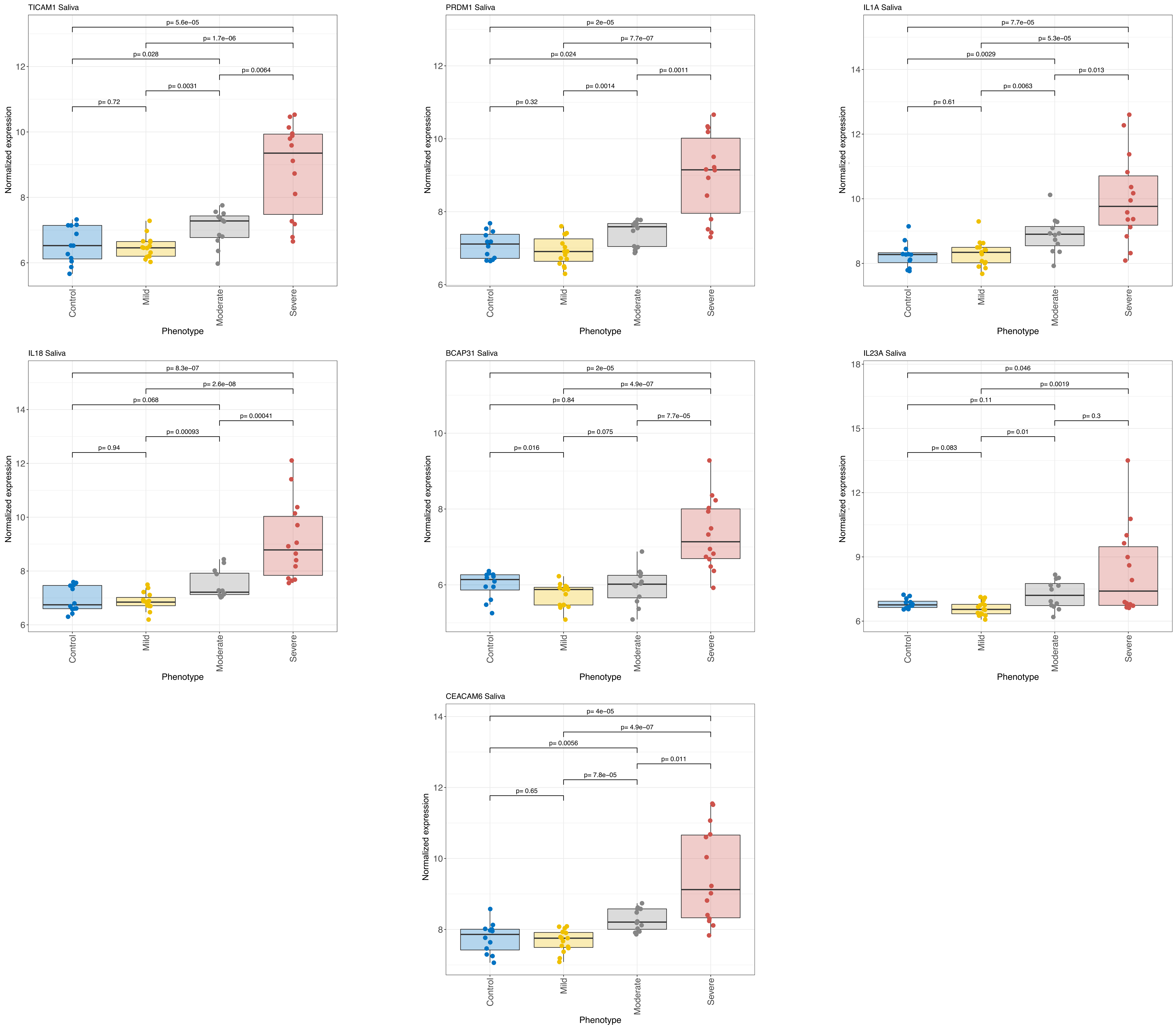

TICAM1 Saliva
p= 5.6e−05
p= 1.7e−06
p= 0.028
p= 0.0064
p= 0.0031
p= 0.72
Normalized expression
6
8
10
12
Control
Mild
Moderate
Severe
Phenotype
PRDM1 Saliva
p= 2e−05
p= 7.7e−07
p= 0.024
p= 0.0011
p= 0.0014
p= 0.32
Normalized expression
6
8
10
12
Control
Mild
Moderate
Severe
Phenotype
IL1A Saliva
p= 7.7e−05
p= 5.3e−05
p= 0.0029
p= 0.013
p= 0.0063
p= 0.61
Normalized expression
8
10
12
14
Control
Mild
Moderate
Severe
Phenotype
IL18 Saliva
p= 8.3e−07
p= 2.6e−08
p= 0.068
p= 0.00041
p= 0.00093
p= 0.94
Normalized expression
6
8
10
12
14
Control
Mild
Moderate
Severe
Phenotype
BCAP31 Saliva
p= 2e−05
p= 4.9e−07
p= 0.84
p= 7.7e−05
p= 0.075
p= 0.016
Normalized expression
6
8
10
Control
Mild
Moderate
Severe
Phenotype
IL23A Saliva
p= 0.046
p= 0.0019
p= 0.11
p= 0.3
p= 0.01
p= 0.083
Normalized expression
6
9
12
15
18
Control
Mild
Moderate
Severe
Phenotype
CEACAM6 Saliva
p= 4e−05
p= 4.9e−07
p= 0.0056
p= 0.011
p= 7.8e−05
p= 0.65
Normalized expression
8
10
12
14
Control
Mild
Moderate
Severe
Phenotype

### Figure S25.

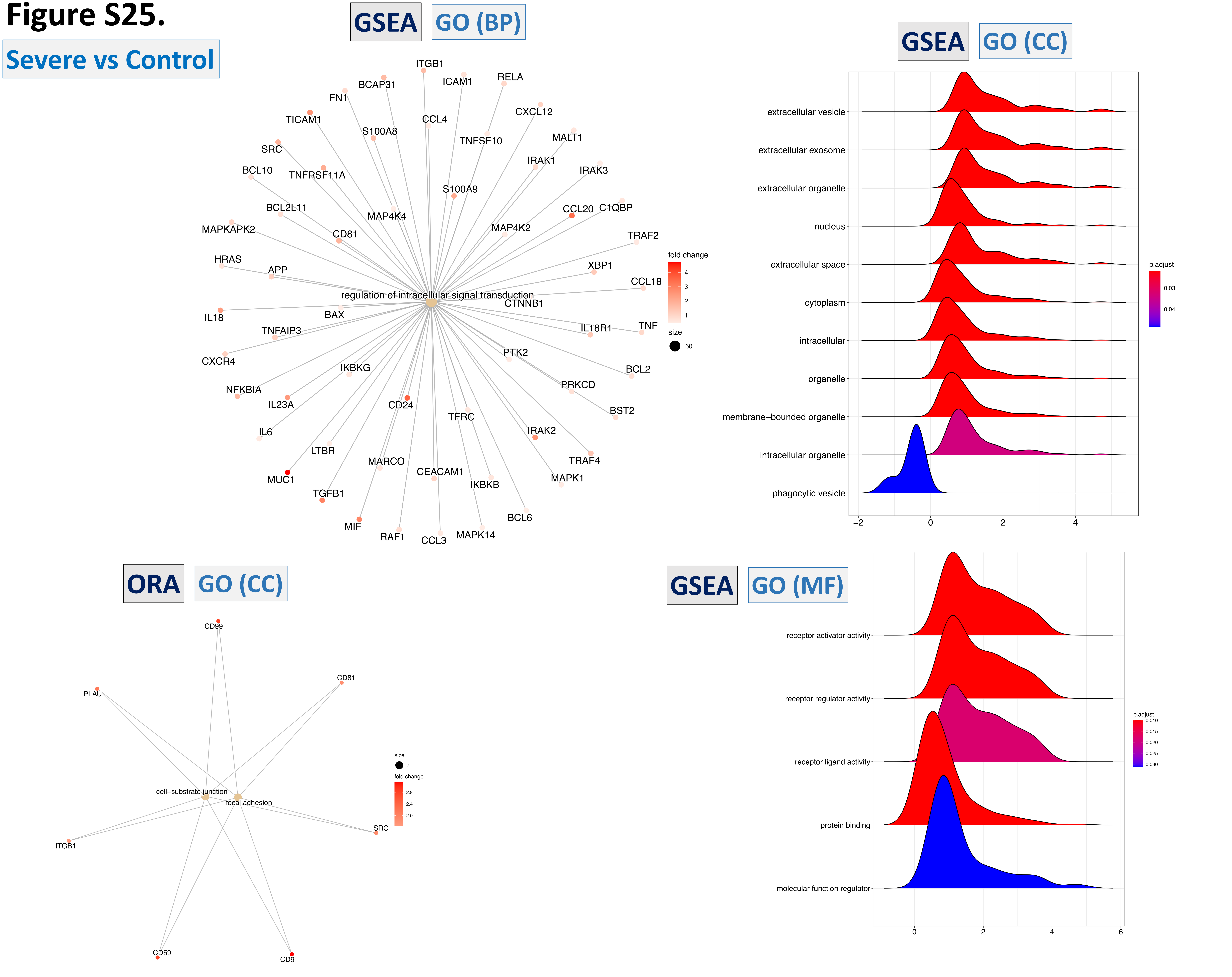

Severe vs Control
GSEA
GO (BP)
regulation of intracellular signal transduction
ITGB1
BCAP31
ICAM1
RELA
FN1
CCL4
CXCL12
TICAM1
S100A8
TNFSF10
MALT1
SRC
IRAK1
IRAK3
BCL10
TNFRSF11A
S100A9
BCL2L11
MAP4K4
CCL20
C1QBP
MAPKAPK2
CD81
MAP4K2
TRAF2
HRAS
APP
XBP1
CCL18
CTNNB1
IL18
BAX
IL18R1
TNF
TNFAIP3
CXCR4
PTK2
IKBKG
BCL2
NFKBIA
PRKCD
IL23A
CD24
BST2
TFRC
IL6
IRAK2
LTBR
MARCO
TRAF4
MUC1
CEACAM1
MAPK1
TGFB1
IKBKB
BCL6
MIF
RAF1
CCL3
MAPK14
fold change
4
3
2
1
size
60
GSEA
GO (CC)
extracellular vesicle
extracellular exosome
extracellular organelle
nucleus
extracellular space
cytoplasm
intracellular
organelle
membrane−bounded organelle
intracellular organelle
phagocytic vesicle
−2
0
2
4
p.adjust
0.03
0.04
ORA
GO (CC)
CD99
PLAU
CD81
cell−substrate junction
focal adhesion
SRC
ITGB1
CD59
CD9
size
7
fold change
2.8
2.4
2.0
GSEA
GO (MF)
receptor activator activity
receptor regulator activity
receptor ligand activity
protein binding
molecular function regulator
0
2
4
6
p.adjust
0.010
0.015
0.020
0.025
0.030

### Figure S26. Severe vs Mild

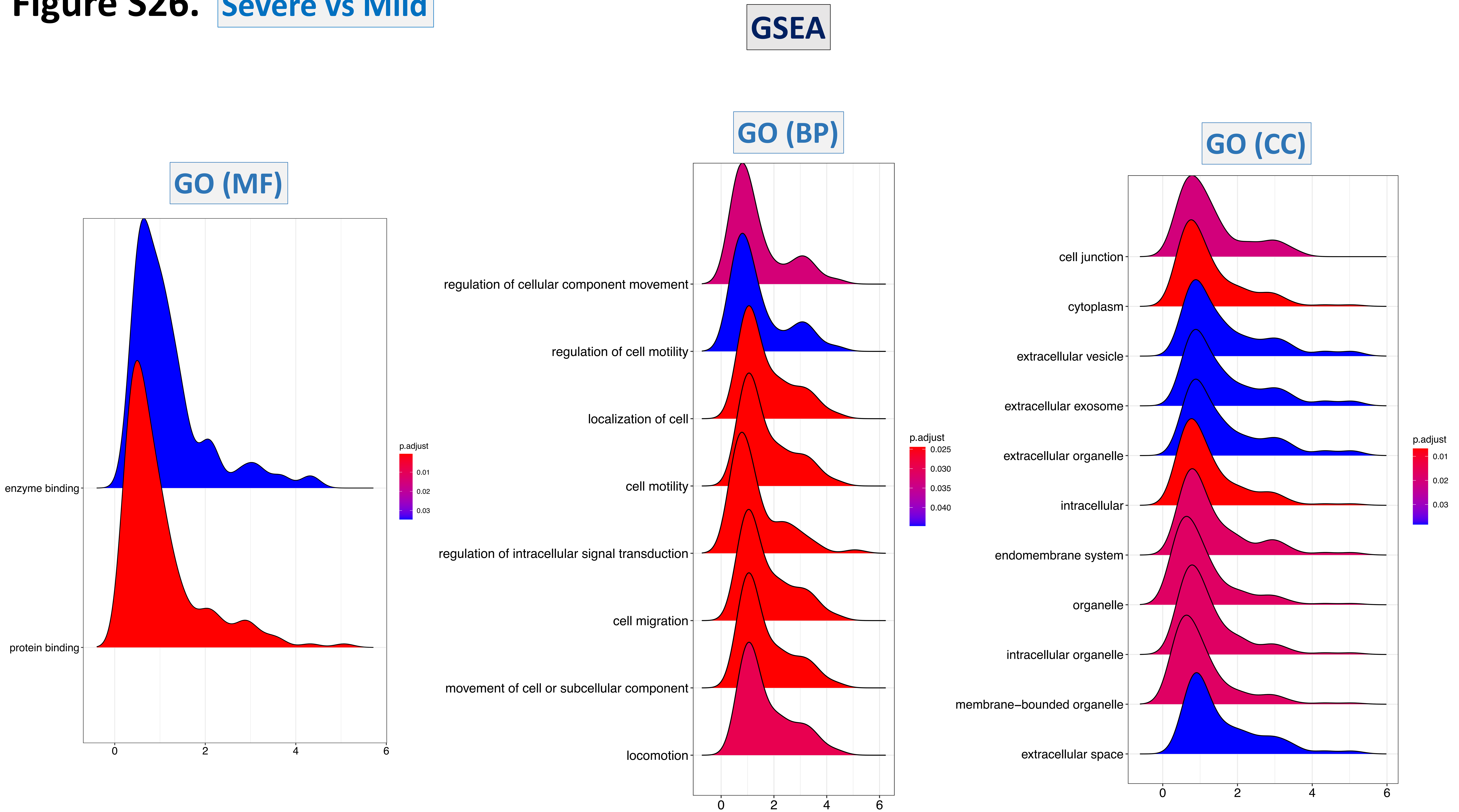

### Figure S27. Severe vs Moderate

GSEA

GO (BP)

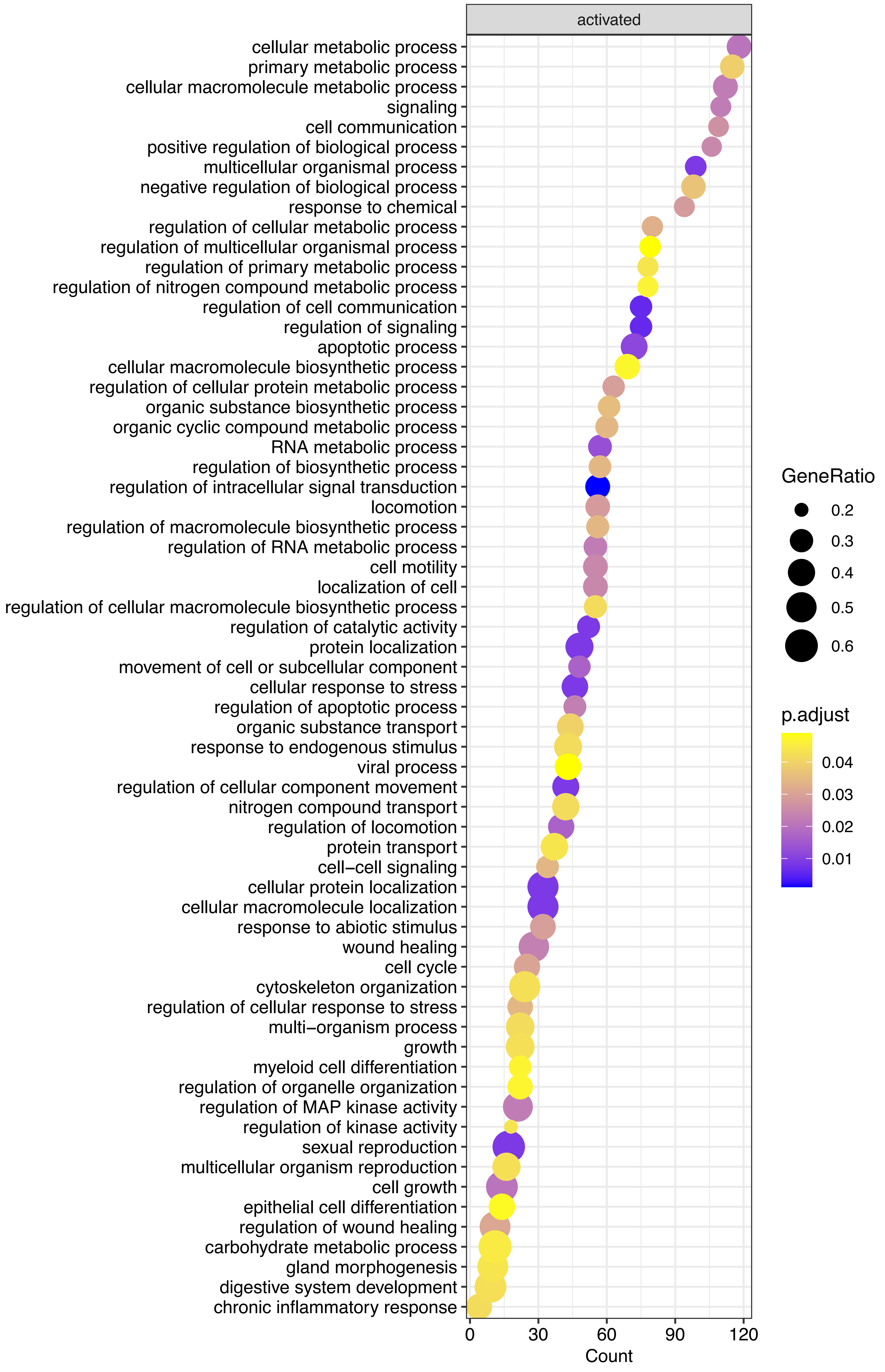


GO (MF)

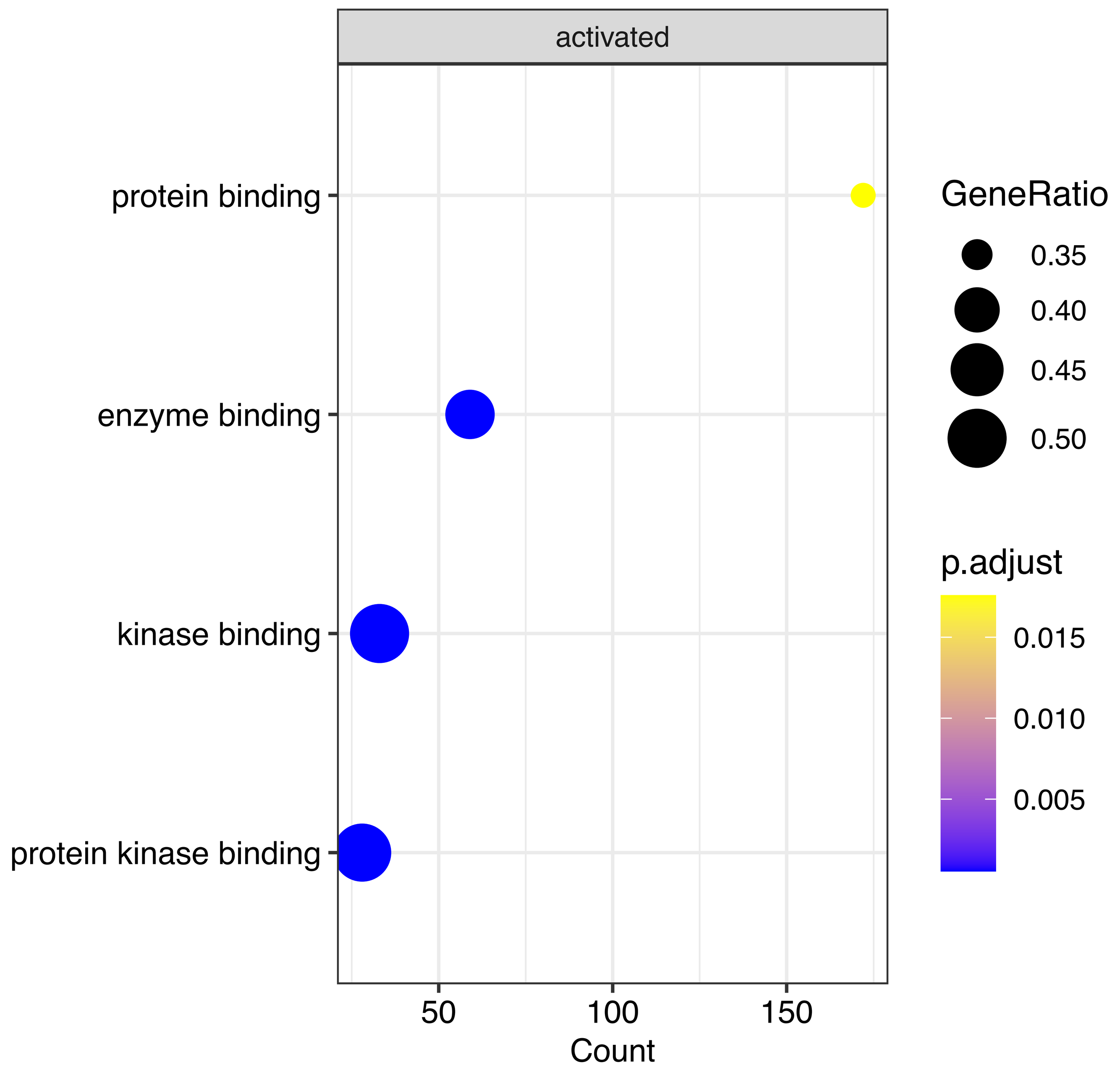


GO (CC)

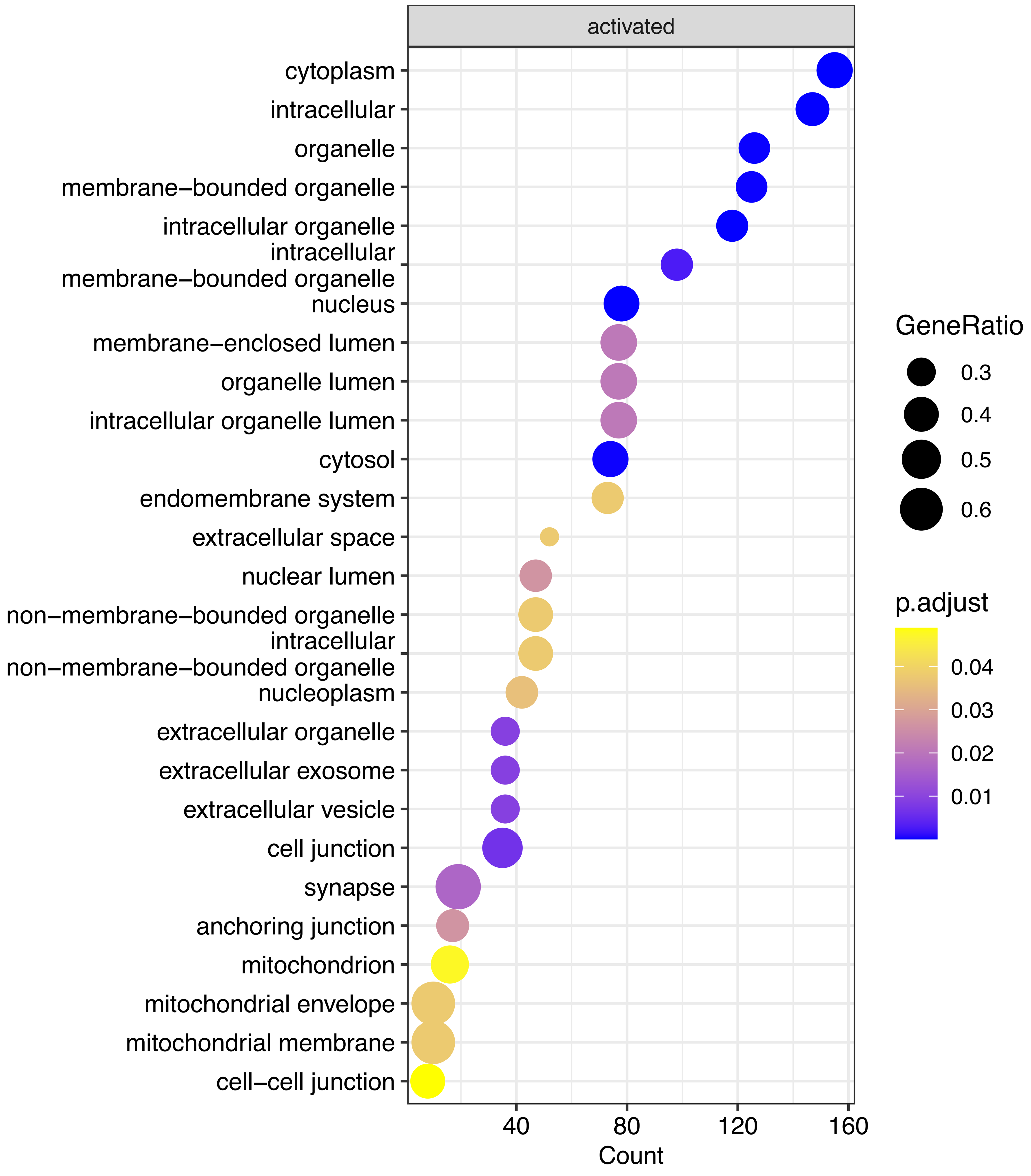

### Figure S28. Moderate vs Mild GSEA

GO (MF)

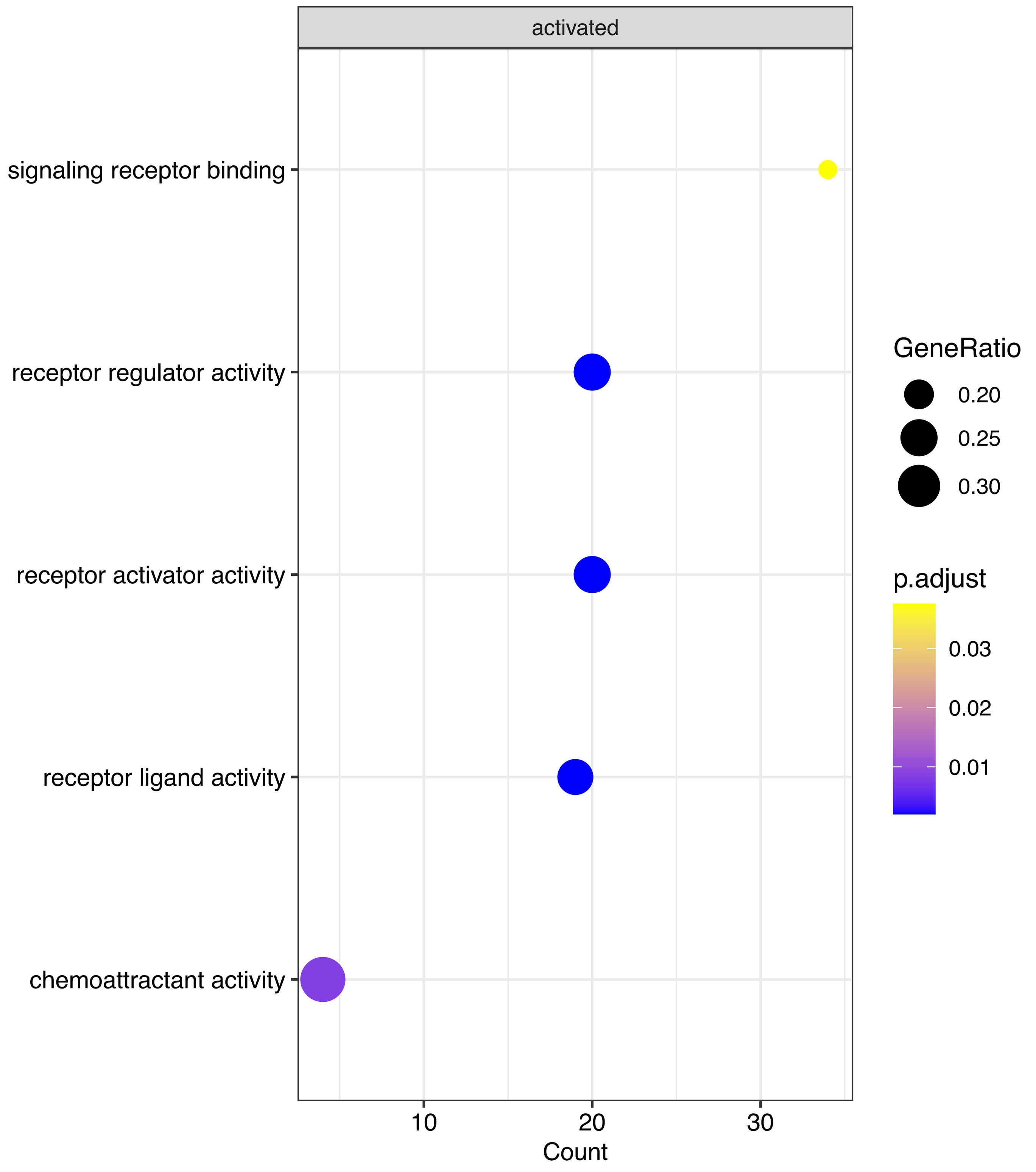


Reactome

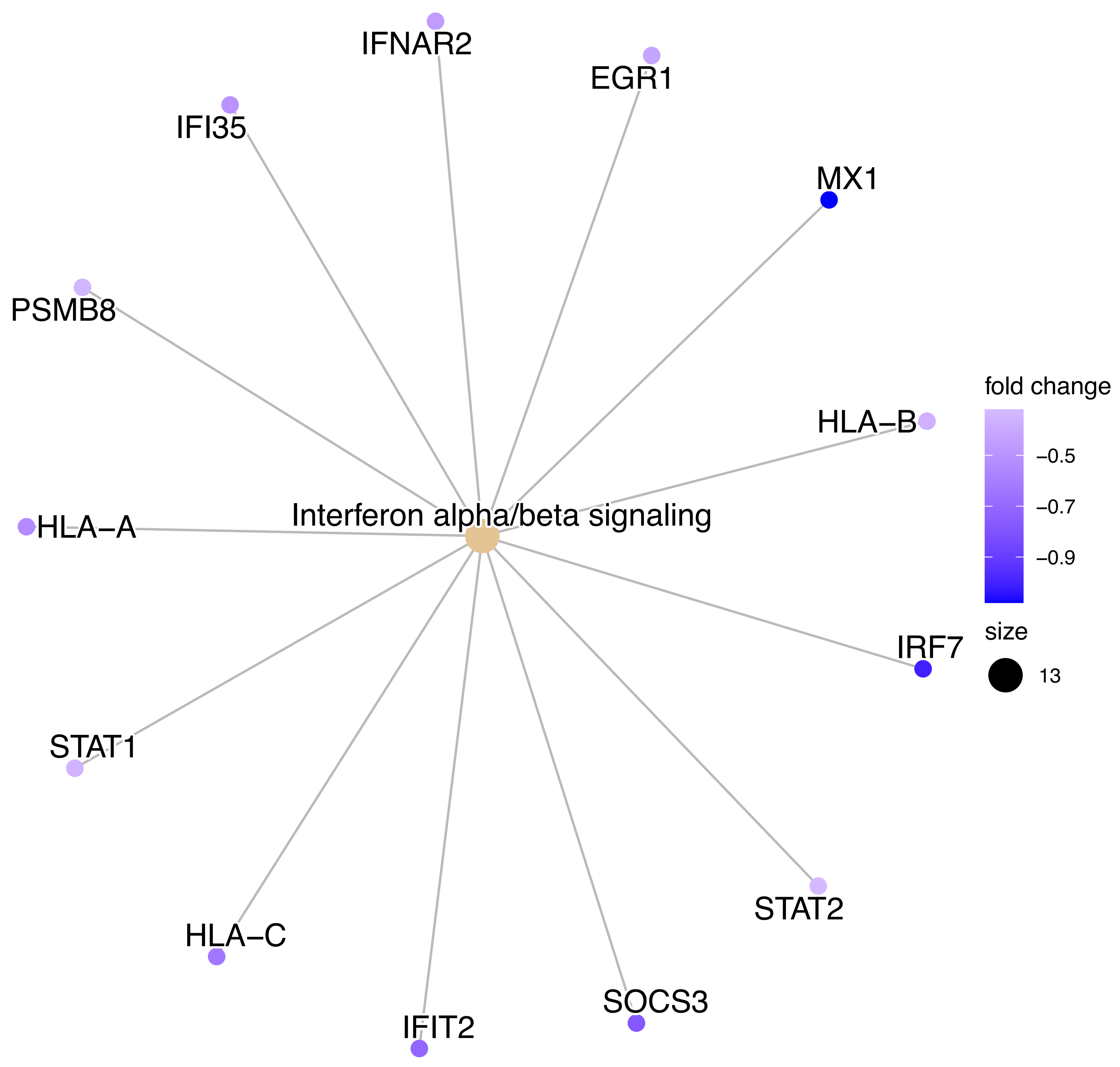


GO (BP)

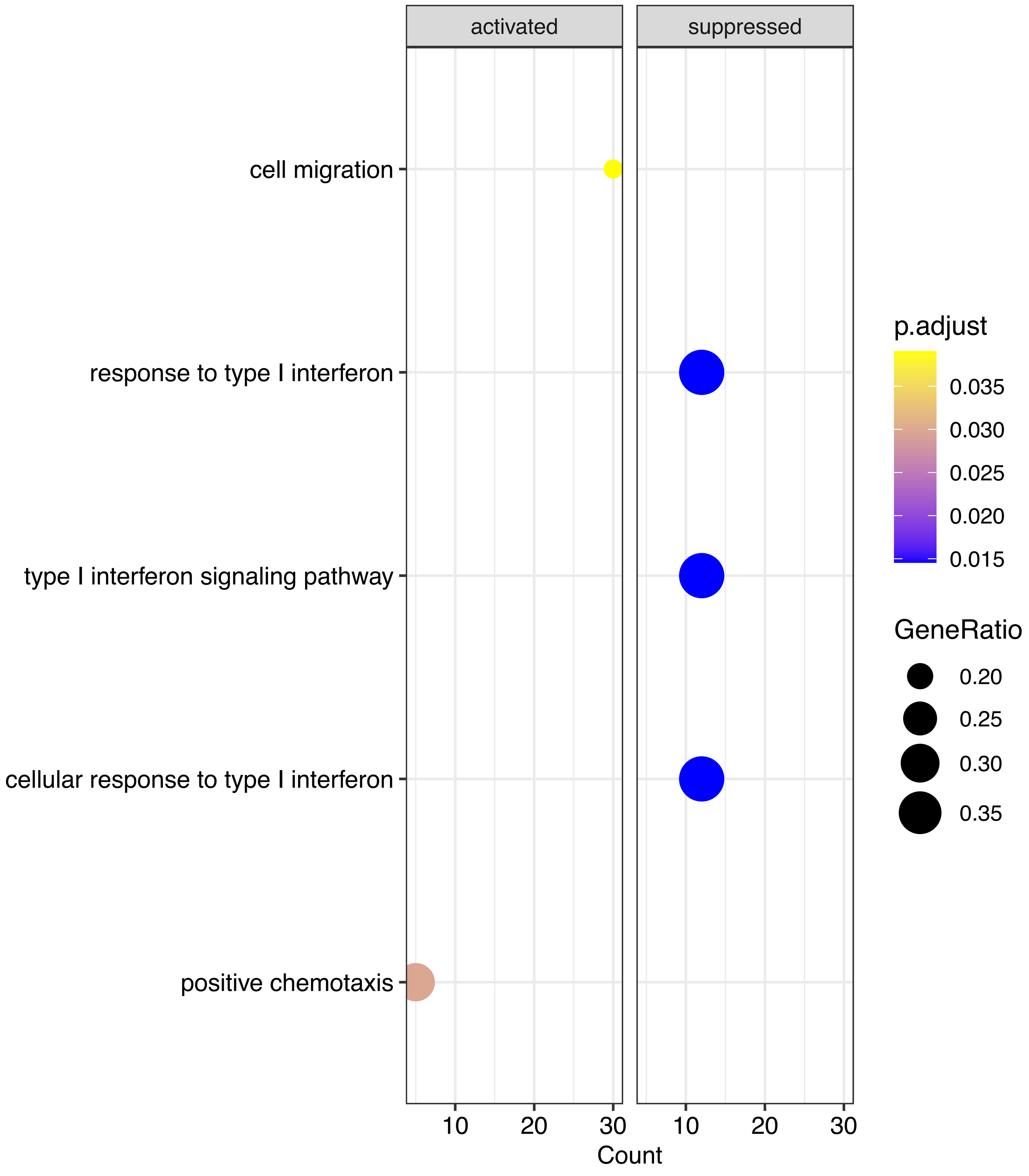

**Figure S29.** Moderate vs Control

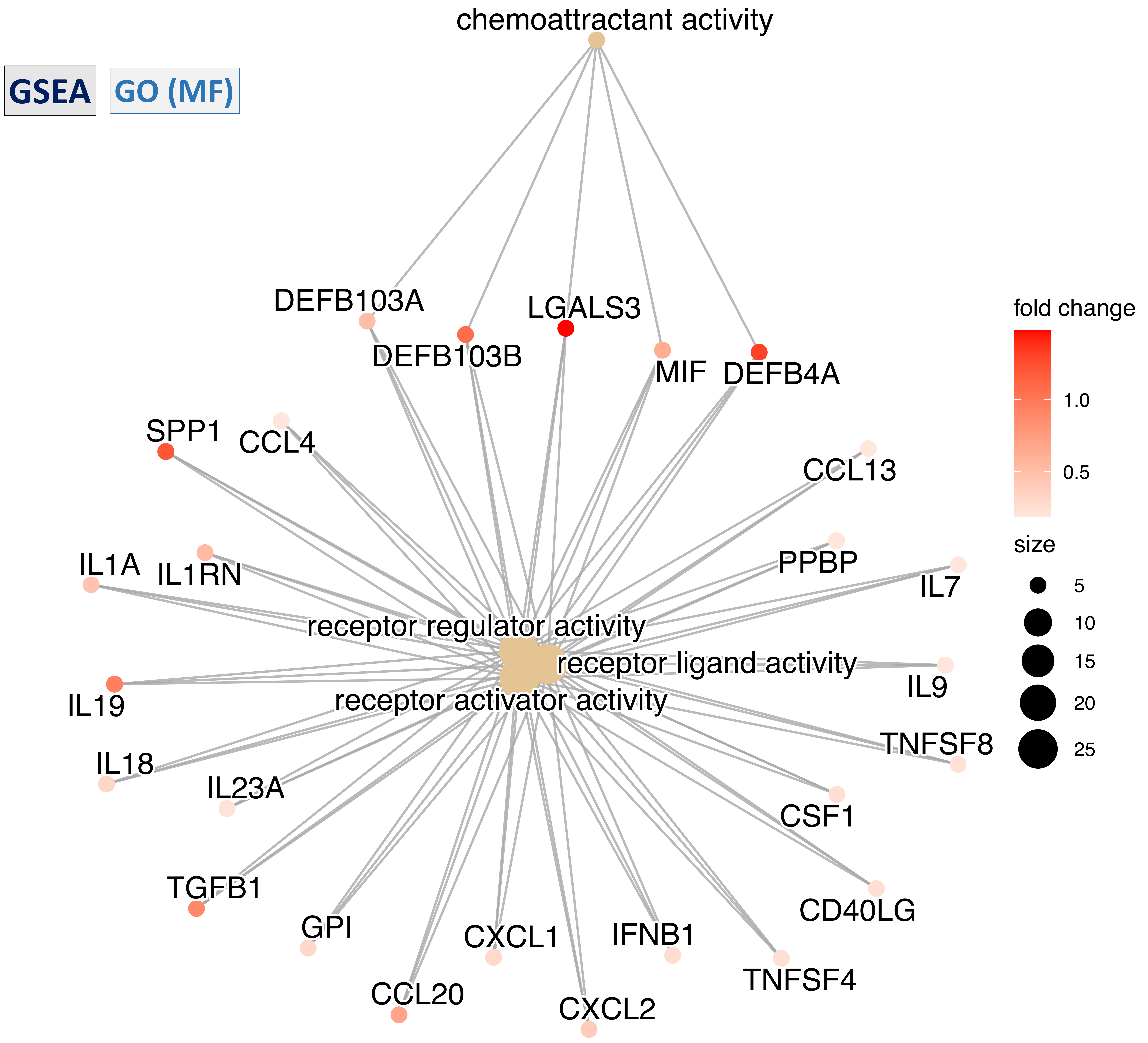

### Figure S30. Mild vs Control

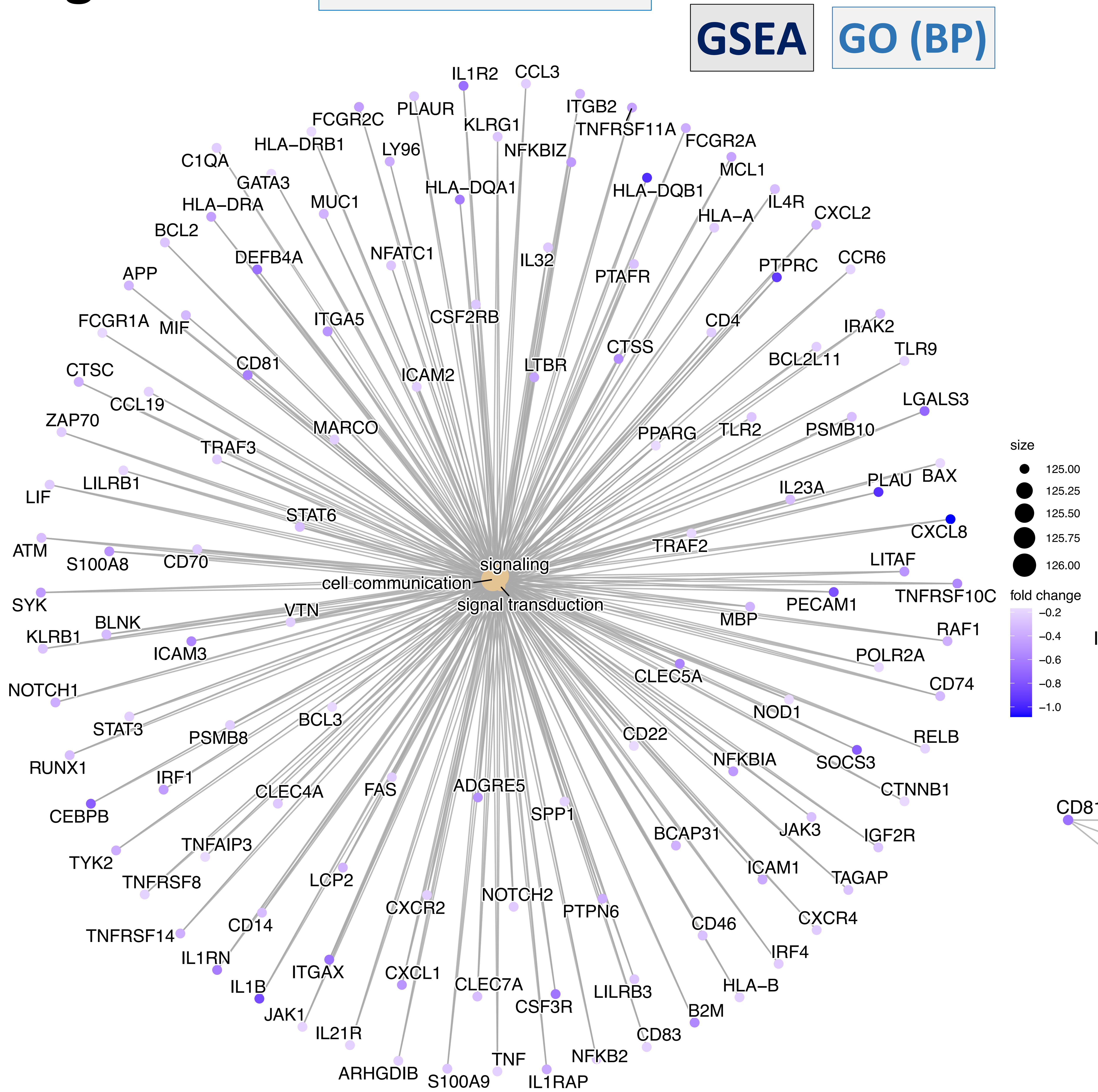


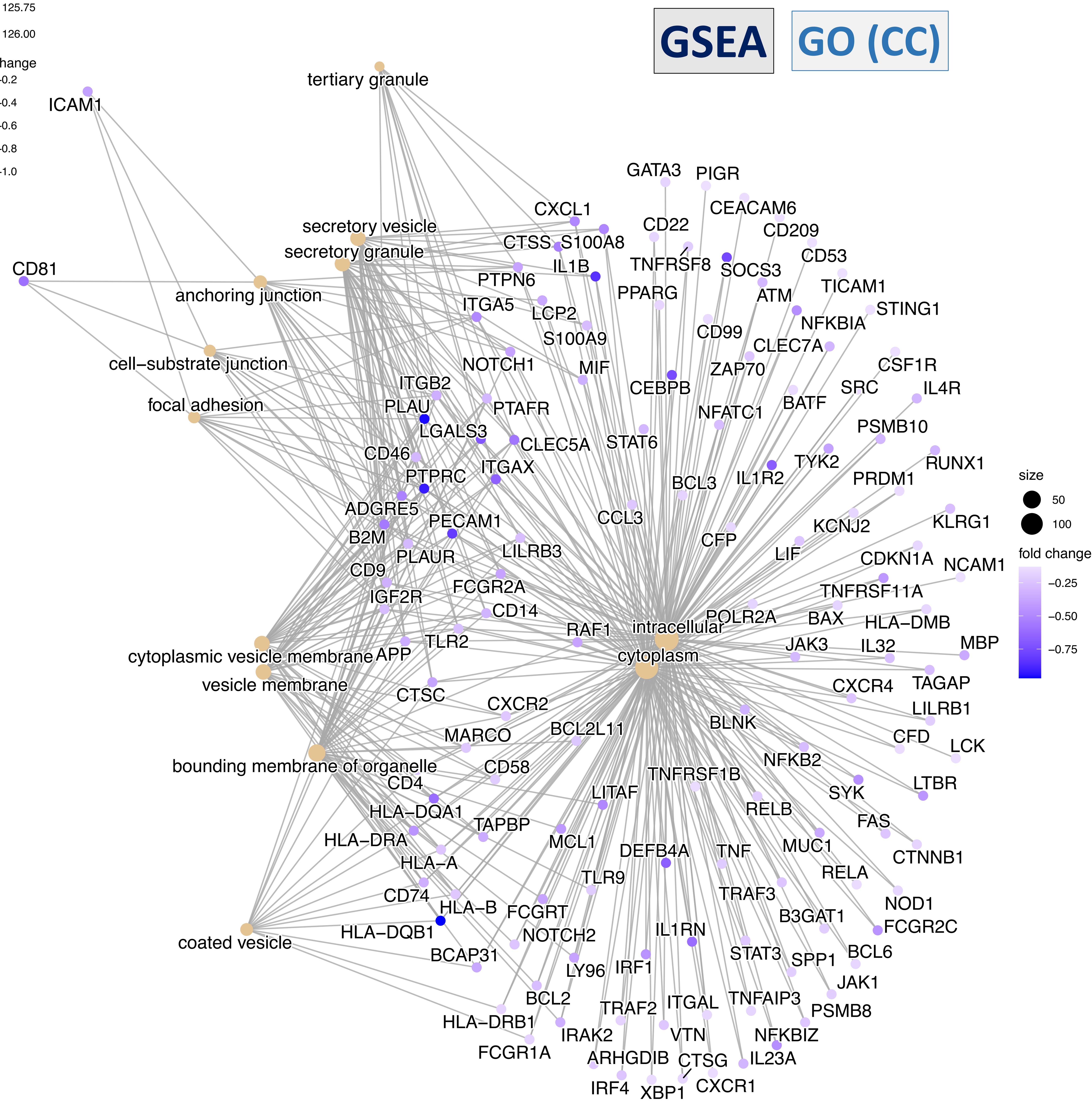

### Figure S31.

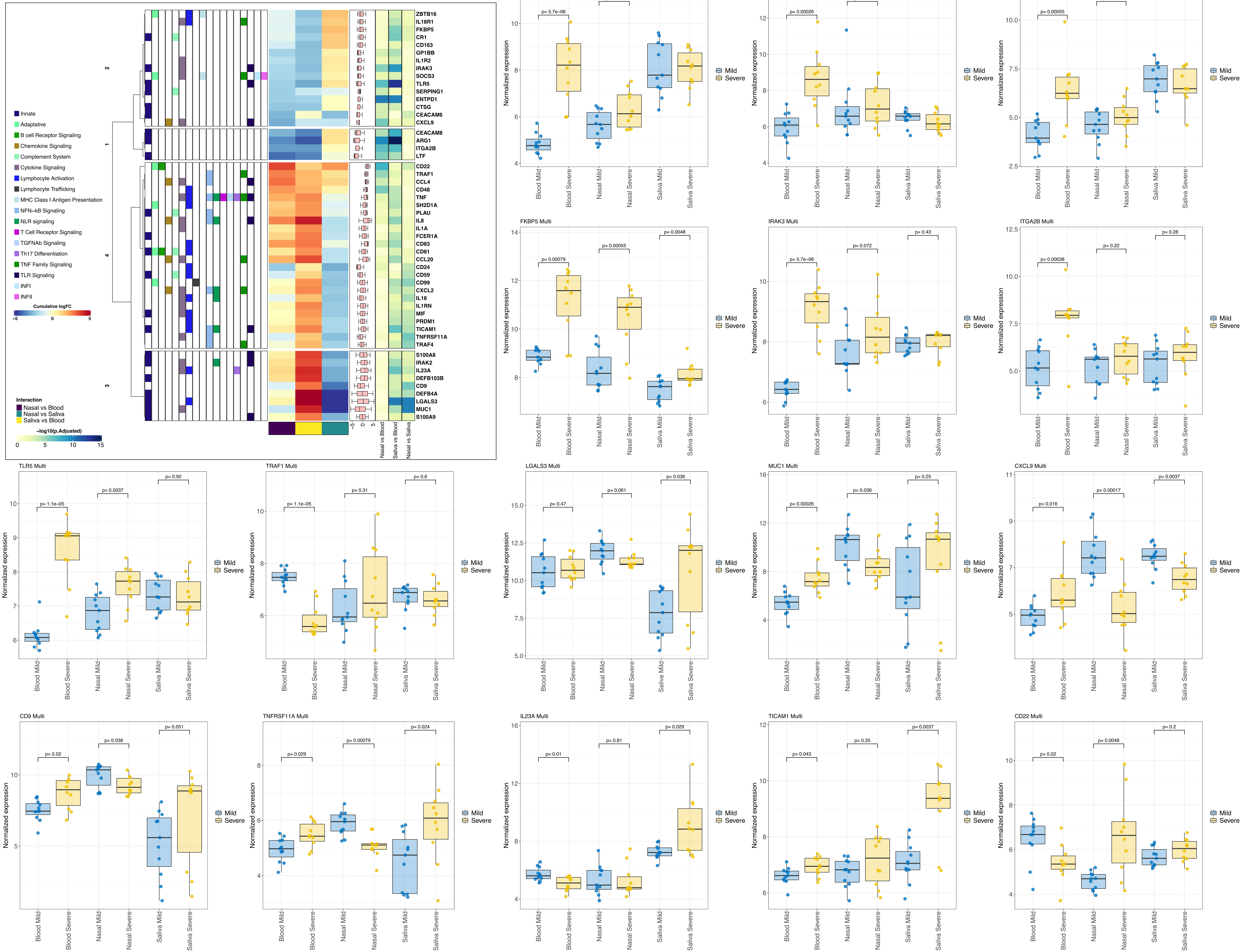

ZBTB16
IL18R1
FKBP5
CR1
CD163
GP1BB
IL1R2
IRAK3
SOCS3
TLR5
SERPING1
ENTPD1
CTSG
CEACAM6
CXCL9
CEACAM8
ARG1
ITGA2B
LTF
CD22
TRAF1
CCL4
CD48
TNF
SH2D1A
PLAU
IL8
IL1A
FCER1A
CD83
CD81
CCL20
CD24
CD59
CD99
CXCL2
IL18
IL1RN
MIF
PRDM1
TICAM1
TNFRSF11A
TRAF4
S100A8
IRAK2
IL23A
DEFB103B
CD9
DEFB4A
LGALS3
MUC1
S100A9
Nasal vs Blood
Saliva vs Blood
Nasal vs Saliva
Innate
Adaptative
B cell Receptor Signaling
Chemokine Signaling
Complement System
Cytokine Signaling
Lymphocyte Activation
Lymphocyte Trafficking
MHC Class I Antigen Presentation
NFN-kB Signaling
NLR signaling
T Cell Receptor Signaling
TGFNAb Signaling
Th17 Differentiation
TNF Family Signaling
TLR Signaling
INFI
INFII
Cumulative logFC
Interaction
Nasal vs Blood
Nasal vs Saliva
Saliva vs Blood
-log10(p.Adjusted)
ARG1 Multi
p= 5.7e-06
p= 0.11
p= 0.81
LTF Multi
p= 0.00026
p= 0.51
p= 0.43
CEACAM8 Multi
p= 0.00055
p= 0.35
p= 0.56
FKBP5 Multi
p= 0.00079
p= 0.00055
p= 0.0048
IRAK3 Multi
p= 5.7e-06
p= 0.072
p= 0.43
ITGA2B Multi
p= 0.00038
p= 0.22
p= 0.28
TLR5 Multi
p= 1.1e-05
p= 0.0037
p= 0.92
TRAF1 Multi
p= 1.1e-05
p= 0.31
p= 0.6
LGALS3 Multi
p= 0.47
p= 0.061
p= 0.036
MUC1 Multi
p= 0.00026
p= 0.036
p= 0.25
CXCL9 Multi
p= 0.016
p= 0.00017
p= 0.0037
CD9 Multi
p= 0.02
p= 0.036
p= 0.051
TNFRSF11A Multi
p= 0.029
p= 0.00079
p= 0.024
IL23A Multi
p= 0.01
p= 0.81
p= 0.029
TICAM1 Multi
p= 0.043
p= 0.35
p= 0.0037
CD22 Multi
p= 0.02
p= 0.0048
p= 0.2
Normalized expression
Blood Mild
Blood Severe
Nasal Mild
Nasal Severe
Saliva Mild
Saliva Severe
Mild
Severe
